# Genetically encoded assembly recorder temporally resolves cellular histories *in cellulo* and *in vivo*

**DOI:** 10.1101/2025.07.16.664392

**Authors:** Yuqing Yan, Jiaxi Lu, Zhe Li, Zuohan Zhao, Timothy F. Shay, Shunzhi Wang, Yaping Lei, Yimei Wang, Wei Chen, Patrick Parker, Hongru Yang, Aileen Qi, Yongzhi Sun, Dwight E. Bergles, David Baker, Dingchang Lin

## Abstract

Mapping cellular activity with high spatiotemporal precision in complex tissues is essential for understanding organ physiology, pathology, and regenerative processes. Here, we introduce **<u>G</u>**ranularly **<u>E</u>**xpanding **<u>M</u>**emory for **I**ntracellular **<u>N</u>**arrative **I**ntegration (GEMINI), an *in cellulo* recording platform that leverages a computationally designed protein assembly as an intracellular memory device to record individual cells’ activity histories. GEMINI grows predictably within live cells with minimal interference to cellular functions, capturing cellular activities as tree-ring-like fluorescent patterns in the expanding scaffolds for imaging-based retrospective readout. Absolute chronological information of activity histories was attainable with hour-level accuracy through the integration of fiducial timestamps. GEMINI effectively resolved differential NFκB-mediated transcriptional changes, distinguishing fast dynamics of 15 minutes, and providing quantifiable signal amplitudes. In a xenograft model, GEMINI recorded inflammation-induced signaling dynamics across tissue with cellular resolution, revealing spatial heterogeneity linked to vascular density. When expressed in the mouse brain, GEMINI exhibited negligible impact on neuronal survival, with animals maintaining normal motor and cognitive behaviors. In physiological contexts, GEMINI successfully resolved both transcriptional changes and activity patterns of neurons in the brain. Together, GEMINI provides a robust and generalizable means for spatiotemporal mapping of cell dynamics underlying physiological and pathological processes in both culture and intact tissues.

## Introduction

Cells constantly change their molecular states in response to internal and external cues^1^. Spatiotemporal mapping of these intricate processes at the single-cell level and throughout an intact organ or organism has been a longstanding challenge in biology. Current cell sensing and profiling modalities primarily rely on either endpoint analysis, which takes static snapshots of a large cell population^2^, or real-time sensing, which temporally maps a subset of cells within a limited volume of a given tissue^3,4^. These approaches have practical limitations that prevent a clear path toward organ-wide spatiotemporal mapping at the cellular level.

A strategy toward this goal is to write cellular histories within individual cells for retrospective readout at the endpoint. To this end, signal integrators have been explored to accumulate reporters in the cytoplasm during a user-defined temporal window^5–10^. Though promising, these approaches fail to resolve individual events and their timing within physiologically relevant time periods. This limitation could be overcome by introducing intracellular memory devices to physically write signals. Efforts have been made to develop such devices using nucleic acids, where cellular events are recorded as edits to their sequences and retrieved via sequencing^11–22^. Nevertheless, owing to moderate editing efficiency, many recorders rely on populational analyses to obtain faithful decoding, losing information at the single-cell level. Their relatively slow editing kinetics also limits their capability to capture events with rapid dynamics^23^. Moreover, sequencing methods employed in signal retrieval typically require the structural disruption of cells or tissues, leading to the loss of spatial information. Though many recorders were able to write the temporal order of events^13,14,19–22^, absolute chronological information and the capability of mapping cell dynamics are still unavailable.

Protein assemblies have recently emerged as a promising complementary memory device. In particular, one-dimensional (1D) assemblies have received the most attention due to their ticker-tape-like linear recording^24,25^. During 1D elongation, cellular histories are recorded as linear segmented fluorescent patterns, and signals are retrievable later by confocal imaging. Leveraging distinct intracellular linear assemblies, Lin *et al.* demonstrated recordings that provide absolute temporal information with a hour-level resolution^24^, and Linghu *et al.* resolved the order of events and showed the potential for *in vivo* implementation^25^. Nevertheless, their practical applications have encountered several obstacles, many of which are intrinsic to linear assemblies. For instance, the filamentous assemblies mechanically perturb the plasma membrane and interrupt cell cycles and migration, and their arbitrary spatial orientations pose challenges for imaging-based signal readout. There has been a strong interest in searching for new recording scaffolds, but candidates available in nature are limited. Among over 200,000 proteins structurally characterized and deposited to the Protein Data Bank, only *ca.* 40 have been reported to assemble into expandable lattices in live cells^26^, and even fewer are suited for developing recorders.

Here, we crafted an *in cellulo* recording platform termed **<u>G</u>**ranularly **<u>E</u>**xpanding **<u>M</u>**emory for **I**ntracellular **<u>N</u>**arrative **I**ntegration (GEMINI) that addresses the above challenges. To overcome the limited candidates of intracellular assemblies, we combined computational design and experimental screening to create new intracellular protein assemblies, greatly expanding the existing toolbox. To overcome the geometry-related constraints, we employed granular assemblies that are smaller than cells in all dimensions during growth, thus minimizing mechanical perturbation and deformation of cells. The granular assemblies grow isotropically in three dimensions (3D), allowing accurate readout independent of their orientations, paving the way for large-scale automated signal readout by imaging. A complete GEMINI system is composed of three components: (1) blank subunits that enable steady and predictable assembly of GEMINI; (2) reporter subunits that transduce cellular events into cytoplasmic fluorescent signals recordable by GEMINI; and (3) timestamp subunits that map the growth of GEMINI for temporal decoding (**Fig. 1a**). During recordings, GEMINI reports the real-time cytoplasmic level of reporter subunits as tree-ring-like fluorescent patterns. The intensity of signal bands in GEMINI quantifies the amplitude of the recorded activities. The incorporation of two or more timestamps allows us to derive the growth profile of individual particles, thus decoding the absolute chronological information of signals.

**Fig. 1.**
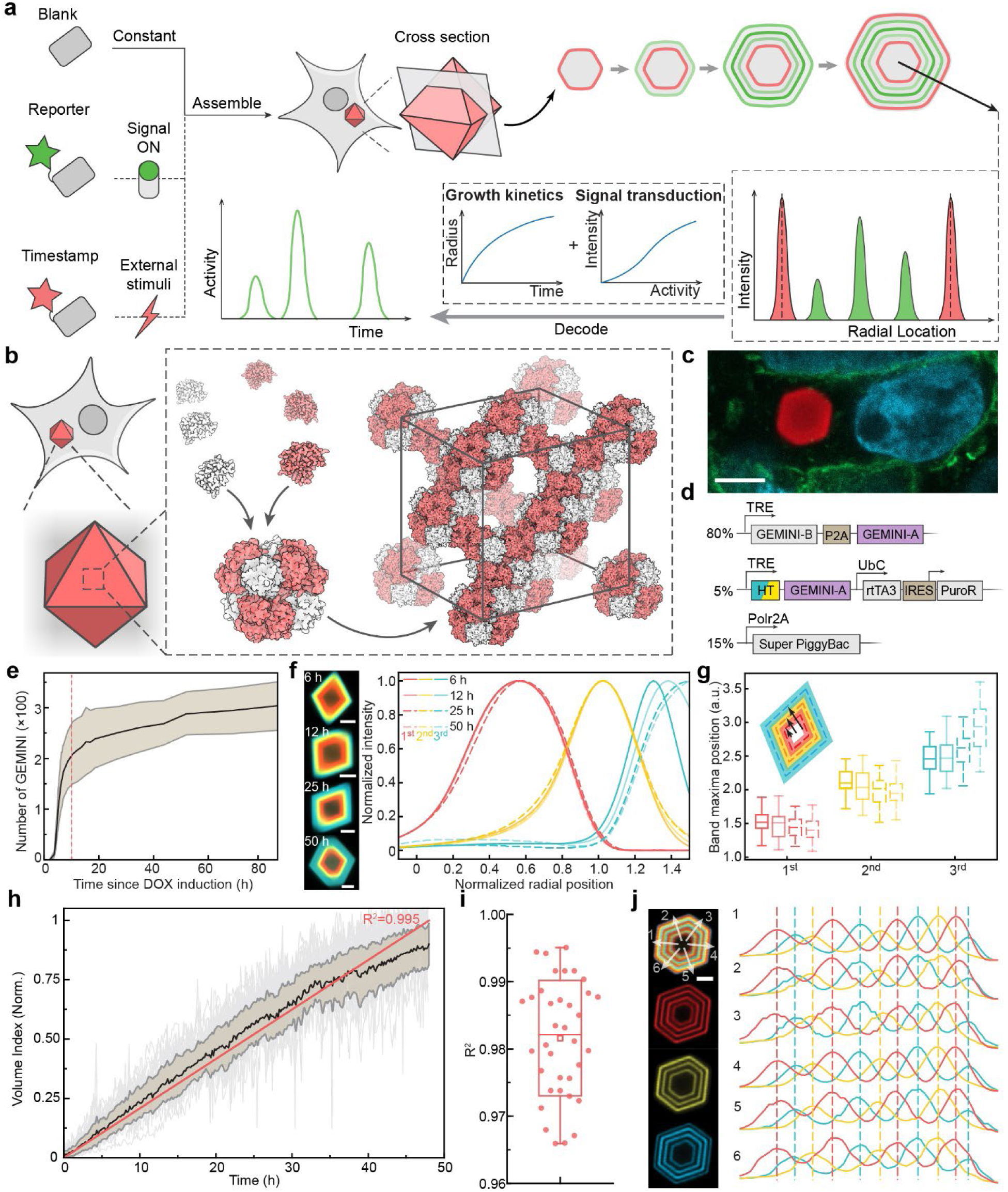
Concept and characterization of GEMINI. **a,** Schematic of the GEMINI recording principle, where blank, timestamp, and reporter components assemble within live cells, forming fluorescence patterns that encode the cell activity history. The encoded information can be decoded based on the known growth profile and activity-dependent signal intensity. **b,** Hierarchical assembly of GEMINI in live cells. Two subunits first assemble into protein cages, which further stack into an expandable lattice. **c,** Image showing GEMINI particle growth in HEK293T cells. Red: GEMINI particles, Cyan: nuclei, Green: cell membrane. Scale bar: 5 μm. **d,** DNA constructs for the development of the clonal HEK cell line expressing GEMINI. **e,** Number of GEMINI particles in the clonal culture over time after DOX induction. Shaded area: ± standard deviation (s.d.). **f,** Images (left) and mean fluorescence profiles (**right**) of GEMINI particles incubated in live cells for various durations before fixation, following identical dye-switch protocols (n = 54/50/52/62 for 6/12/25/50 h). Scale bars: 2 μm. g, Band positions of GEMINI particles in f. **h,** Growth profiles of GEMINI particles in HEK cells (n = 35). Black: mean; shaded area: s.d.; red: fitting. **i,** R² values from the fits of the individual particles’ growth profiles (n = 35). **j,** Image of a GEMINI particle with 11 distinct thin bands (left) and the alignment of fluorescence profiles taken from different directions (right). Scale bar: 2 μm. Box bounds: the 25th and 75th percentiles; whiskers: minimum and maximum; squares: mean; and center lines: median.

Employing the transcription-based reporting mechanism, GEMINI resolves the activation and deactivation of NFκB signaling with a temporal accuracy of *ca.* one hour. Individual events spaced as closely as 15 minutes can be distinguished reliably. GEMINI also achieves a detection sensitivity two orders of magnitude higher than that of cytoplasmic fluorescent transcriptional reporters, with the amplitude of signaling quantifiable. Using an *in vivo* xenograft model, we demonstrated GEMINI’s ability to record both the amplitude and temporal information of inflammation-induced signaling and to correlate cellular responses with local vascular density. We further implemented GEMINI in the mouse brain, showing that GEMINI negligibly impacts neuronal survival and functions, and can be utilized to map the histories of doxycycline (DOX)-mediated transcription and seizure-induced neural activation.

## Results

### Design and screening of the recording scaffold

As intracellular assemblies with expandable lattice are limited in nature, we sought to create new variants by combining computational design and multi-level screenings (**Extended Data Fig. 1a**). In this pipeline, the hierarchical assembly strategy was first adopted, where two-component protein cages with appropriate symmetries were docked into 3D lattices (**Fig. 1b**)^27^. Three distinct lattices were generated and investigated in this work (**Extended Data Fig. 1b**). Before screening, Rosetta-predicted mutations were introduced to the critical residues at the lattice interfaces to expand the library (**See Methods**). We reasoned that intracellular assembly can be mimicked *in vitro* using a buffer with comparable ionic strength to the cytosol. Therefore, we first examined the variants’ assembly by mixing the purified and concentrated subunit solutions *in vitro* in buffers with 0.1-0.5 M NaCl. Among over 30 variants screened *in vitro*, 5 were found to nucleate at a 0.5 M or lower NaCl concentration, which were further evaluated *in cellulo* (**Extended Data Fig. 1c**).

The variants were then expressed in human embryonic kidney (HEK) 293T cells via transient transfection. The two subunits of each variant were linked by a self-cleaving P2A sequence to achieve equimolar expression (**Fig. S1a**). For easier visualization, a subunit tagged with fluorescent protein (FP) was co-expressed in a low fraction (*ca.* 5%) to afford fluorescently labeled assemblies. At 72 hours after transfection, all variants showed clear intracellular precipitants (**Fig. 1c**, **Extended Data Fig. 1d**), proving the efficacy of the design and screening strategies. Assembly-growing cells exhibited morphology comparable to those without assemblies, indicating good tolerance of cells to the cytoplasmic nucleation of the variants. These assemblies, when expressed in live cells, exhibited faceted morphologies, indicating ordered microscopic structures.

*In cellulo* recording favors scaffolds with early nucleation and, ideally, only one assembly per cell. Both features are governed by the nucleation energy barrier, but in opposing ways: a lower barrier facilitates nucleation but leads to multiple nuclei, while a higher barrier results in fewer nuclei but delays nucleation. Among the variants, Lattice #3 v2 and v3 formed multiple assemblies per cell, indicating minimal nucleation barrier, and were therefore excluded from further investigation. Among the remaining candidates, Lattice #1-v2 exhibited an early and synchronous nucleation, making it the most suitable candidate. Notably, it also nucleated efficiently across multiple mammalian cell lines, highlighting its broad applicability (**Fig. S1b**). Therefore, this variant was chosen as the scaffold for GEMINI. While the other variants were not explored further here, they still hold promise for GEMINI recording in specific contexts and further optimization.

### The growth behaviors of GEMINI scaffolds

We next generated and characterized a clonal HEK cell line encoding both the blank and timestamp components of GEMINI, where their expression were mediated by a DOX inducible expression (Tet-ON) system (**Fig. 1d, Fig. S2**)^28^. The timestamp subunit was constructed by fusing HaloTag (HT) to the N terminus of the A chain^29^, which exhibited the lowest impact on their co-assembly, as evidenced by its low equilibrium concentration in the cytoplasm (**Fig. S3a,b**). The same terminus was later used to construct the reporter subunit. This low equilibrium concentration of subunits is critical for achieving sharp transitions of timestamps and distinguishing closely spaced events (**Fig. S3c,d**).

We first characterized the nucleation kinetics in the cell line. Upon the addition of 2 μg mL^-1^ DOX, GEMINI began nucleating in *ca.* 4 hours and plateaued at *ca.* 10 hours (**Fig. 1e, Movie S1**), where most cells grew one assembly in the cytoplasm (**Fig. S4**). This rapid and synchronous nucleation permits simultaneous recording from a large population of cells.

Stable information storage necessitates structurally stable GEMINI with minimal subunit exchange, which was assessed by successively labeling particles with HT ligands (HTL) of distinct colors and monitoring changes in the sharpness and position of color transition bands over time. To achieve this, a fluorophore-free HTL (dark-HTL) was first added to bind the initial HT at induction, creating a dim core in all particles. Sequential staining began at 6 hours post-induction using membrane-permeable Janelia Fluor (JF) conjugated HTLs in the order of JF_669_, JF_608_, and JF_552_, with 6-hour intervals^30,31^. After the last switch, GEMINI particles grew for an additional 6-50 hours before fixation and imaging. The sharpness (**Fig. 1f**) and position (**Fig. 1g**) of the color transition bands remained consistent across all groups, indicating minimal exchange between subunits in GEMINI and those in the cytoplasm. We also observed a noticeable increase in GEMINI’s dimension and a growing outer band over time, indicating continuous lattice expansion (**Fig. 1g**).

To temporally decode signals, the growth profiles of GEMINI should be modeled accurately by timestamps. Linear recorders are advantageous as linear interpolation can be directly employed for decoding, assuming a constant growth rate^24^. Nevertheless, we reasoned that the nonlinear growth of 3D scaffolds could be modeled to achieve a comparable, if not better, temporal resolution.

We first monitored GEMINI growth over 48 hours in HEK cells. The midplane of each particle was mapped via time-lapse imaging and the growth profiles were resolved using an in-house single-particle tracking algorithm (**Movie S2**). As expected, the 3D growth resulted in a nonlinear increase in GEMINI’s radius. We then established a mathematical model to describe the growth (**see Supplementary Note**). In the model, we hypothesized that the subunit’s synthesis rate, cytoplasmic concentration, and degradation rate were all constant in the steady growth phase, affording a constant rate of subunit addition onto GEMINI particles (**Fig. S5**). Based on these assumptions, we derived an equation that linked the radius (R) of particles with time (t): *R* = (*K* + *A*)^1/3^, where *K* and *A* are constants (**See Supplementary Note**). The model was validated by fitting the mean linearized growth profile (defined as the volume index, **See Methods**), showing an R^2^ of 0.995 (**Fig. 1h**). We also examined the fitting of growth profiles from individual particles, which yielded a mean R^2^ of 0.982 (**Fig. 1i**). Minor deviations from linearity were observed near the end of the tracking period, likely due to reduced accuracy in capturing the true midplane as particles grew larger, as well as a possible increase in the energy barrier for assembly resulting from defect accumulation. In addition, fluctuations in individual growth profiles were attributed to tracking inaccuracies and particles’ out-of-plane motion during imaging, rather than intrinsic structural instability. The results demonstrate the high accuracy of the model in describing GEMINI growth, which also corroborates the assumption of steady protein expression, an important indicator of normal cell metabolism.

Scalable and automated signal readout is crucial for *in cellulo* recording, necessitating the imaging of fluorescent patterns regardless of the recorders’ orientation. Granular recorders, with their isotropic growth, are well-suited for this purpose. GEMINI particles maintain an octahedral shape throughout various growth stages, indicating the uniform addition of subunits onto 111 facets (**Extended Data Fig. 1e-j**). To assess recording in various directions, we rapidly switched dyes to create 11 thin bands mimicking patterns from dynamic signaling (**Fig. 1j**). Any non-uniform growth would be evident from pattern misalignment among directions. For a representative particle, fluorescence profiles measured along six distinct orientations were well aligned, demonstrating isotropic growth and orientation-independent recording. In contrast, linear recorders encoded information in a single direction, and a slight tilt from the focal plane significantly deteriorated imaging quality (**Extended Data Fig. 2a**). In 3D culture that mimics the growth in tissue, considerable number of linear recorders oriented out-of-plane, posing challenges on scalable decoding (**Extended Data Fig. 2e**). Although volumetric imaging can map out-of-plane recorders with minimal impact on decoding precision, it substantially lowers throughput (**Extended Data Fig. 2f-h**). Collectively, these results demonstrate that GEMINI has favorable growth behaviors to serve as an intracellular recorder and is well-positioned for scalable recording and high-throughput signal readout.

### GEMINI minimally impacts cell processes

An ideal recorder should exhibit negligible interference with essential cell processes. We first assessed the impact of GEMINI nucleation on survival of HEK cells using a live/dead assay (**Fig. S6**). After growing GEMINI for 4 days, no apparent increase in cell death was observed, indicating its low cytotoxicity (**Fig. S6c-e**). Next, we profiled the morphological features of subcellular structures during GEMINI growth via the Cell Painting assay^32^. While most compartments exhibited minimal structural changes, a decrease in the nuclear area and a subtle increase in mitochondrial area were observed at 48 hours after GEMINI expression (**Fig. S7a-e**), which could be due to potential physical interactions and changes in confluency. Nevertheless, these changes were not observed in U2OS cells expressing GEMINI (**Fig. S7f-j**).

Moreover, we examined cell proliferation by tracking cell density over time, where comparable proliferation rates were observed between cells with and without GEMINI nucleation (**Extended Data Fig. 3a,b**), suggesting a negligible influence of GEMINI growth on cell proliferation. We then further investigated the influence of GEMINI on division at the cellular level, where no apparent disruption of mitosis and cytokinesis was observed following GEMINI nucleation. After division, GEMINI particles entered one of the daughter cells (**Extended Data Fig. 3c**). In contrast, cytoplasmic nucleation of linear recorders like iPAK4 was found to disrupt proliferation by preventing membrane abscission (**Extended Data Fig. 3e,f**).

### Temporal resolution of GEMINI recording

We then investigated if GEMINI can temporally map cellular events. To assess decoding accuracy and resolution, we first introduced an artificial intracellular signal by adding an HTL dye that mimicked the appearance of biomolecules in the cytoplasm (**Fig. 2a**). Since HTL binds to HT almost instantaneously, the time of addition represents the ground truth of the signal onset^24^. Before recording, GEMINI were pre-incubated in an HTL, generating a dim core. The first switch (yellow) occurred at 8 hours post-induction, serving as the first timestamp and marking t=0. The signal dye (violet) was added at t=2-8 hours to different groups, followed by a final switch at t=11 hours as the second timestamp (blue). Cells were incubated for another 11 hours before fixation, and over 200 particles from each group were batch-imaged and analyzed.

**Fig. 2.**
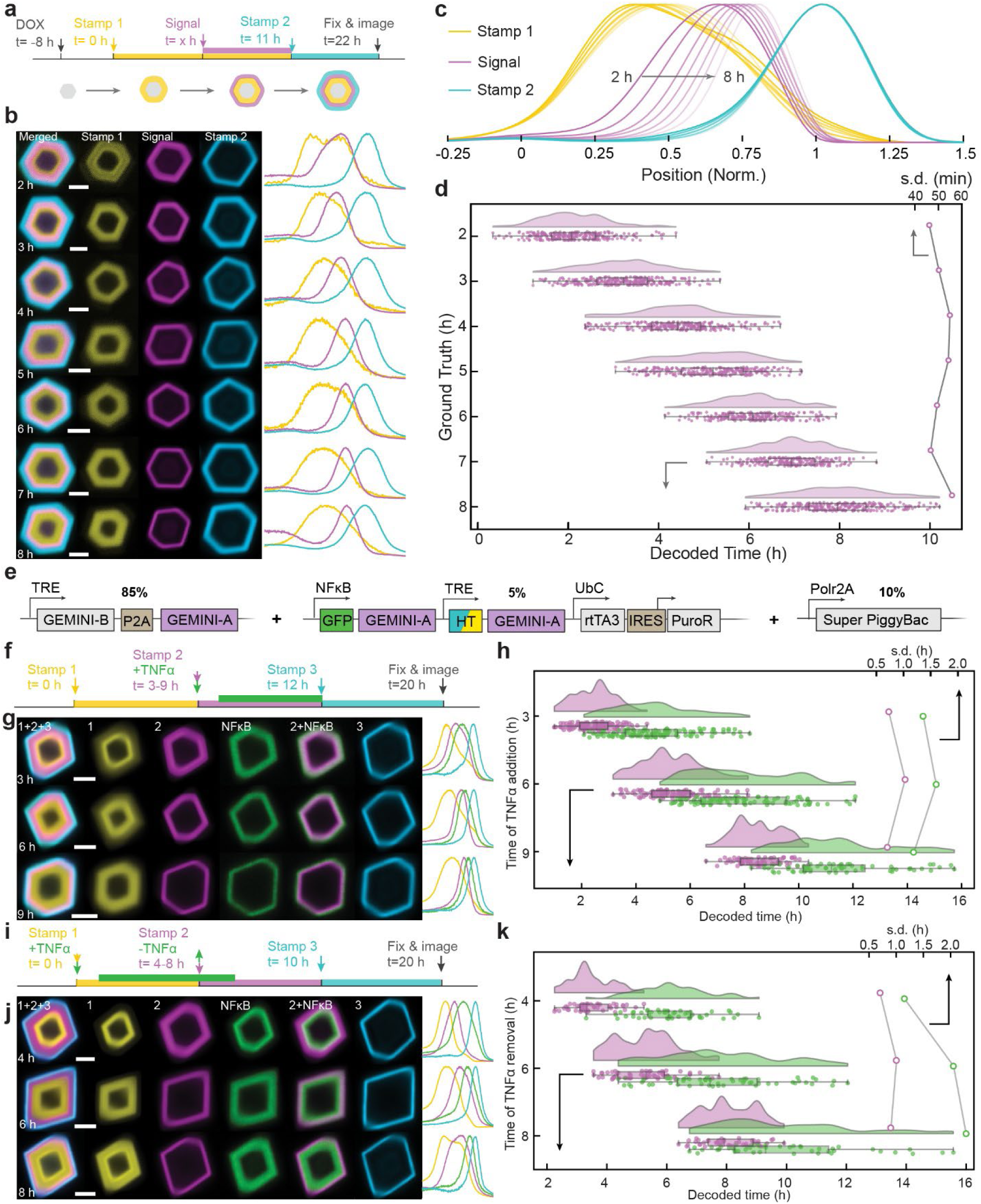
I*n cellulo* recording using GEMINI. **a,** Experimental procedure for testing the precision of temporal decoding. The onset of signal-mimetic dye addition events (blue) is decoded using two timestamps: red (t = 0) and yellow (t = 10 h). **b,** Images (left) and fluorescence profiles (right) of GEMINI particles with signals introduced at t = 2–8 h. Scale bars: 2 μm. **c,** Mean fluorescence profiles of the timestamp and signal channels. **d,** Decoded onsets from individual GEMINI particles (bottom-x, n = 258/275/279/261/225/219/275 for t = 2/3/4/5/6/7/8 h) and the standard deviations (top-x) for each group. **e,** DNA constructs for the development of NFκB-GEMINI cell lines. **f,** Experimental procedure for recording NFκB upregulation in response to TNFα (green). A dye is added along with TNFα as stamp #2 (violet). Both NFκB activation and stamp #2 are decoded using stamps #1 (yellow, t = 0) and #3 (blue, t = 12 h). **g,** Images (left) and fluorescence profiles (right) of GEMINI particles recording NFκB activation at t = 3, 6, and 9 h. Scale bars: 2 μm. **h,** Decoded times (bottom-x) and standard deviations (top-x) for NFκB activation (green) and stamp #2 (violet) signals from individual GEMINI particles (n = 106/79/47 for t = 3/6/9 h). **i,** Experimental procedure for recording NFκB deactivation after TNFα removal (green). A dye is added as stamp #2 (violet) when TNFα is removed. NFκB deactivation and stamp #2 are decoded using stamps #1 (yellow, t = 0) and #3 (blue, t = 10 h). **j,** Images (left) and fluorescence profiles (right) of GEMINI particles recording NFκB deactivation at t = 4, 6, and 8 h. Scale bars: 2 μm. **k,** Decoded times (bottom-x) and standard deviations (top-x) for NFκB deactivation (green) and stamp #2 (violet) signals from individual GEMINI particles (n = 50/59/45 for t = 4/6/8 h). **d,h,k,** Box bounds: the 25th and 75th percentiles; whiskers: minimum and maximum; and center lines: median.

For all resolved patterns, signal onsets were consistently bordered between the two timestamps, demonstrating robustness in capturing the order of events (**Fig. 2b**). The fluorescence profiles from the groups were normalized and compared, where the signal onsets of the individual groups appeared at expected locations corresponding to the time of signals (**Fig. 2c**), indicating successful recording of temporal information. Using our growth-prediction model, we temporally decoded the signals recorded in individual particles (**Fig. 2d**, **see Supplementary Notes**). The results showed close agreement with the ground truths. The timing standard deviation was within the sub-hour range for all groups. We further analyzed the time error between the decoded time and the ground truth, where **75.9%** of the single-particle decoding results fell within ±1 hour of the ground truth, **98.2%** within ±2 hours, and **100%** within ±3 hours, indicating precise temporal decoding is possible at cellular level (**Fig. S8**).

As the radius of GEMINI grows nonlinearly, we expect a decreased temporal resolution if signals are recorded at the later stage of assembly. To quantify this, we initiated the recording at 48 hours post expression and recorded for another 36 hours (**Extended Data Fig. 4a**). Despite a much larger core and slower radial growth (**Extended Data Fig. 4b**), a timing standard deviation of 2-4 hours was still achieved (**Extended Data Fig. 4d**), and most particles (92.9%) still exhibited a time error no more than 6 hours (**Extended Data Fig. 4f**), showing GEMENI’s capability to record at an extended window.

### GEMINI resolves physiologically relevant signals

We then assessed whether physiological events could be captured by GEMINI via activity-dependent expression of the reporter subunit. This approach has three key advantages: (1) transcription directly reflects signaling cascade output, closely representing upstream pathway activity and endogenous protein expression; (2) a large dynamic range is attainable, favorable for GEMINI recording; and (3) transcription-based reporting is modular and easily adaptable to report diverse pathways of interest^33,34^.

As a proof-of-concept, we attempted to record transcriptional dynamics mediated by NFκB signaling, exploiting a synthetic promoter that combines tandem repeats of the NFκB response element (NFκB-RE) with a minimal promoter (P_min_) to drive the expression of the reporter subunit (**Fig. S9a**)^11^. Before recording, we first examined orthogonality between GEMINI and NFκB signaling. We quantified the phosphorylation of IκBα, a key element in the NFκB signaling cascade, via Western blotting (**Fig. S10**). Comparable response to tumor necrosis factor-alpha (TNF-α) were found between the groups with and without *in cellulo* GEMINI, indicating minimal impact of GEMINI on the pathway activation. It is noted that a modest increase in basal IκBα level and its phosphorylation was observed in GEMINI-expressing cells, which may indicate a low-level cellular stress associated with particle formation. Then, we investigated the reporter expression in response to NFκB activation in HEK cells without growing GEMINI (**Fig. S11**). Before activation, reporter subunits had a low basal level in the cytoplasm, and 12 hours of incubation in 10 ng mL^-1^ TNF-α increased the fluorescence intensity by *ca.* 7-fold.

We then developed a clonal HEK cell line encoding all three components of GEMINI for DOX-induced recording of NFκB signal, termed NFκB-GEMINI cell line (**Fig. 2e**, see Methods). Upon DOX induction, dark-HTL was added to obtain a dim core, followed by switching to an HTL dye (yellow) 6 hours after induction as the first timestamp (t=0). TNF-α was then added at t=3, 6, or 9 hours to separate groups (green), together with another HTL dye (violet) as an accompanying timestamp. At t=12 hours, all groups were switched to the third HTL dye (blue) as the last timestamp, and cells were incubated for another 8 hours before fixation (**Fig. 2f**). In the images of GEMINI particles (**Fig. 2g**) and the normalized mean fluorescence profiles (**Fig. S12a**) from the groups, the onset of NFκB activation appeared at the anticipated locations in all three conditions, where the signal appeared slightly later than the accompanying timestamp, owing to delays by transcription and translation. We then temporally decoded both the NFκB activation and its accompanying timestamp. The mean decoded times for the timestamps closely agreed with the ground truths, indicating the minimal impact of NFκB activation on the GEMINI growth and decoding accuracy. The decoded times for NFκB activation were 4.82±1.35, 7.72±1.59, and 10.71±1.18 hours for t=3, 6, and 9 hours, respectively, showing a consistent delay of *ca.* 2 hours to TNF-α induction. It is noted that the temporal distribution of NFκB signals is broader than that of the second timestamps, which could be attributed to the heterogeneity in cellular responses to TNF-α.

Similar to recording signal activation, we reasoned that deactivation, the other key aspect of signaling dynamics, is also resolvable. This is because particles served as a reservoir for reporter-subunit uptake, significantly accelerating their removal from the cytoplasm. Indeed, a sharp decay of the NFκB signal was observed consistently in GEMINI particles. To further test this, we designed a recording course similar to that for NFκB activation but having TNF-α constitutively applied while emulating the deactivation by its removal (**Fig. 2i**). In the test, the first (yellow) and last (blue) timestamps marked t=0 and 10 hours, respectively, and TNF-α was removed from different groups at t=4, 6, and 8 hours (green), with the addition of an accompanying HTL dye (violet) marking TNF-α removal (**Fig. 2j**). Accurate timing of the deactivation onset, defined as the peak of NFκB bands in particles, was achieved (**Fig. 2k, Fig. S12b**). Using the accompanying timestamps as references, we found the reporter subunit peaked at *ca.* 2 hours after TNF-α removal (6.04±1.15, 7.75±2.06, and 9.84±2.28 hours for the t=4-, 6-, and 8-hour groups, respectively), owing to the continued expression before mRNA degradation. This is difficult to achieve using conventional fluorescence or luciferase transcriptional reporters, due to the relatively long half-life and cytoplasmic retention of the reporter proteins.

### GEMINI resolves fast dynamics and signal amplitudes

GEMINI’s ability to resolve both signal activation and deactivation presents an opportunity to map signaling dynamics that involve repetitive ON and OFF cycles. To examine this, we simulated NFκB dynamics by adding and removing three doses of 10 ng mL^-1^ TNF-α over 40 hours (**Fig. 3a**), applying timestamps with each addition and removal. All events were successfully captured by individual GEMINI particles (**Fig. 3b**), accurately mapping the timing of both the ON and OFF phases (**Fig. 3c**). These results collectively demonstrate GEMINI’s capability to temporally resolve signaling dynamics of physiological relevance in live cells.

**Fig. 3.**
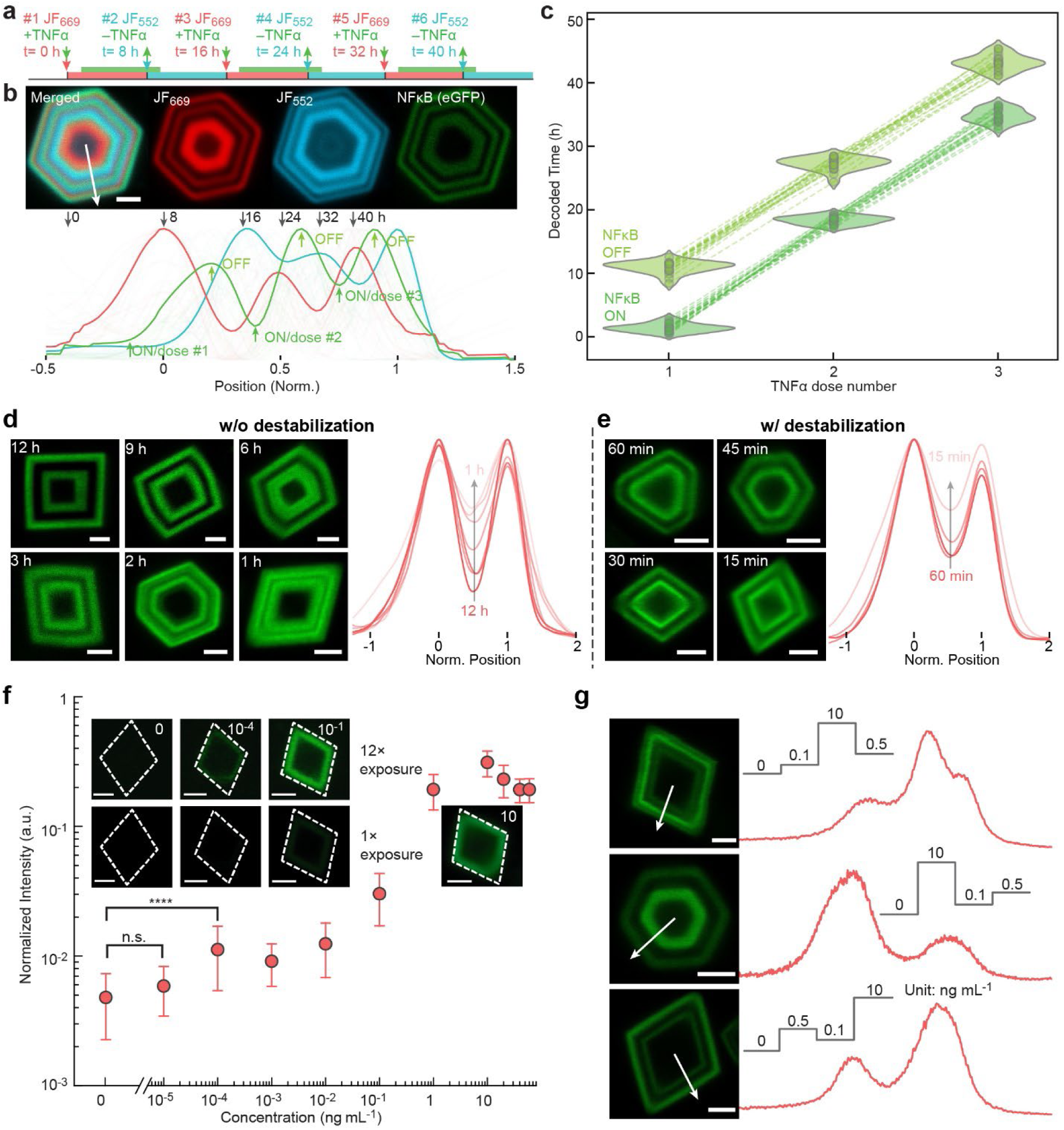
Recording multiple events and dynamics in live cells. **a,** Experimental procedure for multi-event recording. Three ON-OFF cycles were introduced, with a stamp applied at each TNFα addition and removal. **b,** Image (top) and the mean fluorescence profiles (bottom) of the timestamp and signal channels. Scale bar: 2 μm. **c,** Decoded times for ON and OFF events during each TNFα dose (n=21 particles). **d,** GEMINI particles encoding dual NFκB activation signals with decreasing intervals (*left,* 12 to 1 hours) and the mean fluorescence profiles (*right,* n = 15/17/13/16/8/8 for 12/9/6/3/2/1 h). **e,** GEMINI particles encoding boosted dual NFκB activation signals with decreasing intervals (left, 60 to 15 minutes) and the mean fluorescence profiles (right, n = 18/17/19/20 for 60/45/30/15 min). Scale bars: 2 μm. **f,** Dose-dependent intensity of signals recorded by GEMINI. Insets: images of single GEMINI particles capturing signal amplitudes at various doses. The images in the top row are the same particles as the bottom, but with a 12 times of exposure level. Circles: mean; whiskers: s.d.. **g,** GEMINI particles (left) recording varied signal amplitudes and the corresponding fluorescence profiles (right). TNFα doses of 0/0.1/0.5/10 ng/mL were applied in distinct sequences. Scale bars: 2 μm.

We then characterized the fastest dynamics GEMINI could resolve by measuring the separation of two NFκB activation peaks recorded at various time intervals. After 8 hours of GEMINI expression, cells were incubated successively in two doses of 10 ng mL^-1^ TNF-α for 8 hours each, with decreasing intervals from 12 to 1 hour (**Fig. 3d**). In representative particles from each group, a clear correlation was observed between the interval length and the separation of the two peaks. At intervals of 6 hours or longer, the two bands were separated with minimal overlap. The bands started to merge as the interval decreased. When a 1-hour interval was applied, two bands were substantially merged but still distinguishable.

Although capturing the hour-level dynamics is adequate for mapping diverse cellular events, there is significant scientific interest in resolving even faster dynamics. We hypothesized that this could be achieved by accelerating the turnover of cytoplasmic reporter subunit and its mRNA (**Fig. S13a**). This would produce a sharp decay in the signal band upon deactivation and reset the system more rapidly for capturing subsequent events. To achieve this, we incorporated the P1N4 domain, which was reported to enhance protein turnover by an order of magnitude^35^, to destabilize both the reporter subunit and its mRNA. A clonal HEK293T cell line expressing the destabilized reporter subunit, termed NFκB-GEMINI-Boost, was developed accordingly (**Fig. S13b**). These modifications enabled the resolution of even faster dynamics, where distinct signaling events separated by as short as 15 minutes could still be distinguished (**Fig. 3e**).

Furthermore, we investigated whether the signal amplitude could be quantified by the band intensity. A dose-response test on NFκB signaling was performed at TNF-α doses ranging from 10^-5^ to 60 ng mL^-1^, with a TNF-α-free group included for comparison (**Fig. 3f**). The peak of NFκB bands, normalized by the intensity of HTL staining, measured signal amplitude. We found cells responded to a dose as low as 10^-4^ ng mL^-1^, two orders of magnitude lower than the detection limit of the cytoplasmic transcriptional reporter (**Fig. S11b**). GEMINI recording also exhibited a dynamic range of *ca.* 60-fold, over 7 times higher than the cytoplasmic benchmark. The improved detection limit and dynamic range can be attributed to GEMINI’s concentrating effect, where cytoplasmic signals are condensed and amplified. We then sought to record the response to trains of stimuli, where differential amplitudes were induced by low (0.1 ng mL^-1^), medium (0.5 ng mL^-1^), and high (10 ng mL^-1^) doses of TNF-α at various sequences. Cells were incubated in each concentration for 8 hours (**Fig. 3g**). In representative particles, distinct plateaus of fluorescent intensities were observed in the expected order, showing GEMINI’s capability to resolve and quantify dynamics with varying signal amplitudes.

### GEMINI records inflammatory responses *in vivo*

To evaluate whether GEMINI could record physiologically relevant signaling events *in vivo*, we first established a xenograft model to monitor inflammation-induced NFκB activation in immunodeficient mice^11^. This was achieved by subcutaneously implanting the NFκB-GEMINI cell line. Inflammation was induced through intraperitoneal (i.p.) injection of lipopolysaccharide (LPS), a well-established method for triggering acute inflammatory responses via activation of Toll-like receptor 4 (TLR4) on host immune cells^36^. Upon stimulation, host-derived cytokines, including TNF-α, diffuse into the xenograft through circulation, activating NFκB signaling in the GEMINI-expressing cells.

We first examined if GEMINI-expressing cells in xenograft could respond to inflammation *in vivo*. Ten days post-implantation, after the xenograft had developed into a palpable tumor, DOX was administered to initiate GEMINI expression (**Fig. 4a**). LPS was then administered at a dose of 3 mg kg^-1^ six days after DOX induction (designated as day 0), and tumors were harvested on day 4 for GEMINI (HTL dye) and vasculature (anti-CD31 staining) labeling. Widespread GFP signal, indicative of NFκB activation, was observed throughout the tumor tissue (**Fig. 4b, Extended Data Fig. 5**). Notably, GFP intensity exhibited substantial spatial variation, reflecting heterogeneity in the cellular response to the systemic inflammatory signal.

**Fig. 4.**
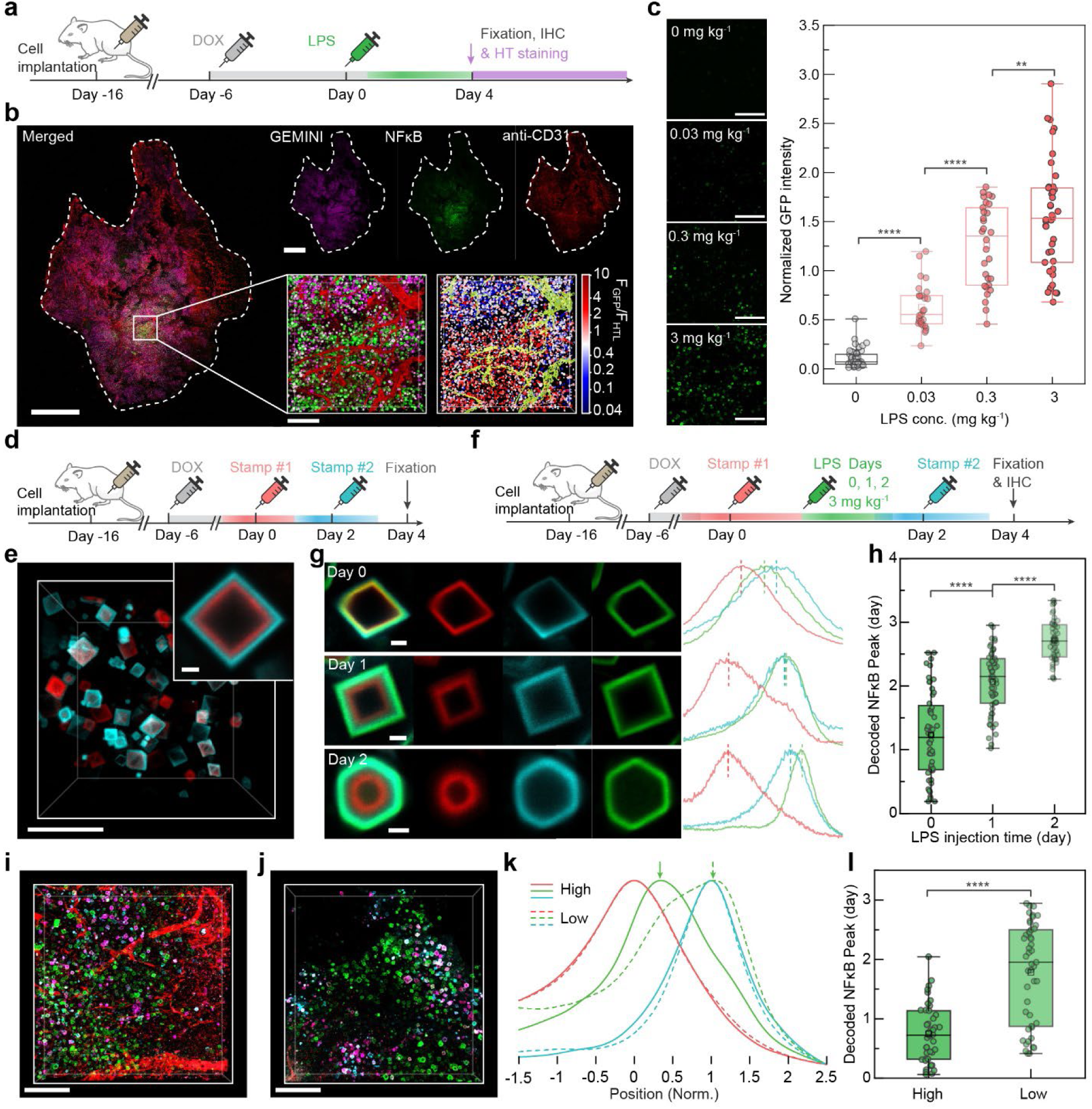
In vivo recording of inflammatory response. **a,** Experimental procedure for *in vivo* recording of LPS-induced inflammation. **b,** Section of a HEK293T xenograft showing GEMINI recording of NFκB activation following systemic injection of 3 mg kg^-1^ LPS. GEMINI: violet, NFκB: green, vasculature (anti-CD31): red. Insets: magnified 3D image (*left*) and the corresponding heatmap displaying the normalized GFP intensity (F_GFP_/F_HTL_) of individual particles (*right*). Scale bars: 1 mm, 100 μm (*insets*). **c,** Images of GEMINI recording NFκB signal at various LPS doses (*left*) and the dose-dependent signal intensity (*right*). (n=61/27/31/42 for 0/0.03/0.3/3 mg kg^-1^ LPS, 3 mice per group). Scale bar: 100 μm. **d,** Experimental procedure for *in vivo* timestamping via systemic injection of HTL dyes. **e,** Image of GEMINI particles in xenografts labeled with two timestamps; inset: magnified image of a GEMINI particle. Scale bars: 20 μm, 2 μm (*inset*). **f,** Experimental procedure for recording NFκB signal with LPS administered at various times. **g,** Images (*left*) and fluorescence profiles (*right*) of GEMINI particles recording NFκB activation in mice injected with LPS on days 0, 1, or 2. Scale bar: 2 μm. **h,** Decoded timing of NFκB activation peaks in groups receiving LPS on days 0–2 (n = 52/59/55 for t = 0/1/2 days; 3 mice per group). **i,j,** 3D images of xenograft tissue with high (**i**) and low (**j**) vascular density. LPS was administered on day 0. Scale bars: 100 μm. **k,** Mean fluorescence profiles of NFκB signals recorded in GEMINI particles located in regions with high (solid) and low (dashed) vasculature density. **l,** Decoded peak timing of NFκB activation for GEMINI particles from high- and low-vascularization regions (n = 42/47 for high/low; 3 mice). **c,h,i,** Box bounds: the 25th and 75th percentiles; whiskers: minimum and maximum; squares: mean; and center lines: median.

We next tested whether GEMINI could resolve varying levels of inflammation *in vivo*, as demonstrated *in vitro* with different TNF-α doses. A range of LPS concentrations (0.03–3 mg kg^-1^) was administered on day 6, with a control group receiving equivolume saline. Analysis of GFP intensity (normalized to HTL dye staining) from individual particles revealed a clear dose-dependent response (**Fig. 4c**), confirming GEMINI’s ability to encode relative levels of inflammation.

To resolve absolute timing, we developed an *in vivo* timestamping strategy. While the *in vitro* sequential dye-switch approach is not directly applicable, we leveraged the high bioavailability of several HTLs, enabling timestamping via retro-orbital injection^31^. Due to the rapid clearance of HTLs from circulation, each administration produces a transient pulse that labels the particle’s instant surface with a sharp fluorescent band (**Extended Data Fig. 6**, **see Supplementary Notes**). By administering HTL dyes at distinct time points at a dose of 10 nmol per animal, sharp timestamps were introduced *in vivo* (**Fig. 4d**). Two timestamps applied 48 hours apart produced clearly distinguishable layers in the resulting GEMINI particles (**Fig. 4e**).

We then examined GEMINI’s ability to retrospectively decode the timing of inflammatory events. After initiating GEMINI expression, mice were divided into three groups, receiving 3 mg kg^−^¹ LPS on days 0, 1, or 2, respectively. Timestamps were applied on days 0 and 2 for temporal decoding (**Fig. 4f**). As expected, later LPS administration corresponded to a consistent outward shift of the GFP band relative to the timestamp markers (**Fig. 4g**). Temporal decoding of individual particles using the model validated in culture showed that the NFκB signal peaked *ca.* one day after LPS injection in each group (**Fig. 4h**). We noted that, compared to *in vitro* measurements, the timing showed larger variation within each group. We attribute this variability to a combination of factors, such as heterogeneous cytokine diffusion within the tumor, diverse cellular responses to inflammatory cues, and lower precision of *in vivo* timestamping.

Among factors causing the observed heterogeneity, we reasoned that differences in local vascularization played a critical role, which could influence cytokine accessibility and therefore timing of NFκB activation. Indeed, histological examination revealed significant variation in vasculature density across regions (**Fig. 4i,j**). To test this, we decoded NFκB signals in GEMINI particles from regions with high and low vascular density, defined by the volume occupancy of >8% and <2%, respectively. A noticeable delay in NFκB signaling was observed for the poorly vascularized region (**Fig. 4k**), where the mean peak time were *ca.* 1.8 days post LPS injection as compared to 0.7 days for the well vascularized region (**Fig. 4l**). Such delay in NFκB activation could be attributed to the longer time necessary for cytokine diffusion to the region. This study highlights the correlation between vascularization and cytokine diffusion within tumors and showcases a quantitative analysis of signaling dynamics within tumors in response to systemic signals.

### Implementing GEMINI to the mouse brain

We then asked if GEMINI could be implemented into native tissues, such as neurons within the mouse brain. To test this, we created a single adeno-associated viral vector (AAV) encoding both the blank and timestamp subunits (btAAV) and delivered it at a dose of 1×10^12^ vector genomes (vg) per mouse via intracranial injection (**Extended Data Fig. 7a**). Sham groups were injected with equivolume saline for comparison. GEMINI nucleated efficiently across brain regions (**Extended Data Fig. 7b**), with particles appearing as early as day 5. Particle size increased over time, indicating their continued growth (**Extended Data Fig. 7c**). Similar to *in vitro* growth, most neurons nucleated a single GEMINI particle (**Extended Data Fig. 7d**). Segmentation enabled accurate registration of each particle to its corresponding neuron, preserving their spatial information within the tissue (**Extended Data Fig. 7e-h**).

To evaluate biocompatibility, we assessed neuronal and inflammatory responses following GEMINI expression. Histological analysis of the hippocampal CA1 region from day 7 to 14 post-injection revealed no significant changes in neuronal density or astrocyte reactivity despite increased GEMINI particle density (**Extended Data Fig. 8a,b**). By day 14 of expressing GEMINI in brain regions such as the cortex and hippocampus, high-density GEMINI particles were consistently observed (**Fig. 5a**). On day 14 post-expression, histological analyses of various subregions within the hippocampus revealed no significant differences from sham-injected controls in neuronal density or astrocyte reactivity (**Fig. 5b,c; Extended Data Fig. 8e,f**). Furthermore, we analyzed morphological features of GEMINI-expressing neurons, focusing on soma and nucleus (**Extended Data Fig. 9a-g**). While soma sphericity and nuclear volume remained unchanged, we observed a slight increase in soma volume and a modest decrease in nuclear sphericity (**Extended Data Fig. 9d-g**). These subtle changes may reflect the occupation of cytoplasmic space and nuclear deformation due to large GEMINI within some cells. The results indicate that *in cellulo* GEMINI exhibits a low impact on neuronal survival and morphology.

**Fig. 5.**
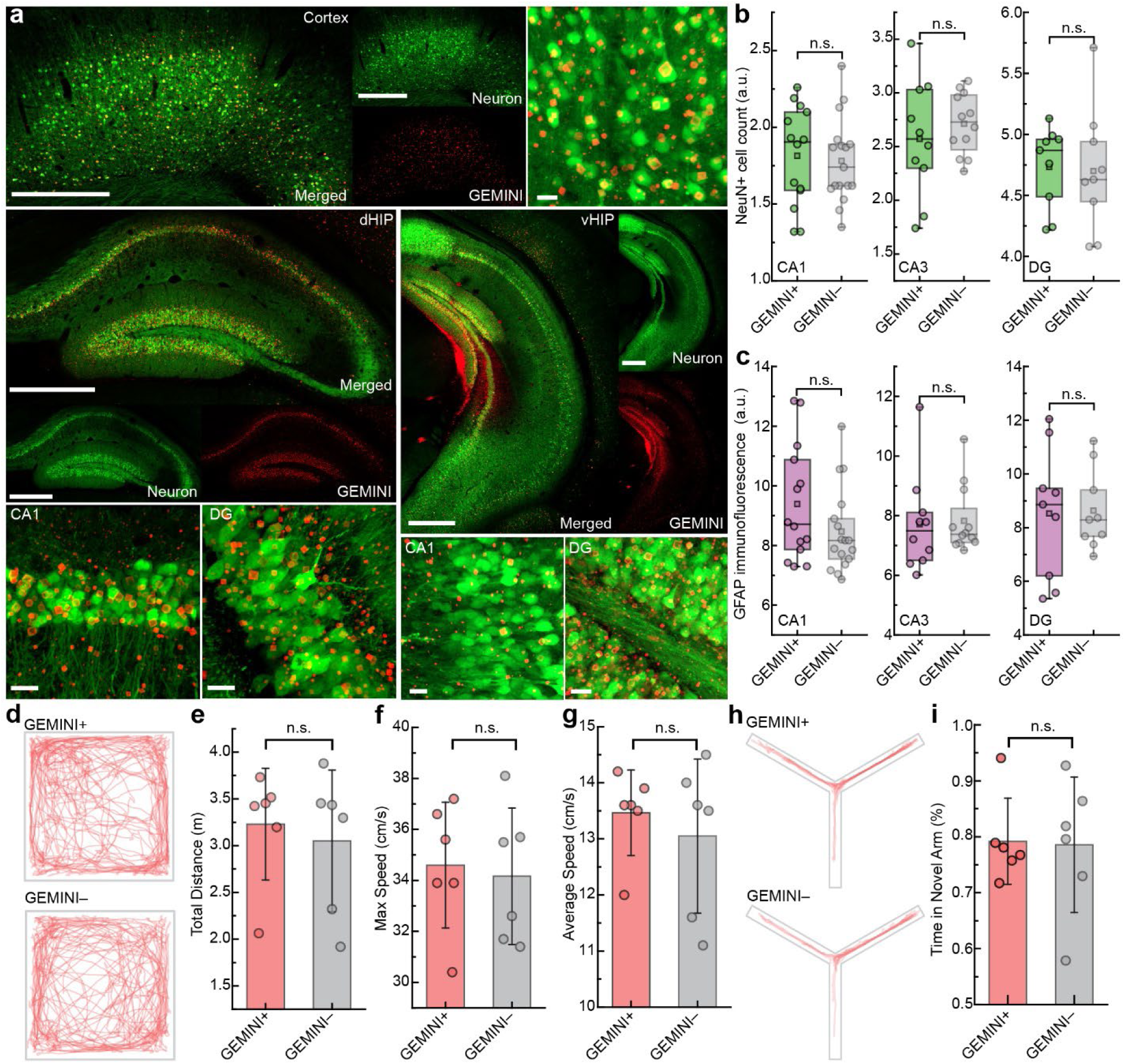
Implementing GEMINI to the mouse brain. **a,** Images showing high-level GEMINI expression in the cortex (top) and hippocampus (bottom). Scale bars: 500/20 μm for low/high magnification. **b,c** Comparison of the neuronal density (NeuN+ cells) and immune response (GFAP immunofluorescence) between groups with and without GEMINI expression (n=5 mice per group). **d,** Traces of mice with and without GEMINI expression in an open-field arena. **e-g,** Comparison of total traveling distance (**e**), maximal speed (**f**), and average speed (**g**) between the GEMINI+ and – groups (n=6 mice per group). **h,** Traces of mice with and without GEMINI expression in a Y-maze. **i,** Comparison of time spent in the novel arm between the GEMINI+ and – groups (n=6 mice per group). **b,c,** Box bounds: 25th and 75th percentile; whiskers: minimum and maximum; squares: mean; and center lines: median. **e-g, i, bars:** mean; whiskers: s.d..

To assess the functional impact, we performed *in vivo* two-photon calcium imaging in *thy1-jRGECO1a* transgenic mice injected with btAAV in the primary visual cortex (**Extended Data Fig. 9h–l**). GEMINI-expressing neurons exhibited calcium transient patterns and ΔF/F of calcium responses comparable to GEMINI– neurons, suggesting minimal impact of GEMINI on spontaneous neuronal activities.

We then evaluated whether the *in vivo* GEMINI expression impacted behavior. The btAAV was injected bilaterally into brain regions involved in motor control and memory, including the primary motor cortex (M1), dorsal (dHIP), and ventral hippocampus (vHIP), with the behavioral tests performed on day 14. In open-field tests, no differences in total traveling distance, maximal speed, or average speed were observed between GEMINI and sham groups (**Fig. 5d-g**), suggesting preserved motor function. In the Y-maze test, mice from both groups spent comparable duration in the novel arms, indicating that GEMINI minimally impacts short-term memory (**Fig. 5h,i**).

To further evaluate fine motor coordination, we unilaterally injected btAAV into the M1 region. A positive control group expressing diphtheria toxin subunit A (dtA) and a negative control group receiving saline were also included (**Extended Data Fig. 10a**). Horizontal ladder rung walking tests were performed on day 14 (**Extended Data Fig. 10b,c**). GEMINI-expressing mice performed comparably to saline controls, while dtA-expressing mice exhibited impaired coordination and longer passing time (**Extended Data Fig. 10d,e**). Postmortem analysis revealed preserved brain symmetry in GEMINI mice, whereas dtA mice showed mild shrinkage in the injected hemisphere (**Extended Data Fig. 10f**). Histological analysis further confirmed minimal neuronal loss associated with GEMINI expression (**Extended Data Fig. 10g,h**).

Together, these results indicate that *in vivo* expression of GEMINI in the brain exhibits biocompatibility and has low impact on neurons in cellular, functional, and behavioral levels.

### GEMINI records transcription and seizure-induced activity in the mouse brain

Before recording cellular events in the brain, we first examined if the systemic timestamping demonstrated in xenograft is also applicable to the brain. The blood-brain-barrier (BBB)-crossing HTL dyes, including JF_669_ and JF_552_, were employed in this study^37,38^. For both dyes, a dose of 20 nmol per mouse was found adequate to afford a sufficiently bright band while avoiding competition with the later stamps. As a demonstration, we injected JF_669_ and JF_552_ with a 24-hour interval (**Fig. 6a**). Two sharp peaks were found in the expected order, demonstrating successful timestamping (**Fig. 6b, c**).

**Fig. 6.**
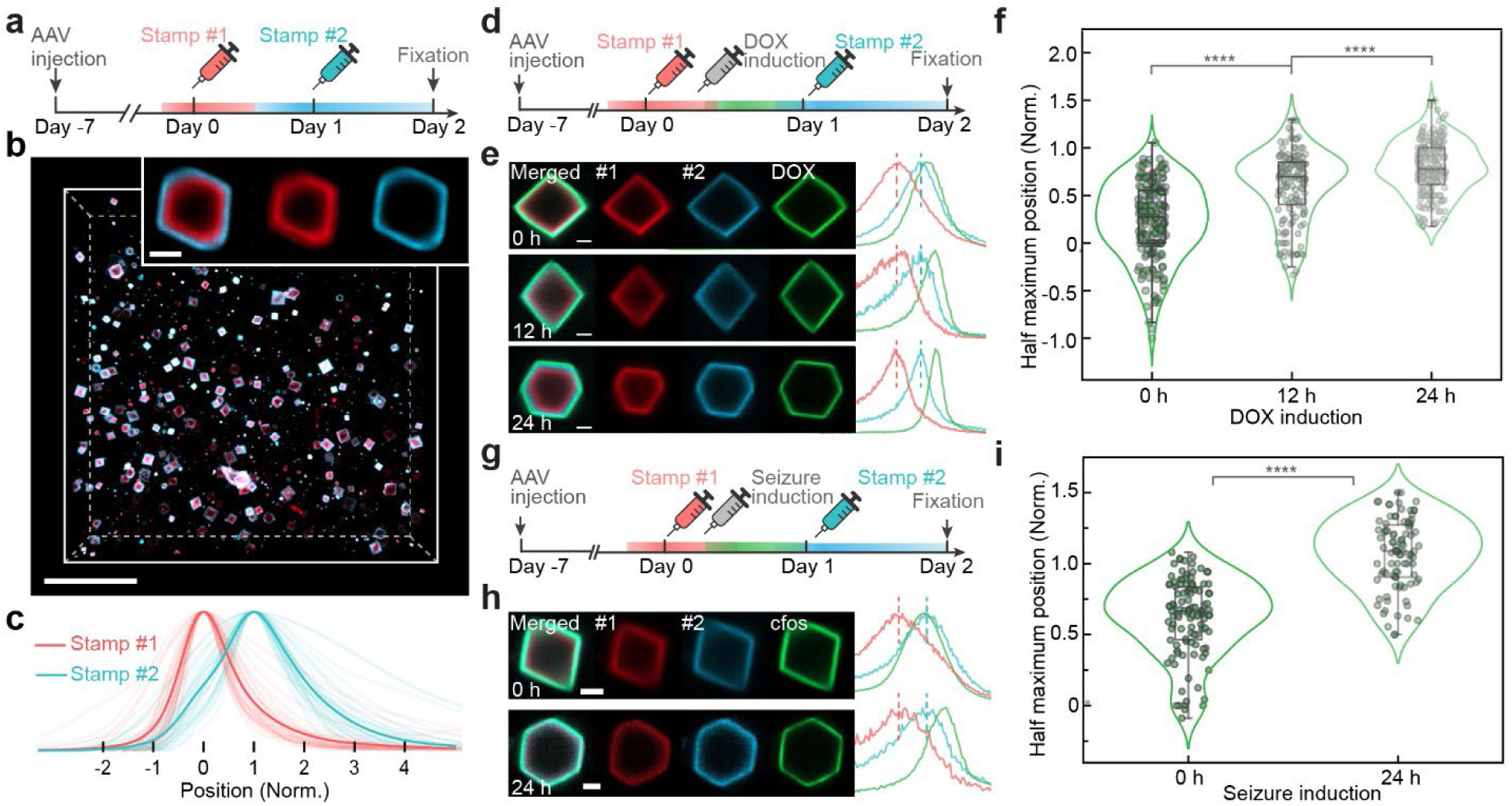
Recording in the mouse brain. **a,** Experimental procedure of *in vivo* timestamping in the brain, where dyes are systemically delivered via retro-orbital injection. **b,** Images of GEMINI particles in the mouse brain with two timestamps applied. Inset: magnified image of a GEMINI particle. Scale bars: 50/2 μm for low/high magnification. **c,** Mean fluorescence profiles of stamps #1 and #2 (n = 36, from 3 mice). **d,** Experimental procedure of *in vivo* recording of DOX-induced expression. **e,** Images (*left*) and fluorescence profiles (*right*) of GEMINI particles that encode DOX-mediated expression signal induced at 0 (*top*), 12 (*middle*), and 24 h (*bottom*). Scale bars: 2 μm. **f,** Normalized radial positions of half-maximum intensity in individual GEMINI particles that encode signals induced at 0, 12, 24 h (n= 179/150/187 for 0/12/24 h, 3 mice per group). **g,** Experimental procedure of *in vivo* recording of seizure-induced *cfos* activation. **h,** Images (*left*) and the fluorescence profiles (*right*) of GEMINI particles that encode *cfos* activation signal induced at 0 (*top*) and 24 h (*bottom*). Scale bars: 2 μm. **i,** Normalized radial positions of half-maximum intensity in individual GEMINI particles that encode signals induced at 0 and 24 h (n= 111/80 for 0/24 h, 3 mice per group). **f,i,** Box bounds: 25th and 75th percentile; whiskers: minimum and maximum; and center lines: median.

We then recorded transcriptional histories in the brain. We first tracked DOX-mediated transcription using the Tet-ON system, where a second AAV encoding constitutively expressed rtTA3G and TRE-promoter-driven reporter subunit was co-injected at a dose 10% of total AAV (**Fig. S14a**). At 7 days post-injection, the first timestamp was applied (JF_669_), followed by i.p. administration of 40 mg kg^-1^ DOX at 0, 12, and 24 hours after the first stamp. The second stamp was introduced at 24 hours using JF_552_ (**Fig. 6d**).

We first observed the relative positions of signal bands to the timestamps, where a later induction resulted in an outward shift of signal bands (**Fig. 6e**). Normalized mean profiles revealed that band positions coincided with the timing of DOX induction, indicating that temporal information of distinct stimuli was captured (**Fig. S14c**). It is noted that the signal peak exhibited a considerable delay, as DOX persisted in the tissue and kept the signal ON for an extended period^39^. We then analyzed signals recorded in individual GEMINI particles and statistically compared the band positions (**Fig. 6f**). Significant differences were observed among the three groups, indicating that signals with distinct onsets were distinguishable. It is noted that the recorded signals exhibited a much broader distribution than recordings in the culture or xenograft model, which could reflect greater cellular heterogeneity in nature tissue and variations in the pharmacokinetics of HTL dyes and DOX across the blood-brain barrier. The asynchronous nucleation and different growth rates (**Extended Data Fig. 7c**), caused by the varied multiplicity of infection (MOI) in local AAV delivery, could also play a role.

We then asked if GEMINI could temporally resolve the transcription of immediate early genes (IEGs), like *cfos*, that are regulated by neural activity^40^. To test this, we constructed a signal transduction circuit jointly controlled by neural activity and DOX (**Fig. S14b**). Recording was initiated by 40 mg kg^-1^ DOX (i.p.), and neurons were activated at various times by 50 mg kg^-1^ (i.p.) pentylenetetrazole (PTZ), a GABA-A receptor antagonist that induces seizures in multiple brain regions^41^. Two timestamps were applied with a 24-hour interval for temporal decoding (**Fig. 6g**). The individual particles (**Fig. 6h**) and the mean fluorescence profiles (**Fig. S14d**) from the 0-hour and 24-hour groups showed expected band positions corresponding to the timing of seizure induction. Statistical analysis revealed significance between the two groups, demonstrating the recording of temporal information. We further examined if the seizure induction could be recorded by driving the expression of reporter subunit directly using the *cfos* promoter. To facilitate comparisons, the genes for direct and DOX-mediated signal reporting were co-expressed for GEMINI recording (**Fig. S15a**). Seizure induction was accompanied by delivery of a timestamp to mark the timing of the stimulus (**Fig. S15b**). The seizure-induced activation was clearly resolvable in both methods, while notable background signals were found before seizure induction in the direct-reporting channel, likely due to spontaneous neural activities (**Fig. S15c**). While both methods are valuable, the DOX-mediated strategy is better suited for distinguishing activities of interest from historical events. Together, these studies demonstrate that GEMINI can be used to assess diverse activity-induced transcriptional events in the native tissue.

## Discussion

*In cellulo* recording using engineered protein assemblies provides new opportunities for resolving cell dynamics within diverse tissues. Here, we exploited computational design and multi-level screening to expand the toolbox of intracellular assemblies for GEMINI development. The obtained scaffold is relatively compact, with its subunits (GEMINI-A/B: 19.0/13.4 kDa) comparable in size or even smaller than several genetically encodable reporters for cellular history^6,7^. The methodology also holds the promise to create more assemblies for *in cellulo* recording and beyond, unleashing great potential for future advancements.

Compared to linear recorders, GEMINI exhibits distinct advantages. First, it lowers the mechanical perturbation of cells. Linear recorders often stretch cells as they grow, potentially introducing artifacts through mechano-signaling^42^. In addition, the resultant cell deformation, while tolerable in culture, could be a major concern *in vivo*. Second, GEMINI particles write signals isotropically, allowing readout regardless of their spatial orientation. In contrast, the uniaxial elongation of linear recorders imposes significant challenges in retrieving signals from tilted recorders. Overall, isotropic recording is essential for minimal cellular perturbation and scalable signal retrieval, offering an attainable path to *in vivo* organ-wide recording at the cellular level.

GEMINI exhibits predictable growth necessary for temporal recording. Using our model, the growth profile of GEMINI can be precisely resolved with as few as two timestamps. Exploiting this capability, artificial signals were temporally decoded with mean errors generally under one hour for individual particles. Even high temporal resolution is achievable if more timestamps are employed as references for decoding.

To map physiologically relevant signals, we utilized transcriptional dynamics as a proxy for their corresponding cascades. This modality has been validated and broadly employed in fluorescent and luciferase transcriptional reporters, readily transferrable to GEMINI in an plug-and-play manner^33,34^. A potential limitation involves the noticeable delay of signals with respect to the actual events. Nevertheless, the delays were predictable and thus could be systemically subtracted in data analysis. Protein-based GEMINI also permits the employment of other reporting mechanisms that act on proteins directly to eliminate delays^43^.

Moreover, as the signal readout is through fluorescence microscopy, there is a negligible barrier to utilizing GEMINI by most biological laboratories. It could serve as a powerful alternative to existing reporter assays without additional instrumentation while obtaining additional spatiotemporal details. Moreover, GEMINI could complement real-time fluorescent biosensors for time-resolved tracking of cell activities^3^, especially when hour-level temporal resolution is sufficient or real-time imaging is not accessible.

Besides its broad *in vitro* applications, GEMINI can provide valuable insight into intricate cell dynamics when deployed *in vivo*. Its genetically encodable nature, combined with imaging-based retrospective signal retrieval, represents a feasible route toward organ-wide cellular-level recording across space and time. Notably, the *in vivo* timestamping is crucial for *in cellulo* recording. This is particularly challenging in the brain, where the BBB limits the delivery of many potential timestamping molecules^44^. Our demonstration of systemic timestamping represents a significant advance in applying GEMINI *in vivo*.

Using a xenograft model, we showed GEMINI’s potential for tissue-wide, cellular level recording of signaling history. This study revealed pronounced cellular heterogeneity in NFκB activation in response to systemic inflammation and identified heterogeneous vascularization as an important contributing factor. This approach, combining engineered cells and *in vivo* transplantation, offers a promising platform for preclinical research, particularly in developing tumor models to map oncogenic signaling and evaluate therapeutic responses. Moreover, GEMINI exhibited low cytotoxicity in native tissues like mouse brain and can temporally resolve cellular activities such as DOX-induced transcription and activity-dependent *cfos* activation. It is noted that the temporal resolution of recording in the brain was lower than that observed in culture or xenografts, which could be attributed to factors such as the inherent heterogeneity of neurons, less accurate timestamping, variability in GEMINI expression, and less accurate predictive model for decoding. Disentangling these factors remains a challenge. Nevertheless, GEMINI holds great potential for further optimization.

Looking ahead, several advancements are desired to improve spatiotemporal mapping of cell dynamics within intact tissues. First, we need to implement GEMINI organ-wide, where systemic AAV delivery^45,46^ and transgenic mouse development^47^ are better for global coverage and homogenous expression than local AAV injection. Second, we need a high-throughput method to retrieve signals from individual cells across an organ. We believe the combination of tissue expansion^48^ and light-sheet microscopy (LSM)^49^ holds a great promise. We believe GEMINI can be expanded to a scale resolvable by LSM, similar other protein assemblies^50^. Third, decoding models must be modified for *in vivo* conditions, accounting for potential differences in the growth behavior within living tissues. However, real-time tracking remains challenging due to current limitations in intravital imaging and the constraints of prolonged observation under animal welfare guidelines. As a result, alternative strategies, such as multi-timestamping or statistical analysis at discrete time points, may be employed as a proxy. Fourth, we need an automated data analysis pipeline to resolve signal dynamics from individual GEMINI particles precisely. Classical computer vision and deep learning could be employed jointly to achieve this goal.

In addition, it would be valuable to achieve (1) recorders with a smaller subunit, possibly using single-component scaffolds, (2) multiplexed recording of diverse signaling pathways, (3) tracking of faster dynamics such as kinase activity and biomolecule concentrations, (4) further improved temporal resolution *in cellulo* and *in vivo*, and (5) *in vivo* implementation to other tissues, organs, and across various stages of development.

Finally, it is important to acknowledge that GEMINI, as a protein-based recorder, does not inherently support inheritance across cell divisions. In future applications, especially when intergenerational continuity is critical, integrating GEMINI with lineage tracing methods could allow reconstruction of lost historical information through clonal relationships.

We believe that continued optimization in these areas would help transform GEMINI into a platform broadly applicable to spatiotemporally resolve cellular histories in animals, enabling deeper insight into the mechanisms underlying health and disease.

## Methods

### Protein assembly design

The design of protein assemblies followed the method we reported recently^27^. Briefly, validated protein cages were docked to target the F4_1_32 and I432 space groups. Dockings were performed in Pymol by alignment and translations. For the docking of the F4_1_32 space group, two tetrahedral cages (TET-1/-2) were aligned with every C2 symmetry axis along the x, y, and z coordinate axes and center of mass at the origin of the coordinate. While keeping TET-1 fixed at the origin, TET-2 was rotated along the z-axis for 90° and translated along the [1, 1, −1] direction to dock with TET-1. The hexamer between the two cages with D3 dihedral symmetry was extracted and aligned with its C3 axis along the z-axis and its C2 axis along the x-axis and with its center of mass at the origin. I432 space group was docked between two octahedral cages (OCT-1/-2), where OCT-2 was translated only without any rotation. In both cases, the D3 assembly units contained the crystal contact and were used for the Rosetta sequence design. In the RosettaScripts framework, input monomers were symmetrized into D3 dihedrals, and then we sampled the interface distances by translation along the z-axis (dihedral axis) within ±1.5 Å of the docked conformation without a rotational DOF. For each translation, the dihedral interface was designed with rigid protein backbones by packing a rotamer with layer design restrictions at interacting residues. Then, designs were filtered by ddG <0, solvent-accessible surface area >200, clash check ≤2 and unsatisfied hydrogen bonds ≤2 before being visually inspected for hydrophobic packings. The schematics of protein structures were rendered using Protein Imager^51^.

### Protein expression and purification

Synthetic genes were purchased from IDT (Integrated DNA Technologies) as plasmids in pET29b vector with a hexahistidine affinity tag. Plasmids were cloned into BL21* (DE3) E. coli competent cells (Invitrogen). Single colonies from agar plate with 100 mg/L kanamycin were inoculated in 50 mL of Studier autoinduction media, and the expression continued at 37 °C for over 24 hours. The cells were harvested by centrifugation at 4000 g for 10 minutes, and resuspended in 35 mL lysis buffer of 300 mM NaCl, 25 mM Tris pH 8.0 and 1 mM PMSF. After lysis by sonication and centrifugation at 14000 g for 45 min, the supernatant was purified by Ni^2+^ immobilized metal affinity chromatography (IMAC) with Ni-NTA Superflow resins (Qiagen). Resins with bound cell lysate were washed with 10 mL (bed volume 1 mL) of washing buffer (300 mM NaCl, 25 mM Tris pH 8.0, 60 mM imidazole) and eluted with 5 mL of elution buffer (300 mM NaCl, 25 mM Tris pH 8.0, 300 mM imidazole). Eluted proteins were further purified by size exclusion chromatography (SEC) in 150 mM NaCl, 25 mM Tris pH 8.0 on a Superose 6 Increase 10/300 gel filtration column (Cytiva), before concentrated by 10K concentrators (Amicon).

### Library generation of mutated protein assembly

For structurally validated protein assemblies, their computational design models and X-ray models were further used for more extensive cycles of Rosetta sequence design, where diverse, mutated sequences were generated from fixed interface backbones. The new design models were visually inspected for their differences in the side-chain interactions from the validated designs. Alternative sequences near the core of the interfaces were selected either individually or as groups to build mutation libraries for *in vitro* screening of GEMINI scaffolds.

### *In vitro* protein assembly screening

Protein assembly subunits with Rosetta predicted mutations were screened by batch crystallization experiments *in vitro*, aiming at assembly conditions with ionic strength lower than 500 mM NaCl. Equimolar ratios of protein subunits were mixed at 5-50 μM concentration in 150 mM, 300 mM, and 500 mM NaCl, 25 mM Tris pH 8.0. Crystallization results were inspected by optical microscopy imaging for up to two weeks.

### Cloning and molecular biology

GEMINI constructs were cloned into a PiggyBac plasmid backbone (Addgene #104454) for *in vitro* study and clonal cell line development, and into an AAV plasmid backbone (Addgene #100854) for AAV packaging and *in vivo* study. Plasmids were constructed using standard Gibson assembly. In brief, the vector was linearized by double restriction enzyme digestion and purified by the GeneJET gel extraction kit (Thermo Fisher Scientific). DNA fragments were obtained by polymerase chain reaction (PCR) amplification and then combined with the linearized backbones by Gibson ligation. Genes for the protein assembly variants were obtained by DNA synthesis (Twist Bioscience). The *cfos* promoter was cloned from Addgene #47907; the NFκB reporter promoter was cloned from Addgene #82024; rtTA3G was cloned from Addgene #120309; HT was cloned from #177882; GFP was cloned from Addgene #177881; mCherry was cloned from Addgene #81041. A 6×GGS linker was employed to connect the HT or FPs with the assembly subunits. All plasmids were verified by whole-plasmid sequencing or Sanger sequencing around the cloned regions. The key plasmids used in this work were listed in **Table S1** and are available on Addgene. The amino-acid sequences of protein variants we developed to assemble *in cellulo* were listed in **Table S2**.

### Cell culture and transfection

HEK293T and other mammalian cells (gifts from Dr. Inoue Takanari and Dr. Konstantinos Konstantopoulos, originally acquired from the American Type Culture Collection) were grown and passed following standard protocols as described previously^24^. Cells at low passage number (<15 passages) were seeded at a confluence of 30% onto 35-100 mm dishes coated with gelatin (STEMCELL Technologies) or 14-mm glass-bottom dishes (CellVis, D35-14-1.5-N) coated with 0.1 mg mL^-1^ poly-D-Lysine (P6407, Sigma-Aldrich). Dulbecco’s modified Eagle medium (DMEM, Corning) supplemented with 10% FBS (VWR) and 1% penicillin–streptomycin (Millipore Sigma) was used as the culture medium. All cells were maintained at 37 °C in a humidified atmosphere of 5% CO_2_. When cells reached 50–70% confluence, genes were delivered using 25K linear polyethylenimine (PEI, Polysciences) following the protocol reported previously^52^ or Lipofectamine 3000 transfection kit (Thermo). The cells were then further incubated at 37 °C and 5% CO_2_ for ∼8-24 hours before experiments.

### Clonal cell line development

The clonal HEK cell lines with stable integration of the gene of interest were generated using the PiggyBac transposon system^53^. Briefly, HEK293T cells were co-transfected with the PiggyBac transposon plasmids carrying the gene of interest and a puromycin-resistant gene, and a Super PiggyBac transposase expression vector (NovoPro). 1mg mL^-1^ PEI was complexed with the plasmids for transfection. At 48 hours after transfection, cells were selected using 2 ug mL^-1^ puromycin (Gibco, A1113803). Selection was maintained for 1–2 weeks to ensure stable integration. Clonal cell lines were then established by limiting dilution, where single cells were plated into 96-well plates and allowed to expand. Clones were screened to confirm the presence and expression of the transgene. Positive clones were further functionally validated, and the selected colony was expanded, maintained under selection conditions, and cryopreserved for future use.

### AAV production and purification

AAVs were produced and purified following the method reported previously^45^. Briefly, HEK293T cells were transfected with the AAV cargo plasmid of interest, helper plasmid, and the AAV-PHP.eB rep-cap plasmid (Addgene #103005). Cells were incubated for 48– 72 hours post-transfection before harvesting. Viral particles were isolated from the cell pellet and supernatant via freeze-thaw lysis or SAN digestion and clarified by centrifugation. Purification of AAV was carried out using iodixanol gradient ultracentrifugation to ensure the purity of viral particles. Viral titers were determined through quantitative PCR (qPCR). The AAVs were concentrated into a titer of 1×10^14^ vg ml^-1^ before use. The final AAV preparations were stored at 4 °C for short-term use (< 1 month) or -80°C for long-term storage.

### Confocal imaging

Confocal imaging was performed using Zeiss LSM780, LSM800, or LSM980 microscopes. Lambda scan mode (LSM780, LSM980) was used to image GEMINI particles with multi-color labeling. The excitation laser wavelengths for various fluorescent proteins or dyes were: 405 nm (Hoechst 33342), 488 nm (GFP, YFP, JF_525_), 561 nm (mCherry, JF_552_, JF_608_), 633 nm (JF_669_). Single crystals were acquired with x63 oil immersion objectives. In each Lambda scan, a 32-channel QUASAR detector was utilized to simultaneously acquire a hyper-spectral stack of images with distinct collection wavelengths in the range of 410 nm - 695 nm. The multispectral images were then unmixed with the built-in linear unmixing algorithm in ZEISS Zen Blue software. Images of individual fluorescent labels were acquired in the same instrumental configuration as the references for linear unmixing. The quality of spectral unmixing was assessed by examining the residual signals after unmixing.

### Time-lapse imaging

Cells expressing the target constructs were grown on glass-bottom culture dishes (CellVis) and imaged under a Zeiss Axio Observer wide-field epifluorescence microscope at 37 °C and 5% CO_2_. Focus across frames was automatically calibrated by a Definite Focus 3 module.

### Vertebrate animals

Adult (4-8 weeks) female athymic nude mice J:NU (Jackson Laboratory) were used in the transplantation of HEK293T xenografts. Adult (5-10 weeks) CD1 (Charles River Laboratories) or C57BL/6 (Jackson Laboratory) mice were used throughout the other study. The mice were housed at 22 ± 1 °C with humidity ranging from 30% to 70% and on a 12-hour light/dark cycle. All animal experiments were conducted in strict accordance with the guidelines and regulations set forth by the Johns Hopkins University Animal Care and Use Committee (ACUC). Experimental protocols were reviewed and approved by the ACUC under protocol number MO21E409 and MO24E325.

### Stereotaxic surgeries

Prior to surgery, animals were anesthetized with 4% isoflurane in air (induction) and maintained at 1-2% isoflurane in air throughout the procedure. Carprofen (5 mg kg^-1^, s.c.) was administered as preoperative analgesia. During surgery, animal body temperature was monitored and maintained at 37°C using a heating pad. Once anesthetized, mice were secured in a stereotaxic frame (Stoelting), and bilateral openings (1 mm in diameter) were made on the skull at the corresponding coordinates. A Nanoliter2020 Injector (WPI) was utilized to inject AAVs at a rate of 100 nL min^-1^. Without special note, 100 nL AAV was injected into each position. After each injection, the micro-capillary needle was held in place for 3 minutes to allow for the diffusion of AAVs into the tissue before moving to the next position. When the injection was completed, the scalp was sealed with Vetbond (3M), and mice were continuously monitored till recovery. The following coordinates were used to target specific brain regions (relative to bregma). Primary motor cortex: AP: +1.0 and +1.4, ML: ±1.5, DV: -0.3 to -1.0 (0.1 increments); Dorsal hippocampus: AP: -1.8 and -2.2, ML: ±1.5, DV: -1.2 to -1.8 (0.1 increments); Ventral hippocampus: AP: -2.8 and -3.2, ML: ±2.8, DV: -2.8 to -4.0 (0.1 increments).

### *In vivo* timestamping via retro-orbital dye injection

*In vivo* timestamps were applied via retro-orbital injection of blood-brain-barrier permeable HTL dyes. Briefly, 100 nmol of a HTL dye (JF_669_ or JF_552_, gift from the Lavis Lab, HHMI) was first dissolved in 100 μL DMSO with 10% Pluronic F127. 400 μL sterile saline was further added to yield the HTL solution with a final concentration of 200 μM. Before injection, the mouse was anaestheized with isoflurance (3% induction and 1.5% during injection). The level of anesthesia was assessed by toe pinch. The mouse is then positioned, and the eyelid of the target eye is gently retracted to expose the globe. A syringe with 31G needle containing 100 μL of the HTL dye solution was carefully inserted into the retro-orbital space with the bevel down, ensuring the needle tip is within the venous sinus. The dye solution was then injected steadily and slowly to avoid damaging the eye and the surrounding tissues. After injection, the mouse was allowed to recover from anesthesia in the home cage, with close monitoring for any signs of discomfort or complications.

### HEK293T Xenograft model

For the *in vivo* NFkB-GEMINI xenograft model, the NFkB-GEMINI clonal cells were resuspended by PBS and mixed with Matrigel® (Corning, 356231) in a 1:2 volume ratio. 5×10^6^ cells were planted subcutaneously in the flank regions of J:NU athymic nude mice. After implantation, tumor size was monitored and all the mice were randomly assigned into different groups. *In vivo* crystal growth was initiated by 40 mg kg^-1^ DOX administrated via i.p. injection every 2 days starting from 10 days after implantation. LPS (Sigma, 437627) was administered via i.p. injection at designated time points.

### Open-field tests

The open-field test was conducted in a 38 x 38 cm arena with 38 cm high walls. Mice were acclimated to the testing room for 1 hour prior to the test. When the test began, each mouse was placed in the center of the arena and allowed to explore for 7 minutes. Animal traveling within the arena was video-recorded. The arena was cleaned with 70% ethanol three times between sessions to eliminate scent cues. Data was analyzed offline with in-house Python codes, where the total traveling distance and average traveling speed were analyzed as indicators of the motor function. All procedures were in accordance with institutional and national animal care guidelines.

### Y-maze tests

The Y-maze test was conducted using a Y-shaped maze with three arms (each 35 cm long, 5 cm wide, with 10 cm high walls). Mice were acclimated to the testing room for 1 hour before the test. Each mouse was placed at the end of the initial arm and allowed to explore the maze for 10 minutes with one arm (the novel arm) blocked. The mouse was returned to the home cage for 20 minutes before re-entering the maze for 5 minutes, when all arms were accessible. The second entrance was video-recorded. The maze was cleaned with 70% ethanol three times between sessions to eliminate scent cues. Data was analyzed offline with in-house Python codes. The duration the mouse spent in the novel arm versus the alternative arm was analyzed. All procedures adhered to institutional and national guidelines for animal care and use.

### Seizure model

Pentylenetetrazole (PTZ, Cayman, 18682) was utilized to induce seizure in mice. PTZ was first dissolved in sterile saline at 9 mg mL^-1^, which was delivered to mice via i.p. injection at a dose of 50 mg kg^-1^. The behaviour of mice was carefully monitored for 1 hour post-seizure induction. The level of seizure was assessed by the Racine scale^54^, where only mice showed a seizure level of 5 or higher were utilized in further study.

### Immunohistochemistry

To prepare the brain samples for immunohistochemistry, mice were first euthanized with CO_2_ flow of 50% of cage volume per minute for 10 minutes, and transcardially perfused with 40 mL of ice-cold phosphate-buffered saline (PBS), followed by 40 mL of 4% paraformaldehyde (PFA) in PBS. After decapitation, the brain was carefully extracted and post-fixed in 4% PFA overnight at 4°C. The brain was then transferred to a 30% sucrose solution in PBS for cryoprotection and stored until fully equilibrated (sank to the bottom of the tube). Once the brain was fully equilibrated, they were embedded in Optimal Cutting Temperature (O.C.T.) compound (Tissue-Tek) and frozen on dry ice. Coronal sections, 50 μm thick, were prepared using a cyro-sectioning machine (Leica 1850 CM). For immunohistochemistry, brain sections were first washed three times with PBS to remove any residual O.C.T. compound. The sections were then blocked in PBS containing 0.3% Triton X-100 and 5% normal donkey serum (Abcam, ab7475) for 1 hour at room temperature and washed three times with PBS. The sections were then incubated with primary antibodies anti-NeuN (GeneTex, GTX132974) and anti-glial fibrillary acidic protein (anti-GFAP, Cell Signaling, #3670) overnight at 4°C with gentle shaking. The sections were then washed for three times with PBS and incubated for 2 hours at room temperature in the dark with corresponding secondary antibodies (Invitrogen). After additional washes with PBS three times, the brain sections were stained with Hochest 33342 (Invitrogen, H3570) for 30 minutes to mark all cell nuclei. The sections were mounted on glass slides with #1.5 coverslips using Aqua-Poly/Mount (Polysciences, 18606) mounting media. The samples were stored in the dark at room temperature for 4-8 hours to allow complete permeation of the media before imaging.

For immunohistochemistry staining of xenografts, tumors was sliced into 150μm thickness instead. Anti-CD31 antibody (BD Biosciences, 550274) was used as primary antibody and Donkey anti-Rat AF647 antibody (Invitrogen, A48272) was used as secondary antibody.

### Image processing and data analysis

Python, MATLAB, ImageJ, Vision4D (arivis), and napari were used for image processing and/or visualization. Fluorescence profiles were extracted along the radial directions of GEMINI particles, perpendicular to one of the edges. The profiles were averaged over a certain width along the radial axis to reduce the noise. A low-pass filter (Butterworth) might be applied to further denoise the profiles. Baseline subtraction might be applied to remove the background intensity outside of the band regions. The profiles before and after filtering were compared to ensure no distortion to the curves. The onset of a fluorescence band was determined by the intercept of the tangent line at 50% peak intensity with the baseline. The locations of the peaks were determined by the intensity maxima.

To track GEMINI growth, we first employed adaptive thresholding and Sobel filters to enhance GEMINI boundaries. After distance transform, GEMINI centers were located by a maxima-finding algorithm, while radius estimation based on intensity cutoffs and background thresholds helps delineate particle regions. In the tracking phase, we employed a linear programming-based approach to connect crystals across video frames. The algorithm detected particles frame-by-frame in reverse order, starting from the last frame, and formed tracks for each crystal based on their position, size, and intensity. Using features like centroid coordinates and area, the algorithm calculated distances between particles in consecutive frames and minimized a cost function through linear sum assignment to optimally match particles. Area and intensity differences are factored into the cost to ensure robust tracking. The tracking results were then examined, and those showing errors in tracking were excluded from further analysis. Overall, the method achieved accurate and consistent tracking for most particles. As the algorithm measured the area of the 2D GEMINI projection *A* (in pixels), we quantified the volume of the GEMINI particles by a volume index defined as *A*^3/2^. The index was normalized for comparing the growth rate of different particles.

### Statistics and reproducibility

Sample sizes are specified in the figure legends. Differences with P < 0.05 were considered statistically significant. For the statistical comparison of the two groups, two-sided t-tests were applied. All fluorescence images presented in the main text, extended data, and supplementary figures are representative images from at least three independent repeats.

## Data availability

The plasmids constructed and used in this work are deposited to Addgene. Data reported in this article is available at GitHub (https://github.com/DCLinLab/GEMINI). Other data and materials comprising raw images, raw videos, and clonal cell lines is available from the corresponding author upon reasonable request.

## Code availability

Custom data analysis code developed and used for this project is available at GitHub (https://github.com/DCLinLab/GEMINI).

## Acknowledgments

We thank Dr. Luke D. Lavis and his team for sharing HTL dyes. We thank Dr. Takanari Inoue and Dr. Konstantinos Konstantopoulos for sharing mammalian cell lines used in Fig. S1. This work is supported by National Institutes of Health (NIH) National Institute of General Medical Sciences (NIGMS) grant R35GM147274 (D.L., Y.Y., J.L., Y.W.), Department of Defense (DoD) Air Force Office of Scientific Research (AFOSR) Young Investigator Program (YIP) grant FA9550-23-1-0174 (D.L., Y.W., H.Y.), NIH BRAIN Initiative grant R21EY035955 (D.L., Y.S.), David and Lucile Packard Foundation (D.L.), Kavli Neuroscience Discovery Institute Distinguished Graduate Fellowship (Y.Y.), the Howard Hughes Medical Institute (D.B.), the Audacious Project at the Institute for Protein Design (Z.L., S.W., and D.B.), Beckman Institute CLOVER Center at Caltech (T.S.).

## Author contributions

D.L. conceived the project. D.L., Y.Y., and J.L. designed the experiments. Z.L., S.W. and D.B. designed the protein assemblies. Z.L. and S.W. performed *in vitro* screening of the assemblies. Y.Y. performed *in cellulo* screening of the assemblies. Y.Y. and J.L. designed and cloned the plasmids. J.L. and Y.Y. performed the confocal and time-lapse imaging. J.L and Y.Y. developed the clonal cell lines. Y.Y., J.L., Y.W., Y.S., A.Q. performed other characterizations in cultured cells and acquired the imaging data. T.S., Y.L., and Y.Y. produced and purified AAVs. Y.Y. and D.L. performed the *in vivo* experiments in the mouse brain and acquired the data. J.L. performed the *in vivo* xenograft study and acquired the data. Y.Y., W.C., J.L. and D.E.B. collected and analyzed the immunohistochemistry data. P.P., W.C., and Y.Y. performed *in vivo* calcium imaing and analysis, Y.Y., D.L., Z.Z.,and J.L. analyzed data and prepared figures. D.L. and Y.Y. wrote the paper. All authors participated in the revision of the manuscript.

---

**Extended Data Fig. 1.**
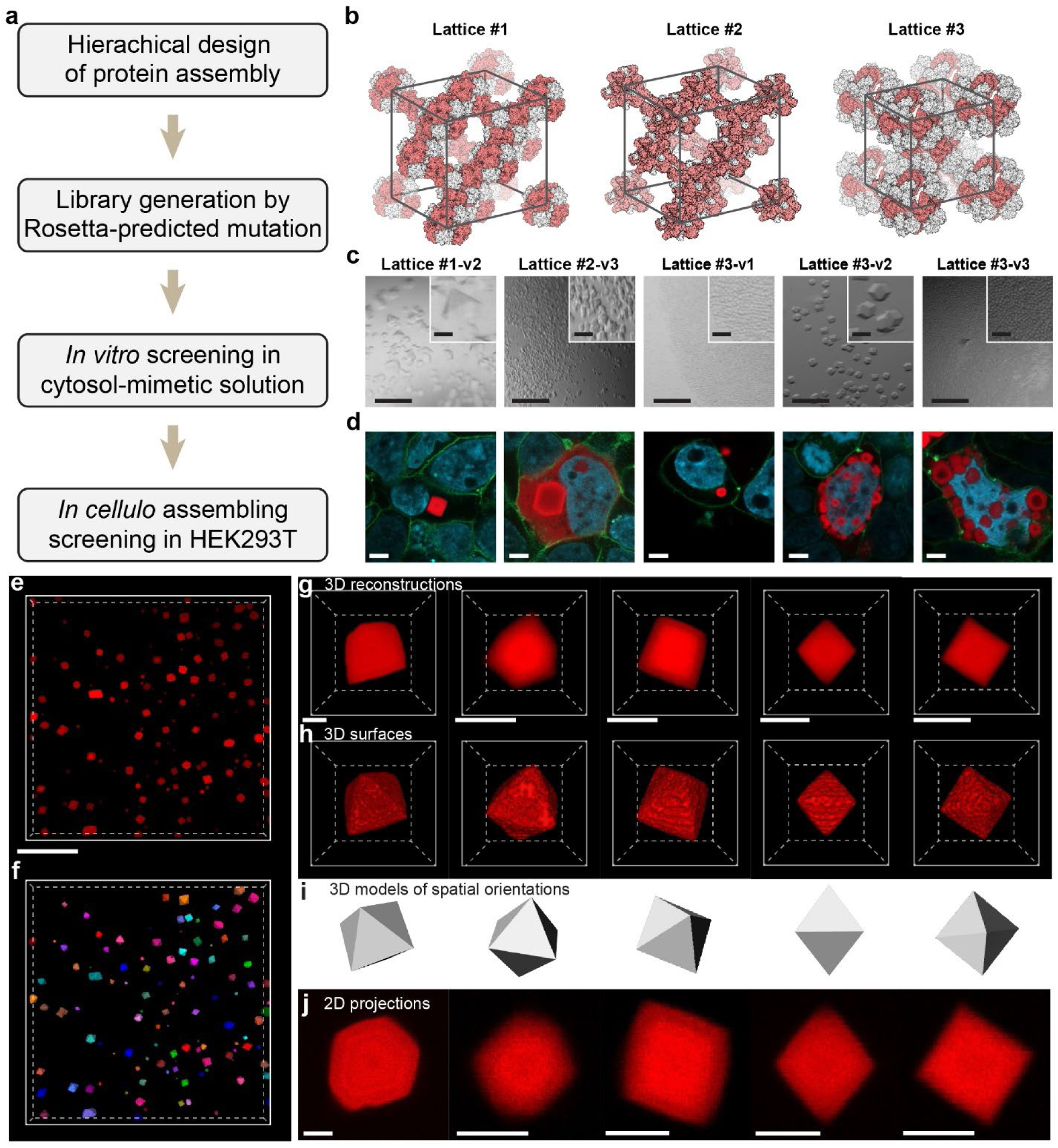
Development of new intracellular protein assemblies and GEMINI morphology. **a,** Experimental procedure to design and select for intracellular protein assemblies suitable as the GEMINI scaffold. **b,** Lattice structures of three variants involved in this study. **c,** Images of *in vitro* assembly of the variants in an ion strength comparable to the cytoplasm (100-500 mM NaCl). Scale bars: 200 μm; insets: 40 μm. **d,** Images showing the assembly of variants that pass the *in vitro* screening, where all the variants can assemble in live HEK293T cells. Scale bars: 5 μm. **e,f,** Low-magnification 3D image of GEMINI particles grown in live HEK293T cells (**e**) and the single-particle segmentation (**f**). **g-j,** 3D-reconstruction (**g**), 3D surfaces (**h**), 3D models (**i**), and 2D projections (**j**) of individual GEMINI particles that exhibit different sizes and spatial orientations. The particles exhibit consistent octahedral shapes. Scale bars: **e,** 50 μm; **g,h,j,** 2 μm.

**Expanded Data Fig. 2.**
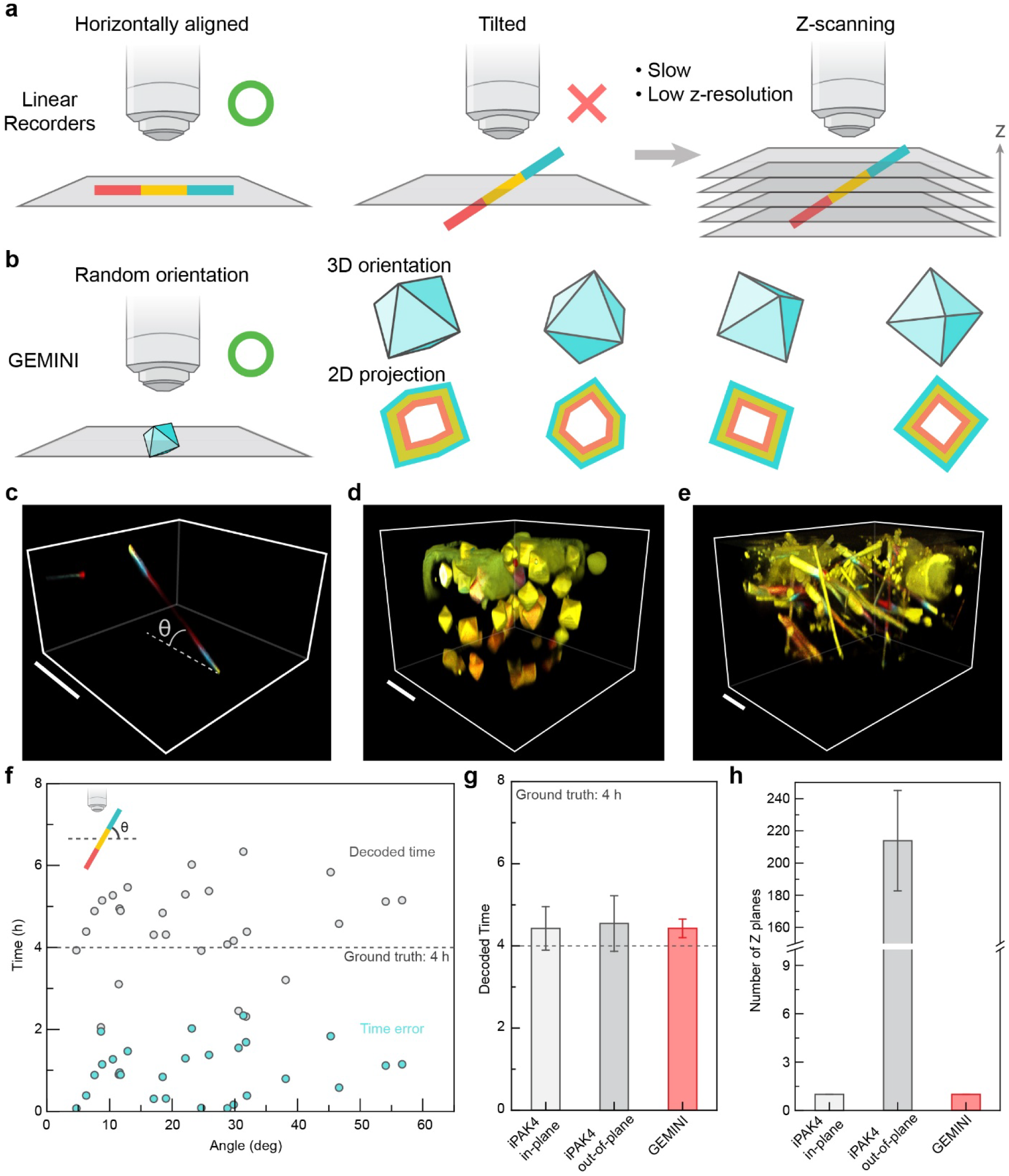
Comparison of linear recorders and GEMINI in signal retrieval. **a,** Schematic of imaging linear recorders by fluorescence microscopy. When the recorder happens to align horizontally, they can be imaged at high resolution. However, for most recorders that are tilted out-of-plane, volumetric imaging is needed to retrieve the recorded patterns, which is slow and exhibits much lower resolution. **b,** Schematic of imaging GEMINI by fluorescence microscopy. High-resolution fluorescence patterns are retrievable regardless of the spatial orientation of GEMINI particles, allowing fast signal retrieval from a large population of cells. **c,** A linear recorder tilted at an angle in culture. Scale bar: 50 μm. **d,e,** 3D cell culture expressing GEMINI (**d**) and iPAK4 linear recorder (**e**). Scale bars: 20 μm. **f,** The decoded time and error of iPAK4 linear recorder as a function of the tilted angle. Timestamps were applied at 0 and 8 hours, while the artificial signal was introduced at 4 hours. **g,** Comparison of the decoded time from in-plane iPak4, out-of-plane iPak4, and GEMINI. **h,** Average number of z-planes needed to obtain complete fluorescent profiles from individual recorders. Whiskers: s.d..

**Extended Data Fig. 3.**
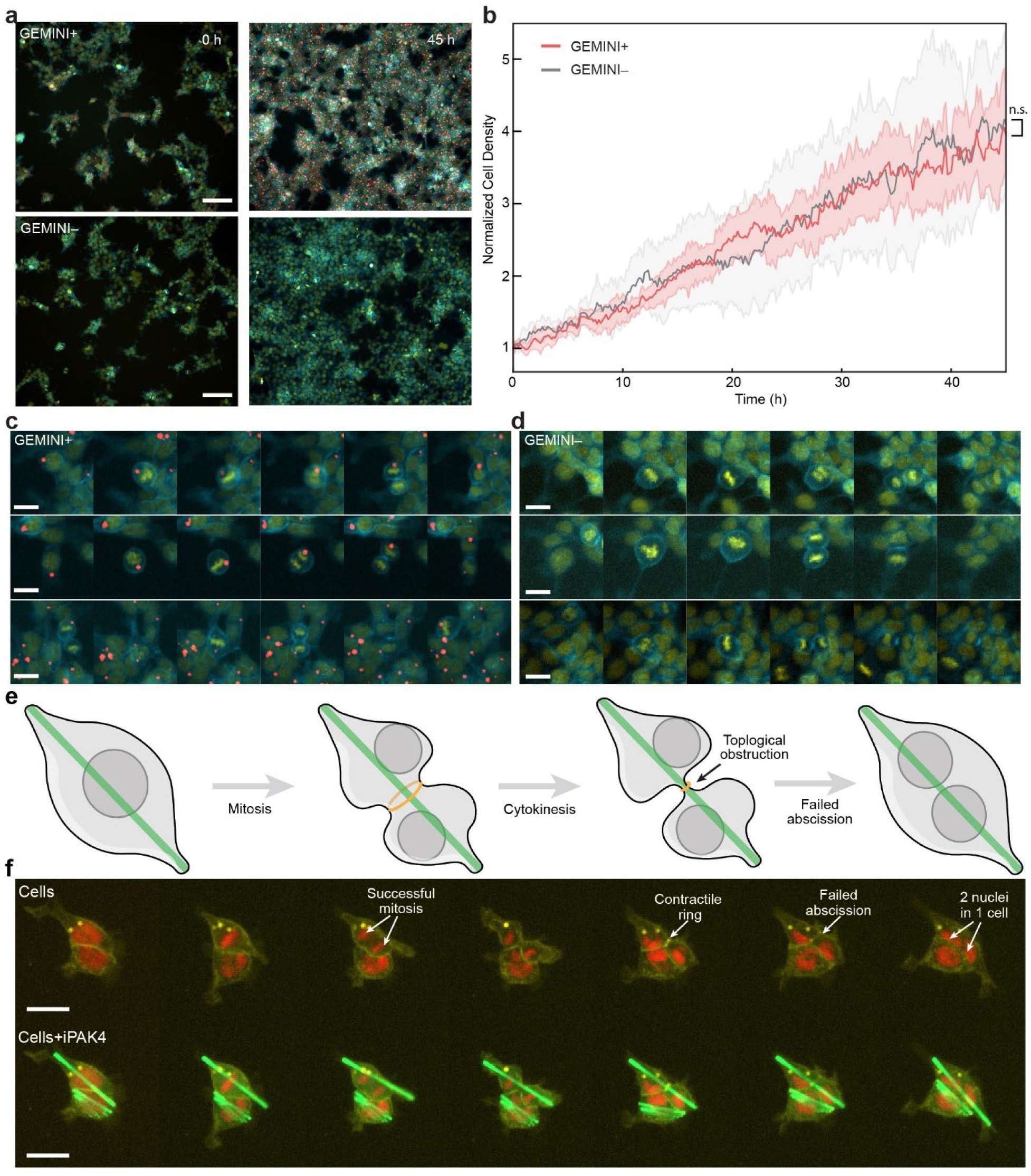
Impact of GEMINI and iPAK4 linear recorder on cell proliferation and division. **a,** Images comparing the cell density with (top) and without (bottom) GEMINI growth at t = 0 h (left) and 45 h (right). Scale bars: 100 μm. **b,** Comparison of the cell proliferation rate between the GEMINI+ and – groups. The two groups show a comparable proliferation rate. **c,d,** Snapshots of the cell division processes of live cells with (**c**) and without (**d**) intracellular GEMINI nucleation, showing GEMINI nucleation has negligible impact on the mitosis and cytokinesis. **e,** Schematic showing the topological constraint imposed by intracellular linear recorders that obstruct the abscission during cell division. **f,** Snapshots of HEK293T cells with an intracellular linear recorder during the division processes. Though mitosis was successful, the contractile ring failed to split the cell into two daughter cells and, therefore, resulted in one cell with two nuclei after division. Scale bars: 20 μm.

**Extended Data Fig. 4.**
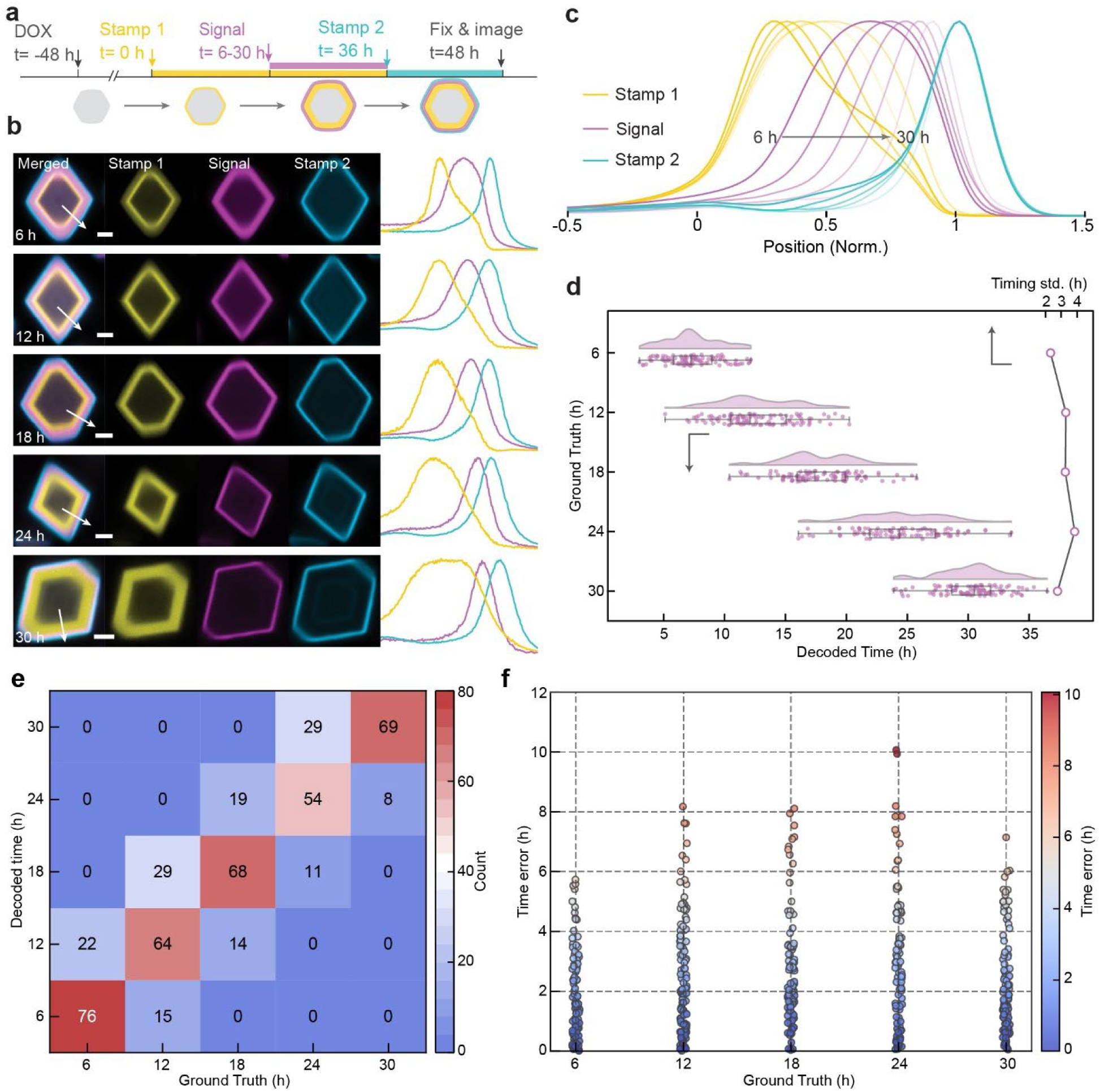
Temporal resolution of recording at the later stage of GEMINI growth. **a,** Experimental procedure of recording at a later stage of GEMINI growth, where the recording did not start until 48 h after the induction of GEMINI expression (denoted as 0 h), when GEMINI particles are much larger in size. The signal (blue) was induced at t = 6/12/18/24/30 h within a 36 h window defined by stamps #1 (red) and #2 (yellow). The cells were fixed at 48 h. **b**, Images (left) and fluorescence profiles (right) of GEMINI particles with signals introduced at t= 6-30 h. Scale bars: 2 μm. **c,** Mean fluorescence profiles of the timestamps and signal channels. **d,** Decoded onsets from individual GEMINI particles (top-x, n= 100/108/101/96/99 for t= 6/12/18/24/30 h) and the standard deviations (bottom-x) for each group. **e,** Confusion matrix comparing the decoded time (y-axis) to the ground truth (x-axis) for GEMINI particles in **d**. The color scale indicates the number of particles decoded to each time, highlighting accurate versus misassigned time points. **f,** Time error distribution for individual GEMINI particles in **d** plotted against their ground truths (x-axis). Each dot represents a single particle, with color indicating the magnitude of time decoding error (color bar, *right*). Box bounds: 25th and 75th percentile; whiskers: minimum and maximum; and center lines: median.

**Extended Data Fig. 5.**
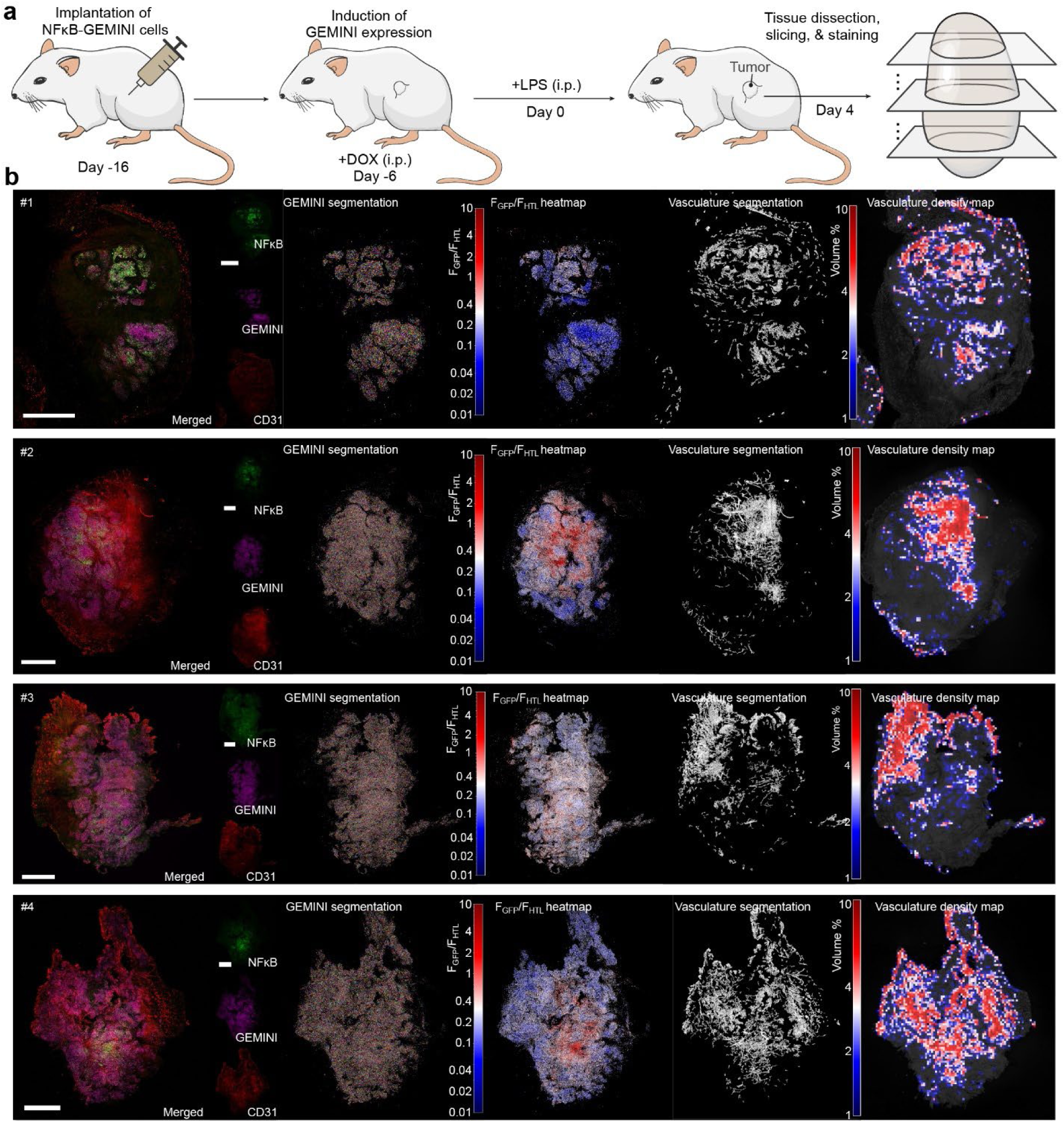
GEMINI expression across a tumor xenograft. **a,** Experimental procedure for recording inflammation-induced NFκB activity in HEK293T xenografts using GEMINI. **b,** Tissue slices from a xenograft following i.p. injection of 3 mg kg^−^¹ LPS. GEMINI particles were segmented, and NFκB signal intensity (GFP), normalized to HTL staining, was visualized as heatmaps across the tissue. Vasculature was labeled via anti-CD31 staining, segmented, and displayed as a vascular density heatmap. Scale bars: 1 mm.

**Extended Data Fig. 6.**
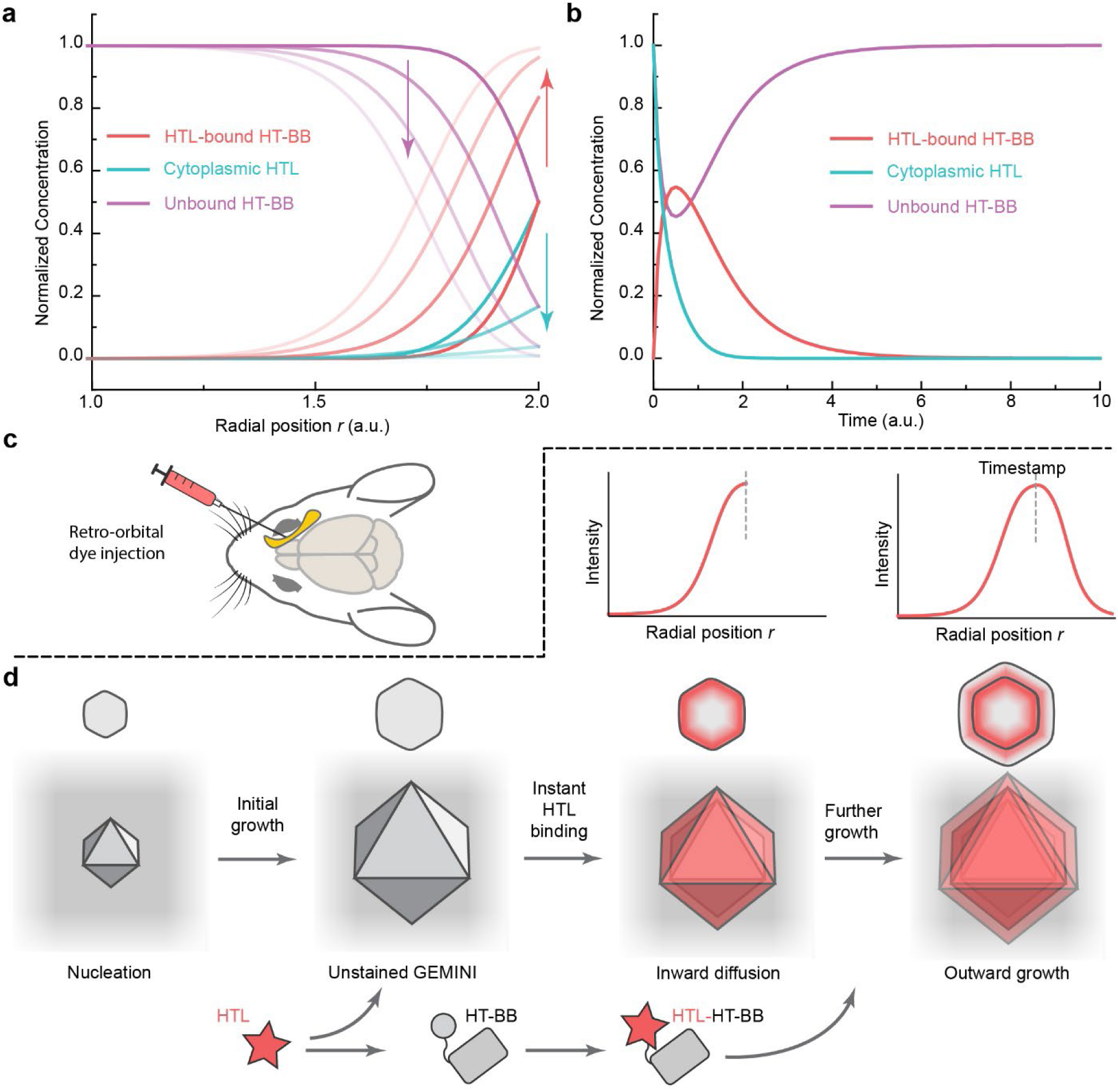
In vivo timestamping. **a,b** Numerical simulation of the HTL diffusion into existing GEMINI and bind with HT (**a**), and binding with cytoplasmic HT that will further grow onto GEMINI (**b**). **c,** Schematic illustration of systemic injection of HTL dyes into the mouse retro-orbital sinus. **d,** Schematic illustration of the HT labeling via systemic delivery, where we reasoned that the peak of the fluorescence band is a good proxy for the time of HTL injection.

**Extended Data Fig. 7.**
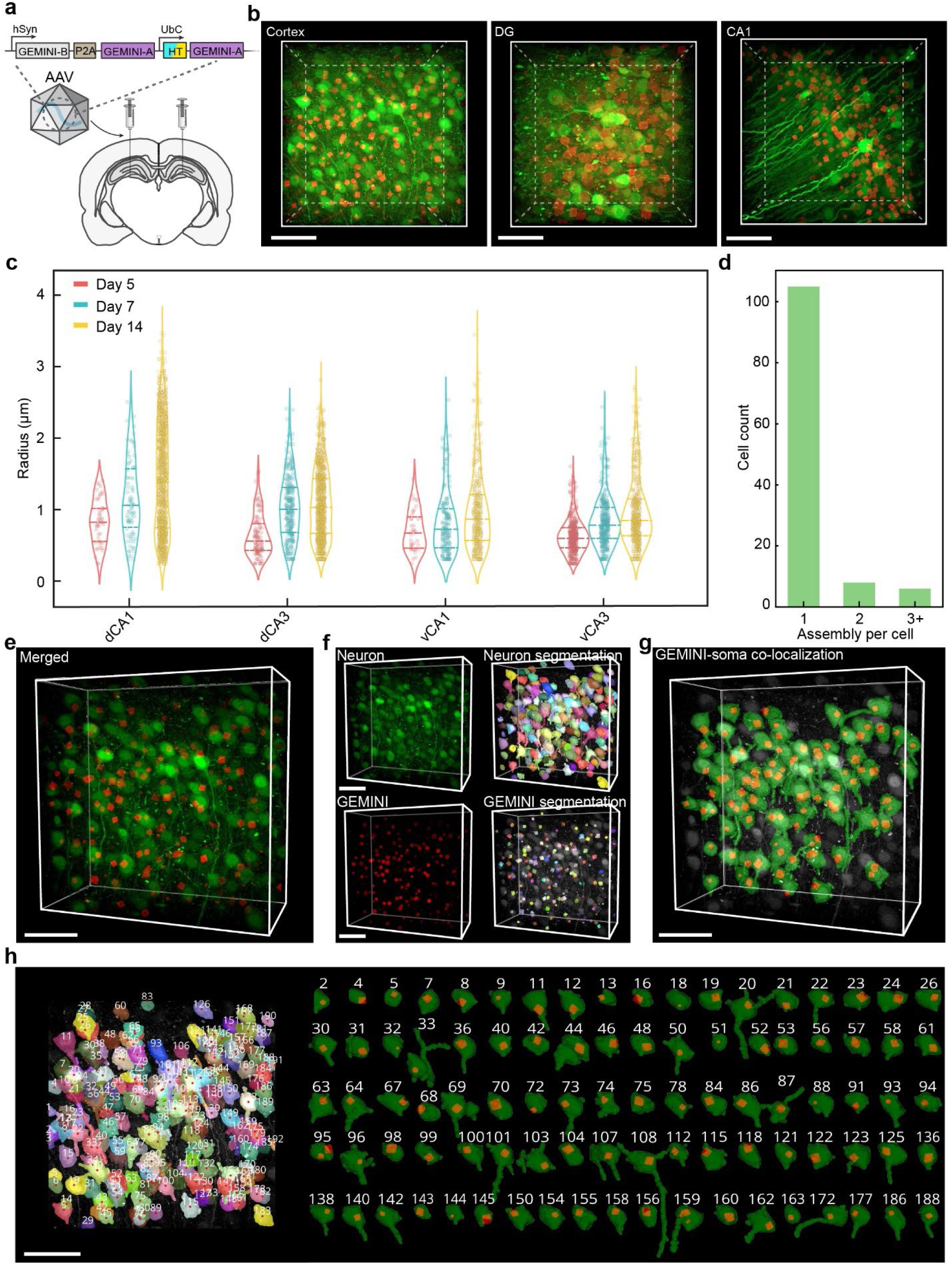
Implementation of GEMINI to the brain and cellular registration. **a,** Schematic showing the DNA construct for AAV packaging and the intracranial injection. **b,** 3D images showing GEMINI expression in various brain regions. Scale bar: 50 μm. **c,** Statistic analysis of the GEMINI particle size in various hippocampal regions at days 5, 7, and 14 (n = 2/3/5 mice for days 5/7/14). **d,** Number of GEMINI particles found per cell. Most cells only nucleate one GEMINI particle. **e,** 3D image of brain tissue expressing GEMINI. **f,** Segmentation of neurons (*top*) and GEMINI particles (*bottom*). **g,** Spatial colocalization of neurons and GEMINI within the tissue. **h,** Spatial labeling of individual neurons within the tissue (*left*) and the registration of each GEMINI particle to the soma of corresponding neurons (*right*). Each neuron was labeled with a unique number. Neurons with GEMINI were isolated and displayed on the right. Scale bars: 50 μm.

**Extended Data Fig. 8.**
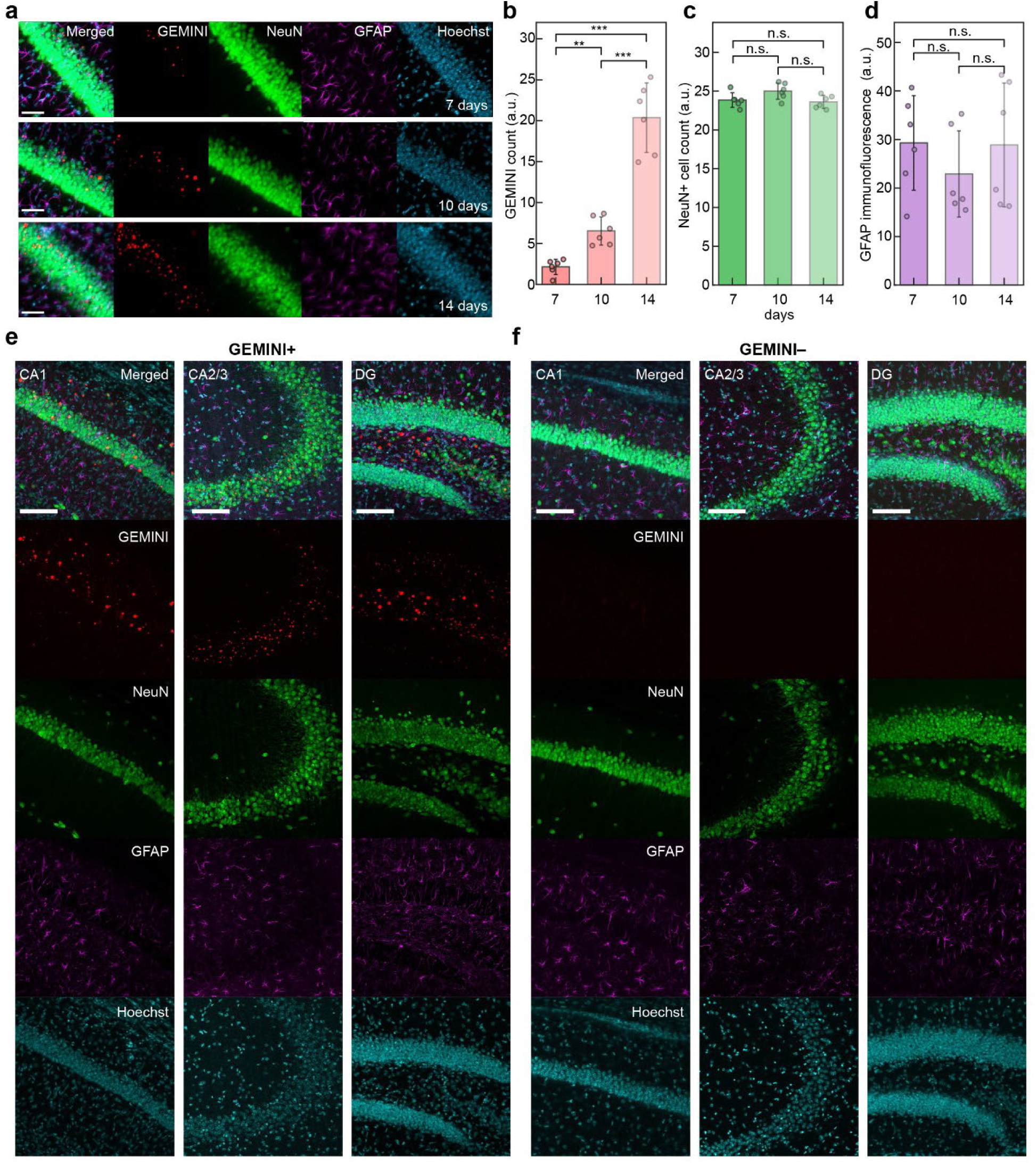
Immunohistochemical characterization of the brain tissue. **a,** Images showing the CA1 of the mouse hippocampus at 7-14 days after the injection of AAV encoding GEMINI. Scale bars: 50 μm. **b,** Comparison of GEMINI density across 7-14 days, where significant increase in GEMINI count was observed. **c,d** Comparison of the neuronal density (NeuN+ cells, **c**) and immune response (GFAP immunofluorescence, **d**) across 7-14 days after the injection of AAV encoding GEMINI. **e,f,** Images showing the CA1 (*left*), CA2/3 (*middle*), and DG (*right*) of the mouse hippocampus with (**e**) and without (**f**) intracellular GEMINI nucleation in neurons. The GEMINI+ group received intracranial injection of AAV encoding GEMINI, while the GEMINI– group received equivolume saline. GEMINI particles (JF_669_, red), neurons (anti-NeuN, green), astrocytes (anti-GFAP, magenta), and Nuclei (Hoechst, cyan), were stained and imaged. Scale bars: 100 μm. **b-d,** bars: mean; whiskers: s.d..

**Extended Data Fig. 9.**
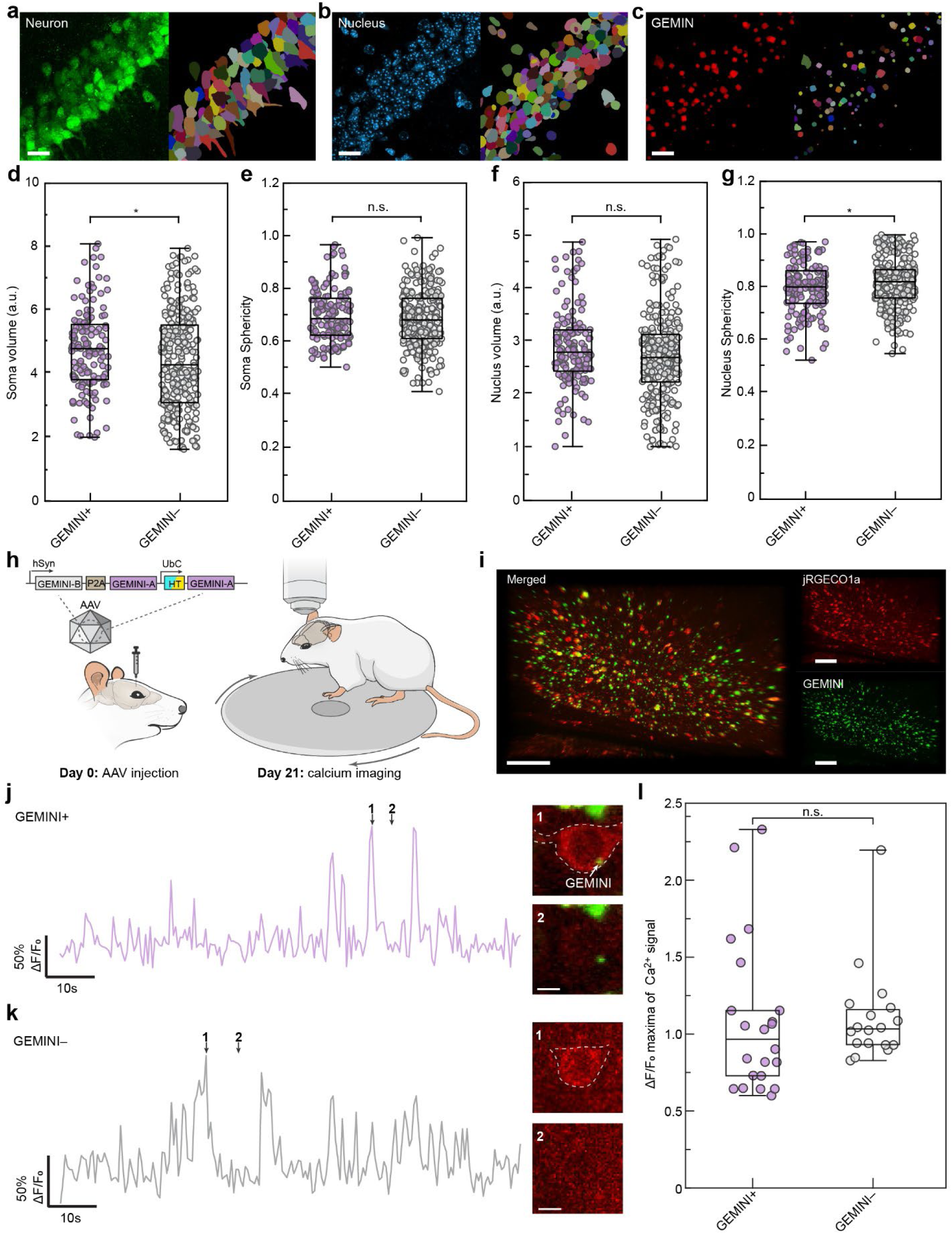
Impact of GEMINI growth on neuronal morphology and firing. **a-c,** Images (left) and their segmentations (right) of neurons (**a**), their nuclei (**b**), and GEMINI (**c**) within the mouse hippocampus CA1. The tissue was harvested 14 days after AAV injection. Scale bars: 20 μm. **d,e,** Comparison of the volume (**d**) and sphericity (**e**) of soma between GEMINI+ and GEMINI– neurons. **f,g,** Comparison of the volume (**f**) and sphericity (**g**) of soma between GEMINI+ and GEMINI– neurons. **h,** Schematic of the experimental design for in vivo characterization of neuronal activity. AAV encoding GEMINI was injected into the primary motor cortex (M1) of Thy1-jRGECO1a-WPRE transgenic mice on day 0, and two-photon calcium imaging was performed on day 21 with mice head-fixed on a running wheel. **i,** Image of the mouse’s cortical area showing the high-level co-expression of jRGECO1a and GEMINI. Scale bars: 50 μm. **j,k,** Calcium traces (left) from GEMINI+ (top) and GEMINI– (bottom) neurons in M1, with corresponding cellular images at indicated time points shown (right). Scale bars: 5 μm. **l,** Comparison of ΔF/F_0_ of Ca^2+^ peak maxima between GEMINI+ and – groups. (n=22/18 neurons for GEMINI+/– groups). **d-g,l,** Box bounds: 25th and 75th percentile; whiskers: minimum and maximum; squares: mean; and center lines: median.

**Extended Data Fig. 10.**
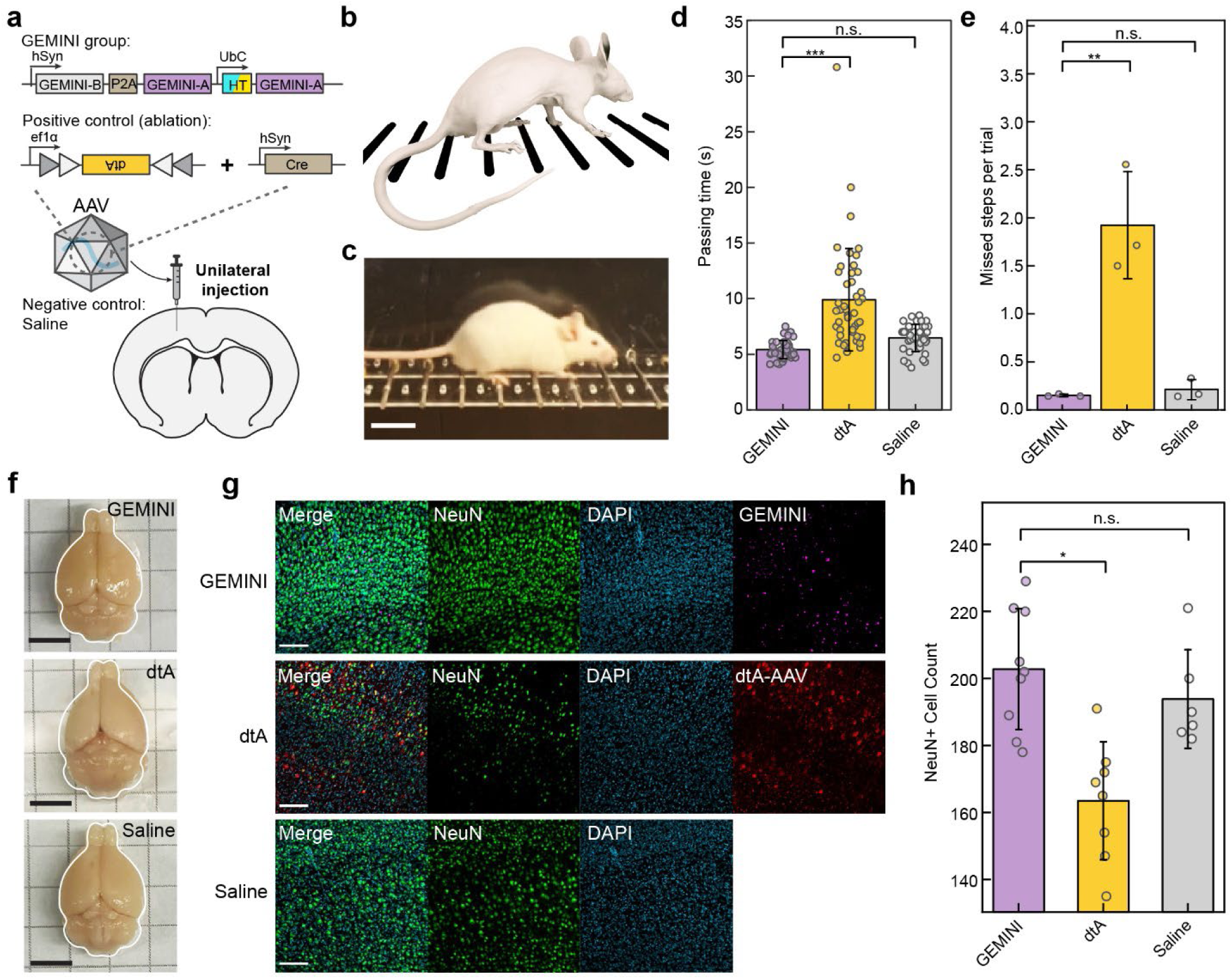
Impact of GEMINI expression on fine motor coordination. **a,** Schematic of experimental design. AAV encoding GEMINI was unilaterally injected into the primary visual cortex (V1) of mice (GEMINI group). A positive control group received AAVs expressing diphtheria toxin subunit A (dtA) at a comparable dose (dtA group), while a negative control group was injected with equivolume saline (saline group). **b,c,** Schematic (**b**) and snapshot (**c**) of the horizontal ladder (HL) rung walking test. The test was conducted 14 days after the AAV injection. Scale bar: 2 cm. **d,e,** Comparison of ladder crossing time (**d**) and mean missed steps per trial (**e**) across the GEMINI, dtA and saline groups. **f,** Images of the brains post-fixation. GEMINI and saline groups have comparable size of both hemispheres, while the dtA group exhibited visible shrinkage on the dtA-injected (*left*) hemisphere. Scale bars: 5 mm. **g,** Images of the M1 regions from GEMINI, dtA, and saline. Neurons (anti-NeuN, green), Nuclei (DAPI, cyan), GEMINI particles (JF_669_, violet), and cells transduced by the dtA AAV (mCherry, red) were stained and imaged. The mice were sacrificed immediately after the HL tests. Scale bars: 50 μm. **h,** Comparison of the neuronal density (NeuN+ cells) among the GEMINI, dtA, and saline groups (3 mice per group). **d,e,h,** bars: mean; whiskers: s.d..

