## Supplementary Information for "Genetically encoded assembly recorder temporally resolves cellular histories *in cellulo* and *in vivo*"

[1]Department of Materials Science and Engineering, Whiting School of Engineering, Johns Hopkins University, Baltimore, MD, USA. [2]Institute for NanoBiotechnology, Whiting School of Engineering, Johns Hopkins University, Baltimore, MD, USA. [3]Kavli Neuroscience Discovery Institute, Johns Hopkins University, Baltimore, MD, USA. [4]Institute for Protein Design, University of Washington, Seattle, WA, USA. [5]Division of Biology and Biological Engineering, California Institute of Technology, CA, USA. [6]Solomon H. Snyder Department of Neuroscience, School of Medicine, Johns Hopkins University, Baltimore, MD, USA. [7]Center for Cell Dynamics, School of Medicine, Johns Hopkins University, Baltimore, MD, USA. [†]Current address: Department of Biomedical Engineering, Southern University of Science and Technology, Shenzhen, China.

*These authors contributed equally to this work.

#

**Supplementary Movie Captions**

**Movie S1 | GEMINI growth in HEK293T cells.** Time-lapse video showing GEMINI growth in the HEK293T GEMINI clonal cell line.

**Movie S2 | Tracking the growth of single GEMINI particles.** Time-lapse videos with single-particle tracking (left) and the resolved growth profiles (right) of GEMINI particles.

### Supplementary Notes

#### Mathematical model of intracellular GEMINI growth

The model is built to describe the growth of GEMINI particles during the steady growth phase, when several assumptions can be applied:

(1) The transcription of the gene encoding GEMINI particles and the degradation of the mRNA reaches an equilibrium and remains stable across different phases of cell cycles, giving rise to a constant mRNA concentration in the cells.
(2) The translation kinetics of GEMINI mRNA is stable across different phases of the cell cycles, giving rise to a constant-rate synthesis of GEMINI building blocks.
(3) The cytoplasmic concentration of the building blocks remains constant (saturation condition) after GEMINI nucleation.
(4) The cytoplasmic building blocks are degraded at a constant rate across different phases of cell cycles.

As the rate of building-block synthesis is equal to the sum of the incorporation rate into GEMINI particles and the degradation rate, and both the synthesis and degradation are in constant rates, we can conclude that the building-block incorporation rate into GEMINI particles is also a constant.

Then, we simplify the GEMINI particles into spheres. We define $t_0 = 0$ when the radius $r = r_0$ . Over $dt$, the GEMINI particle grows by $dr$.

We have

$$\frac{4}{3}\pi(r_0 + dr)^3 - \frac{4}{3}\pi {r_0}^3 = Kdt$$

which can be simplified into

$$4\pi {r_0}^2 dr = Kdt$$

where $K$ is the rate constant for GEMINI growth.

By integrating both sides of the equation, we have

$$\int_{r_0}^{R} 4\pi {r_0}^2 dr = \int_0^t K dt$$

$$\frac{4}{3}\pi R^3 - \frac{4}{3}\pi {r_0}^3 = Kt$$

which can be rearranged into

$$R = ({r_0}^3 + \frac{3}{4\pi} Kt)^{1/3}$$

Let $\frac{3}{4\pi} K = K_0$ and ${r_0}^3 = A$, we have

$$R = (K_0\, t + A)^{1/3}$$

The equation can be used to solve the complete growth profile of individual GEMINI particles if 2 or more timestamps are applied, which will provide $(R_1, t_1)$ and $(R_2, t_2)$ to solve $K_0$ and $A$.

**Mathematical model of HTL diffusion into assemblies.**

We first assume that HaloTag-conjugated building blocks are homogeneously distributed in the entire assembly, which is a valid assumption based on our *in vitro* and *in vivo* results. For the *in vivo* timestamping, as the retro-orbitally injected HTL dyes were found to enter and exit the brain rapidly, on a timescale of minutes, we can further assume that a pulse of HTL dye arrives on the surface of an assembly in the brain at $t = 0$ without further supplement of HTL dye at $t > 0$. Over time, the finite dye molecules diffuse into the assembly and react irreversibly with the HaloTag. The assembly is simplified into a spherical shape.

In this mathematical model, we need to solve a three-dimensional reaction-diffusion problem. We first define several variables:

$\boldsymbol{C_{\mathrm{HT}}(r, t)}$**:** The concentration of HT-BBs without HTL binding in an assembly.

$\boldsymbol{C_{\mathrm{HT-L}}(r, t)}$**:** The concentration of HT-BBs with HTL binding in an assembly.

$\boldsymbol{C_{\mathrm{L}}(r, t)}$**:** The concentration of inward-diffusing HTL without bonding.

We will then derive the **time-dependent 3D reaction-diffusion equation.**

First of all, we will have **Mass Conservation:** the change in concentration $C_{\mathrm{L}}$ over time $t$ at a radial location can be expressed as:

$$\frac{\partial C_{\mathrm{L}}}{\partial t} = Net\ influx\ of\ HTL - Consumption\ of\ HTL\ by\ reaction = \frac{\partial C_{\mathrm{in}}}{\partial t} - \frac{\partial C_{\mathrm{con}}}{\partial t}$$

The **net influx of HTL** is governed by diffusion and therefore can be described by **Fick's second law** (spherical condition, simplified):

$$\frac{\partial C_{\mathrm{in}}}{\partial t} = D(\frac{\partial^2 C_{\mathrm{L}}}{\partial r^2} + \frac{2}{r}\frac{\partial C_{\mathrm{L}}}{\partial r})$$

where $D$ is the diffusion coefficient.

The **consumption of HTL by reaction** can be described by a first-order reaction, where the dye reacts with the HaloTag at a rate proportional to HTL concentration. The reaction rate is:

$$\frac{\partial C_{\mathrm{con}}}{\partial t} = kC_{\mathrm{L}}C_{\mathrm{HT}}$$

Therefore, the reaction-diffusion equation can be written as:

$$\frac{\partial C_{\mathrm{L}}(r,t)}{\partial t} = \left(\frac{\partial^2 C_{\mathrm{L}}(r,t)}{\partial r^2} + \frac{2}{r}\frac{\partial C_{\mathrm{L}}(r,t)}{\partial r}\right) - kC_{\mathrm{L}}(r,t)C_{\mathrm{HT}}(r,t)$$

The total concentration of HaloTag including those with and without HTL binding, is constant at all times and locations:

$$C_{\mathrm{HT}}(r,t) + C_{\mathrm{HT-L}}(r,t) = C_{\mathrm{tot}} = Const.$$

We will have the **initial boundary condition** as a pulse of HTL dye molecules at the assembly surface, represented by a surface delta function:

$$C_{\mathrm{L}}(r,0) = M\delta(r-R)$$

where $M$ is the total number of HTL ligand dye initially appear at the surface, $r$ is the radial distance from the center of the assembly, and $R$ is the radius of the assembly.

At the surface $r = R$, we also have the boundary condition:

$$-D\frac{\partial C_{\mathrm{L}}}{\partial r}|_{r=R} = \frac{dC_{\mathrm{L}}}{dt}|_{r=R} + kC_{\mathrm{L}}(R,t)C_{\mathrm{HT}}(R,t)$$

This partial differential equation (PDE) can be solved numerically, affording a plot shown in **Extended Data Fig. 6a**.

**Mathematical model of HTL binding with free cytoplasmic HaloTag.**

We first assume that the HaloTag-conjugated building blocks (HT-BBs) in the cells have reached a steady state, where their production, consumption, and cytoplasmic concentration are constant. The major route of consumption is their incorporation into the intracellular assembly. A pulse of HTL molecules will appear in the cytoplasm at $t = 0$ and be gradually consumed by irreversible binding with HaloTag. The HaloTag-HTL reaction follows first-order kinetics.

Three variables are included in this model:

$\boldsymbol{C_{\mathrm{HT}}(t)}$: The concentration of HT-BBs without HTL binding.

$\boldsymbol{C_{\mathrm{HT-L}}(t)}$: The concentration of HT-BBs with HTL binding.

$\boldsymbol{C_{\mathrm{L}}(t)}$: The concentration of cytoplasmic HTL.

We then define the following constants:

$\boldsymbol{R_P}$: Production rate of HT-BBs.

$\boldsymbol{R_C}$: Consumption rate of HT-BBs.

$\boldsymbol{k_b}$: Rate constant of HT-HTL binding.

$\boldsymbol{C_{\mathrm{L}}(0)}$: The cytoplasmic concentration of HTL at $t = 0$.

We have $\boldsymbol{R_P} = \boldsymbol{R_C} = \boldsymbol{R}$ at the steady state.

The changes in the concentrations can be written as:

**Rate of change of unbound HT-BB concentration:**

$$\frac{dC_{\mathrm{HT}}(t)}{dt} = R - R \cdot C_{\mathrm{HT}}(t) - k_b \cdot C_{\mathrm{HT}}(t) \cdot C_{\mathrm{L}}(t)$$

**Rate of change of HTL-bound HT-BB concentration:**

$$\frac{dC_{\mathrm{HT-L}}(t)}{dt} = k_b \cdot C_{\mathrm{HT}}(t) \cdot C_{\mathrm{L}}(t) - R \cdot C_{\mathrm{HT-L}}(t)$$

**Rate of change of HTL-bound HT-BB concentration:**

$$\frac{dC_{\mathrm{L}}(t)}{dt} = -k_b \cdot C_{\mathrm{HT}}(t) \cdot C_{\mathrm{L}}(t)$$

The differential equations can be solved numerically, affording the plot shown in **Extended Data Fig. 6b**.

As HT-BBs with and without HTL binding incorporate indiscriminately into the assembly, the cytoplasmic concentration of HTL-bound HT-BBs $\boldsymbol{C_{\mathrm{HT-L}}(t)}$ can well reflect the real-time incorporation of HTL-bound HT-BBs into the assembly and, thus, the change in HTL intensity along the radial axis as the assembly grows larger.

### Supplementary Methods

#### Cell Painting assay and segmentation

To prepare for the assay, cells were seeded onto gelatin-coated 12 mm #1 coverslips placed in 24-well plates. The following day, GEMINI expression was induced by adding 2 µg $mL^{-1}$ DOX. At the designated time point, subcellular structures were stained using the Image-iT™ Cell Painting Kit following the manufacturer's protocol (Invitrogen). Briefly, MitoTracker™ Deep Red was diluted in pre-warmed complete DMEM and incubated with the cells under standard culture conditions for 30 minutes. Cells were then fixed by adding an equal volume of 8% paraformaldehyde (PFA) in HBSS and incubated for 15 minutes at room temperature. After two washes with HBSS, cells were incubated for at least 30 minutes at room temperature in staining buffer (HBSS supplemented with 1% BSA and 0.1% Triton X-100) containing Hoechst 34580, Concanavalin A–Alexa Fluor™ 488, SYTO™ 14 Green, Wheat Germ Agglutinin–Alexa Fluor™ 555, and Alexa Fluor™ 568 Phalloidin. Confocal imaging was then performed using a Zeiss LSM780 equipped with aa spectral detector. For image analysis, the Hoechst channel was segmented using the generalized CellPose-SAM model[1] on the Hoechst channel; these masks served as seeds for watershed-based cell segmentation on the ConA-488 channel. SYTO14 and MitoTracker channels were segmented using adaptive Otsu and MaxEntropy thresholding.

#### Live/Dead Cell Viability Assay

Cell viability was assessed using the Calcein-AM/Ethidium Homodimer-1 (EthD-1) live/dead assay (Thermo), where Calcein-AM and EthD-1 stained live and dead cells, respectively. Hoechst staining was included for total cell segmentation. Cells were seeded in 24-well plates on gelatin-coated 12 mm #1 coverslips and allowed to adhere overnight. GEMINI expression was induced by adding DOX (2 µg $mL^{-1}$) at the designated time points. At designated times post-induction, cells were incubated with a staining solution containing 2 µM Calcein-AM, 4 µM EthD-1, and 5 µg $mL^{-1}$ Hoechst in HBSS for 30 minutes at 37 °C under standard culture conditions. Following staining, cells were washed once with HBSS and immediately imaged using an epifluorescence or confocal microscope. Calcein-AM+ cells (live) exhibited green fluorescence, EthD-1–positive cells (dead) showed red nuclear staining, and Hoechst-labeled nuclei were used to segment all cells. Cells were segmented using the algorithm in the segmentation of Cell Painting dataset. The percentage of dead cells was calculated as the number of EthD-1+ nuclei divided by the total number of Hoechst+ nuclei.

#### Western blotting

Cells were initially collected and washed twice with cold PBS to completely remove the medium. Cell pellets were obtained by centrifugation at 300 × g and lysed on ice using radioimmunoprecipitation assay (RIPA) extraction buffer (Boston BioProducts, BP-115), supplemented with phosphatase inhibitor (Thermo, J63907-AA), for 15 minutes. Lysis was followed by sonication for 15 seconds. The lysate was then centrifuged at 14,000 × g for 15 minutes at 4°C, and the supernatant was collected. Protein concentration was quantified using the BCA protein assay (Thermo, 23227), and 40 µg of protein was loaded for gel electrophoresis. Total protein was separated on a 10% SDS-polyacrylamide gel and transferred to a polyvinylidene difluoride (PVDF) membrane (Bio-Rad, 1620177). Blocking was performed with 5% milk in Tris Buffered Saline with Tween 20 (TBST) at room temperature for 1 hour with gentle shaking. The membrane was then incubated overnight at 4°C with primary antibodies (1:500). Afterward, the membrane was incubated with secondary antibodies for 1 hour at room temperature. Chemiluminescence was detected using a Gel Doc Imager (Bio-Rad) and High-sig ECL Western Blotting Substrate (Tanon, 180-501). Following imaging of the phosphorylated marker, the same membrane was reprobed for the unphosphorylated marker and the loading control after incubation in stripping buffer (Thermo, 46430) at room temperature for 7–10 minutes. The procedure was repeated as described above. Between each step, the membrane was washed three times with TBST

The following primary antibodies were used: anti-Phospho-IκBα antibody (Cell Signaling, 9246S, 1:1000), anti-IκBα antibody (Cell Signaling, 4812S, 1:1000), anti-Vinculin (Cell Signaling, 13901, 1:2000).

The following secondary antibodies were used: HRP-conjugated Goat Anti-Rabbit IgG(H+L) (ProteinTech, SA00001-2, 1:10000), Goat Anti-Mouse IgG (H + L)-HRP Conjugate (Bio-Rad, 1706516, 1:5000).

**Neuron and vasculature segmentation**

For neuron segmentation, the neuron image was first pre-processed slice-wise. The histogram of each slice is matched to a central reference plane was used to homogenize intensity over depth, and Gaussian smoothing is applied to suppress the neurite signal. Background removal is then achieved via grayscale morphological operations[2]. After the volume was rescaled to isotropic voxels, a watershed-inspired Voronoi–Otsu labeling generated an initial 3D segmentation. Connected components with radius lower than 3 μm were discarded, yielding the final soma labels.

For vasculature segmentation, the vessel images were first standardized via z-scoring (subtract mean, divided by standard deviation) and clipped from 2 to 5 to suppress outlier intensities. A difference-of-Gaussians filter was then applied to enhance tubular structures and attenuate background noise. An initial mask of vessels was generated with Otsu thresholding. Small, disconnected components with volume under 1000 $\mu m^3$ were removed.

**Two-photon calcium imaging in head-fixed awake mice**

Neuronal calcium imaging was performed in *Thy1-jRGECO1a* transgenic mice (Jackson Laboratory 030526). On day 6 post-injection of AAV into the primary visual cortex (AP: -2.5, ML: ±2.5, DV: -0.2 to -1.0 (0.2 increments)), dexamethasone (VetOne, NDC#13985-037-02) was administered via drinking water (1 mg $kg^{-1}$) to lower inflammation. On day 7, animals were anesthetized with inhaled isoflurane (0.25–5%) and placed in a custom stereotaxic frame. After hair removal and local administration of lidocaine (1%, VetOne, NDC#13985-222-04), the skull over the right visual cortex was exposed, and connective tissue was carefully removed. The area was rinsed with sterile saline, and a custom-made metal head plate was affixed to the skull using cyanoacrylate (Krazy Glue) and dental cement (C&B Metabond, Parkell Inc.), exposing the injection site. A 3-mm diameter circular craniotomy was made using a high-speed dental drill. The cranial window was sealed with a custom #1 (0.17 mm) 3-mm diameter glass coverslip using Vetbond™, and both the coverslip and head plate were further secured with cyanoacrylate and dental cement. Three hours after implantation, the awake mouse was head-fixed via the metal plate onto a custom stage mounted on a spinning platter. Calcium imaging was performed using a custom-built Bergamo II multiphoton microscope (Thorlabs) equipped with a Discovery NX laser (920 nm for GFP, 1040 nm for jRGECO1a). Images were acquired at 100–150 µm below the dura using a 16× Nikon CFI LWD Plan Fluorite objective at 2 Hz. For analysis, Suite2p was used for local motion correction, with the GFP-GEMINI channel used for registration due to its high signal-to-noise ratio. Registered TIFFs were saved individually and later re-stacked in Zen Blue. Cells containing or lacking crystals were manually cropped and exported. Calcium traces (from the soma, excluding crystals) were extracted manually using napari-time_series_plotter for analysis.

**Horizontal ladder rung walking test**

Following unilateral AAV (or sterile saline for negative control) injection into the primary motor cortex, mice were habituated to a horizontal ladder apparatus (Maze Engineers; 60 cm long, 10 cm wide, 2 cm rung spacing). On day 14 post-injection, locomotor behavior was assessed by recording a minimum of 10 complete runs per mouse. Trials in which mice reversed direction or failed to reach the end of the ladder were excluded from analysis. For each valid trial, two metrics were quantified: (1) the time required to traverse the ladder, excluding periods of immobility, and (2) the number of paw placement errors (defined as slips or misses). Immediately following behavioral testing, mice were transcardially perfused. Brains were collected and processed for immunohistochemistry using an anti-NeuN antibody to assess neuronal density. A secondary antibody conjugated to Alexa Fluor 647 was used to avoid spectral overlap with mCherry-tagged diphtheria toxin A (dtA) expression.

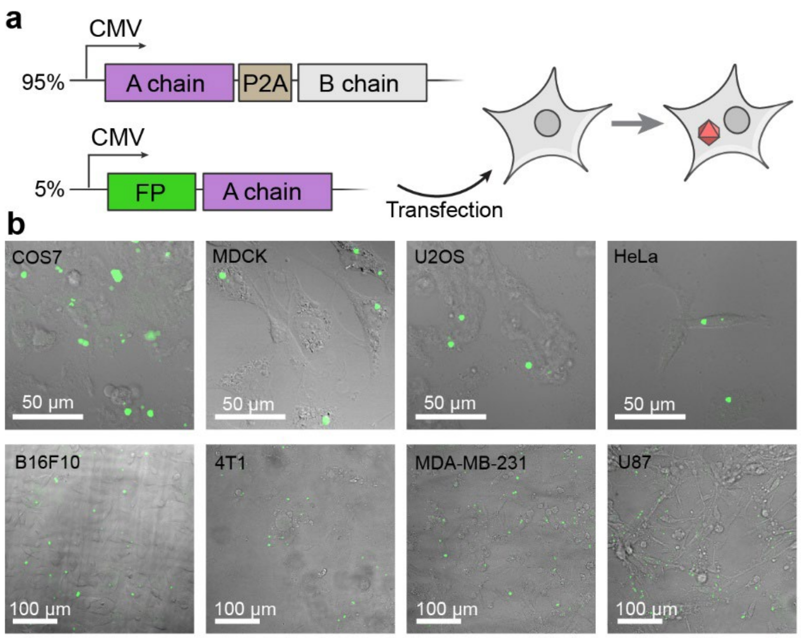


**Fig. S1| Plasmid design and transient transfection. a,** The design of plasmids for the screening of protein assemblies in live cells. The first plasmid has A and B chains of an assembly connected by a self-cleaving P2A peptide to afford their equimolar expression. The second plasmid has a fluorescent protein fused to the N terminus of the A chain. Both genes are driven by the constitutive CMV promoter. They are co-transfected at a 95%/5% ratio to live mammalian cells to obtain fluorescent intracellular assemblies. **b,** Expression of the final GEMINI scaffold (Lattice #1-v2) in various mammalian cell types.

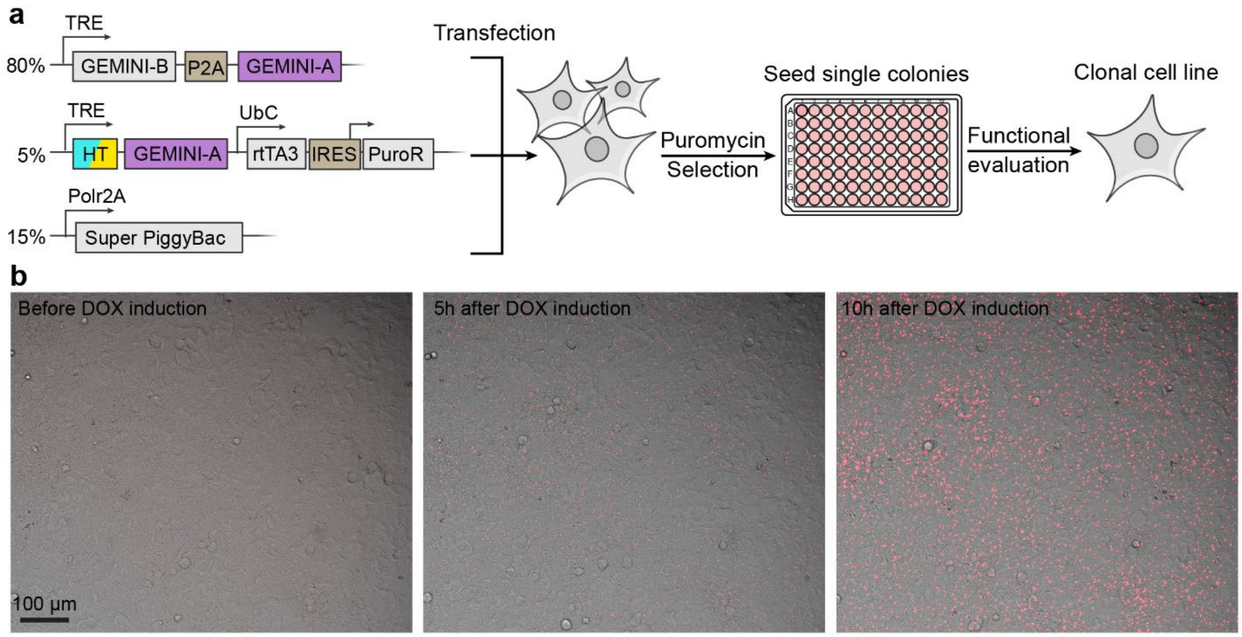


**Fig. S2|Development of the clonal GEMINI HEK cell line. a,** Procedures of clonal cell line development, where the genes of interest are integrated into the genome via the Super PiggyBac transposition system and the doxycycline-mediated expression was exploited for inducible GEMINI expression. The three plasmids are co-transfected to HEK293T cells, followed by puromycin selection, single-colony seeding, colony expansion, and functional evaluation to yield the clonal cell line. **b,** Images showing the intracellular GEMINI expression at various times after the induction of GEMINI expression. GEMINI particles appeared synchronously at *ca.* 5 hours post-induction and continued to grow over time.

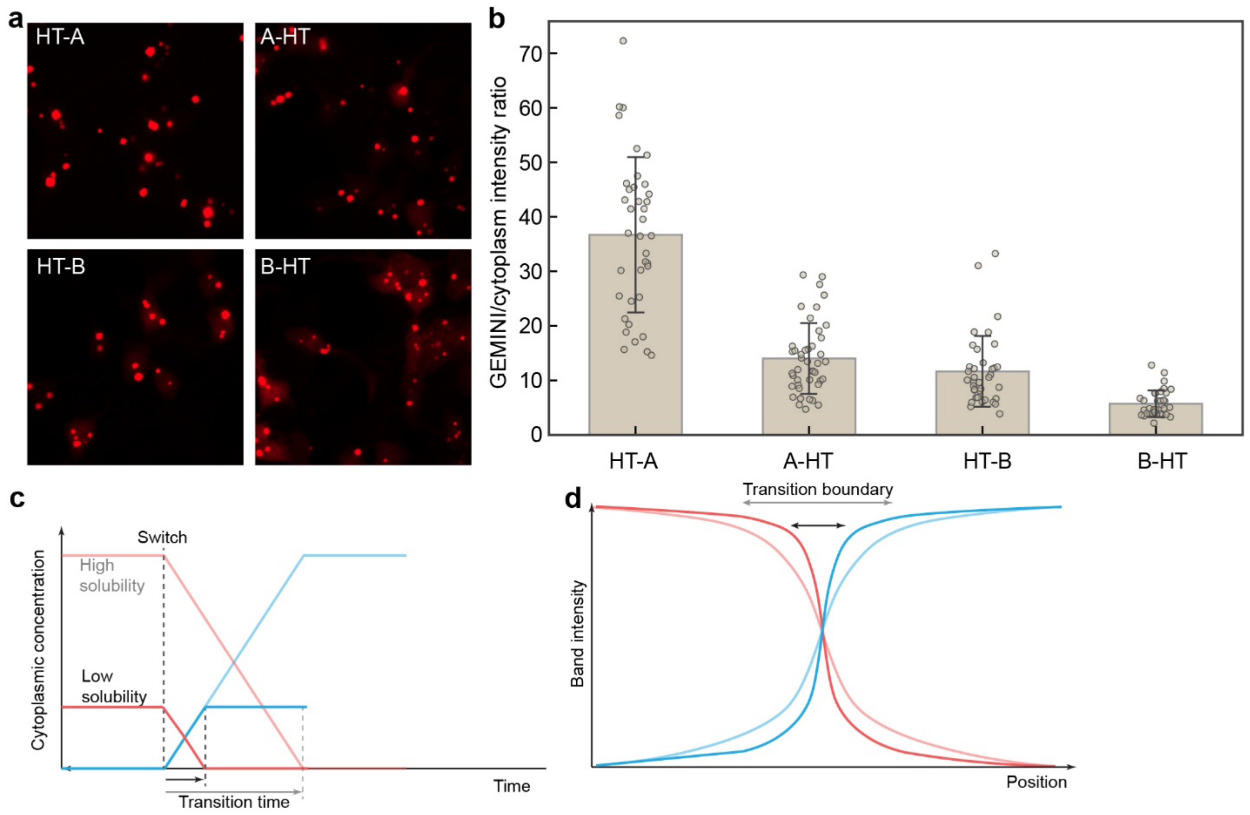


**Fig. S3|Capability of various termini in accommodating protein fusion. a,** Images showing the growth of GEMINI in HEK293T cells with HT fused to various termini of A and B chains. **b,** Comparison of the GEMINI-to-cytoplasm fluorescent intensity ratio among the different fusions. A higher GEMINI-to-cytoplasm intensity ratio indicates the more facile incorporation of the fusion into GEMINI. Fusing HT to the N terminus of the A chain exhibited the highest ratio and was therefore employed in this study to construct the timestamp and signal components. **b,** whiskers: standard deviation. **c,d,** Plots showing how the cytoplasmic concentration determines the band sharpness. A lower solubility results in a shorter transition time after switching to the second band (**c**), therefore affording a sharper transition boundary (**d**).

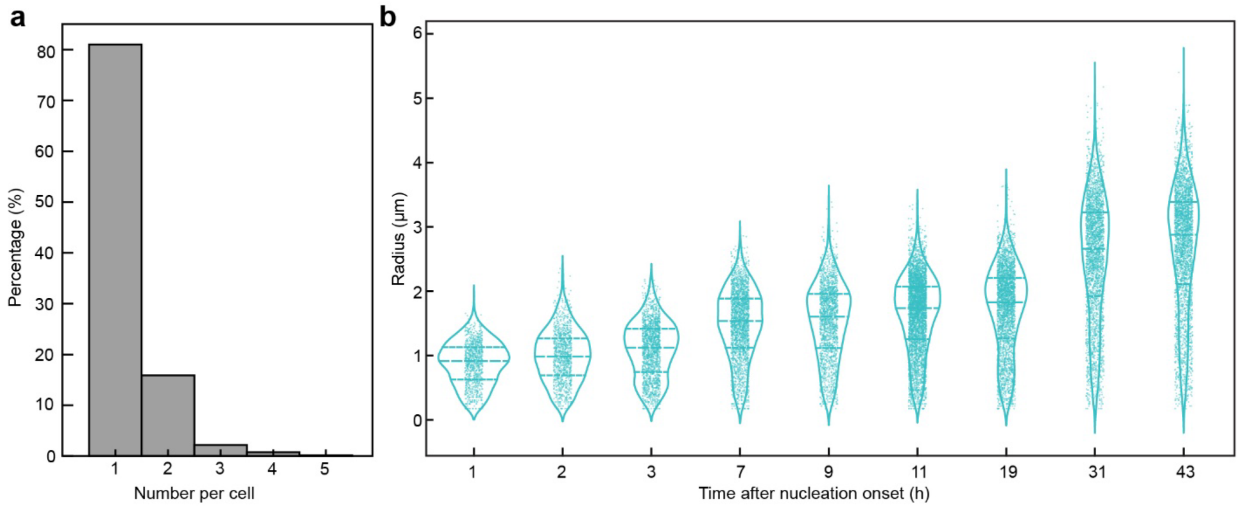


**Fig. S4|GEMINI nucleation and growth behaviors. a,** Analysis of the number of GEMINI nuclei per cell. Most cells only nucleate one GEMINI particle, a favorable property for the intracellular recording application. **b,** Statistic analysis of the GEMINI particle size at various times after GEMINI expression. The trend indicates the continuous growth of GEMINI particles, coinciding with our single-particle tracking results (**Movie S2**).

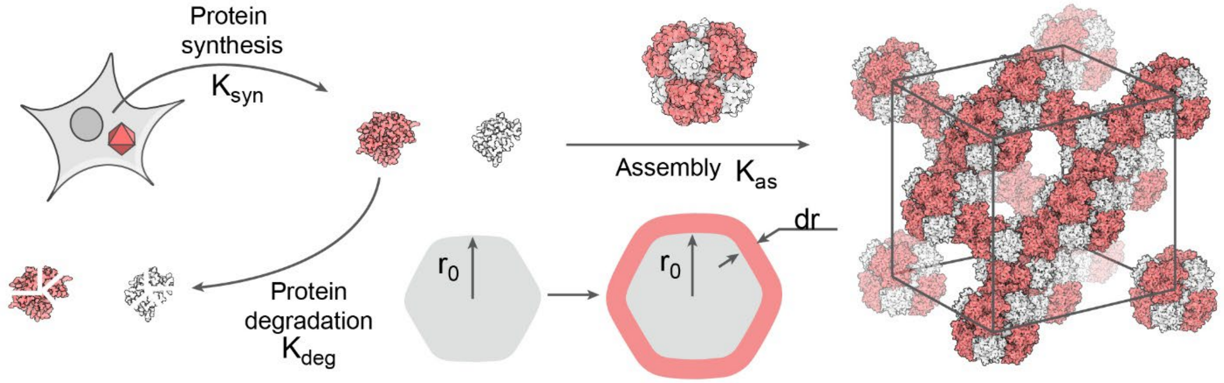


**Fig. S5| Model of intracellular GEMINI growth.** Schematic illustration of the intracellular expression of GEMINI building blocks and their assembly. The synthesized building blocks could either be incorporated into the GEMINI particle ($K_{as}$) or degraded ($K_{deg}$). The building blocks already in GEMINI particles are considered stable and protected from further degradation.

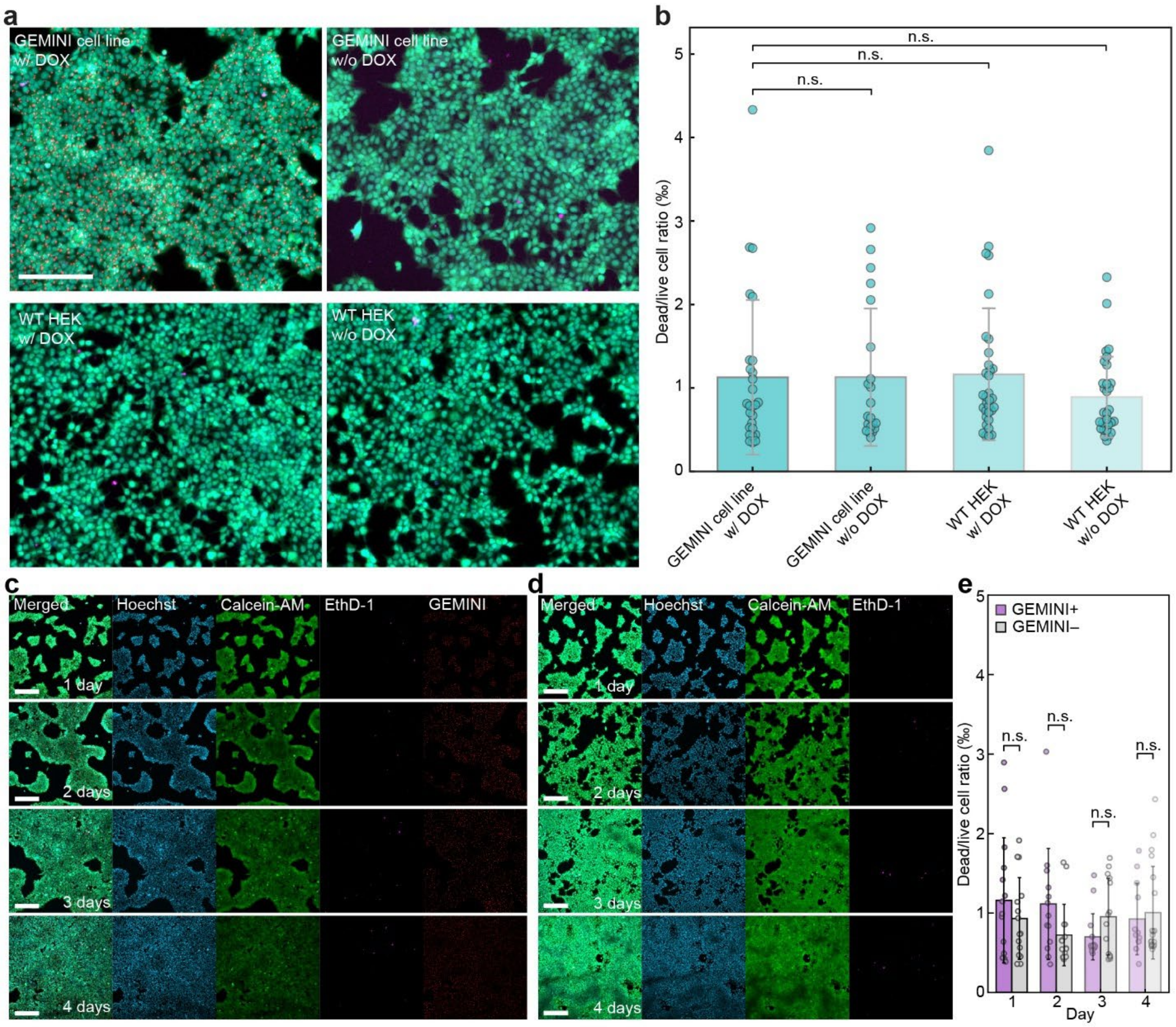


**Fig. S6| The impact of GEMINI on cell viability. a,** Images of cell live/dead assays on GEMINI HEK cell line with and without DOX, and wild-type (WT) HEK cells with and without DOX. The assays were performed at 48 hours post DOX induction. Green: live cells; Magenta: dead cells; Red: GEMINI. Scale bar: 200 µm. **b,** Comparison of the dead/live cell ratio among the four groups. No significant difference was found between GEMINI growing cells with the other three control group. **c,d,** Images of cell live/dead assays on GEMINI HEK cell line with (**c**) and without (**d**) DOX at 1-4 days after induction. Scale bars: 200 µm. **e,** Comparison of the dead/live cell ratio between the GEMINI+ (DOX+) and GEMINI– (DOX–) groups on different days. No significant difference was found between the two group in each day. **b,e,** bars: mean; whiskers: standard deviation.

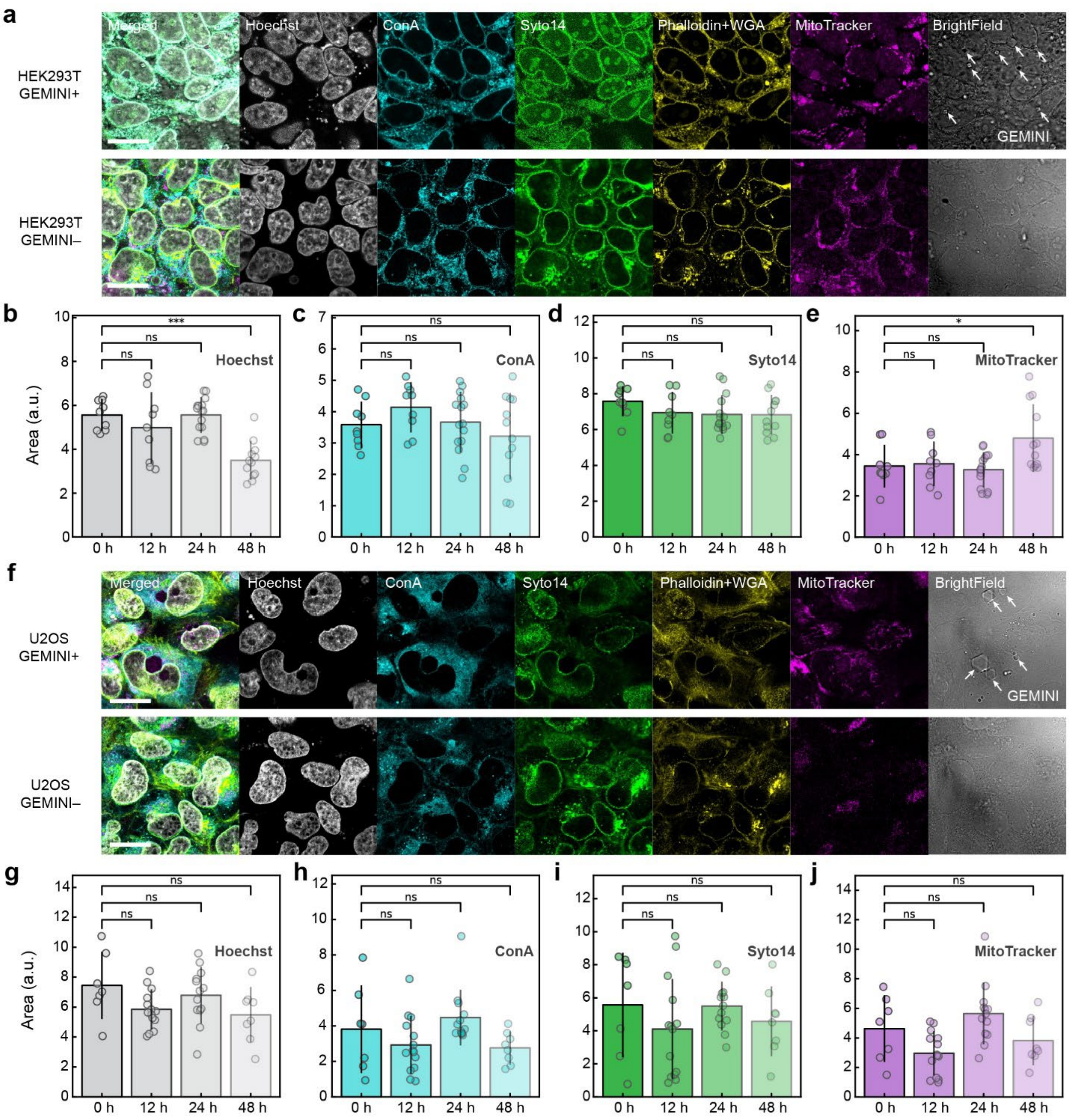


**Fig. S7| Impact of GEMINI growth on the morphology of cells and subcellular structures. a,** Images of the Cell Painting assay on HEK293T cells with (*top*) and without (*bottom*) the induction of cytoplasmic GEMINI growth by DOX. **b-e,** Statistical comparison of the segmented area of Hoechst (**b**), ConA (**c**), Syto14 (**d**), and MitoTracker (**e**) at 0-48 hours after DOX induction. **f,** Images of the Cell Painting assay on U2OS cells with (*top*) and without (*bottom*) the induction of cytoplasmic GEMINI growth by DOX. Bright field images were shown to highlight the GEMINI particles. Scale bars: 20 μm. **g-j,** Statistical comparison of the segmented area of Hoechst (**g**), ConA (**h**), Syto14 (**i**), and MitoTracker (**j**) at 0-48 hours after DOX induction. Hoechst: nucleus; Concanavalin A (ConA): endoplasmic reticulum; Syto14: nucleoli, cytoplasmic RNA; Phalloidin: F-actin; Wheat

Germ Agglutinin (WGA): Golgi, plasma membrane; MitoTracker: mitochondria. Phalloidin (Alexa Fluor 568) and WGA (Alexa Fluor 555) are combined in one channel as they are not separable on the microscope. GEMINI particles were highlighted in the bright-field images. Scale bars: 20 µm. **b-e,g-j,** bars: mean; whiskers: standard deviation. n.s.: no significance; *: $p<0.05$; ***: $p<0.001$.

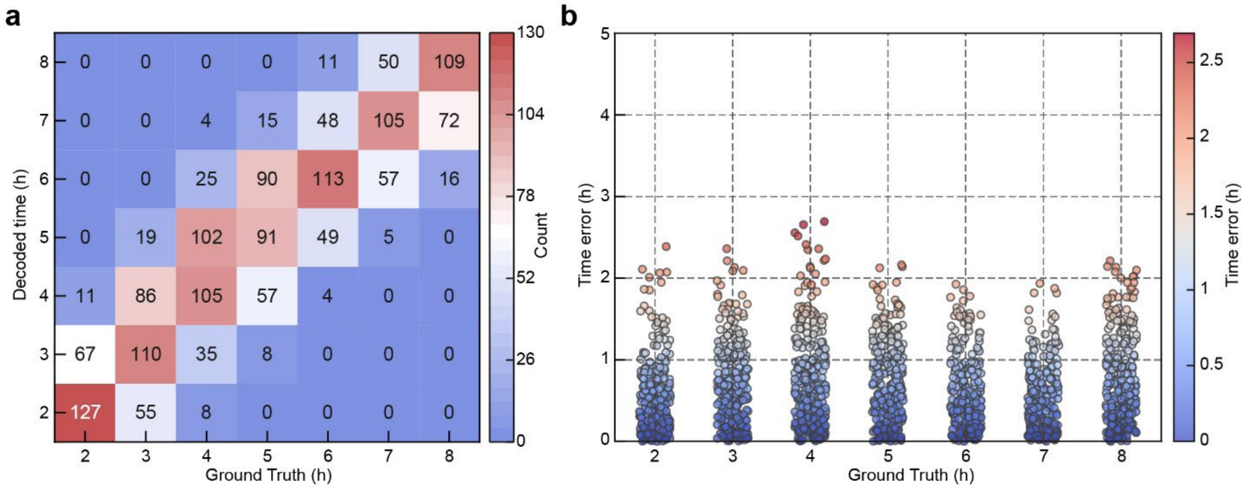


**Fig. S8| Evaluation of GEMINI temporal decoding accuracy. a,** Confusion matrix comparing the decoded time (y-axis) to the ground truth (x-axis) for GEMINI particles in **Fig. 2d**. The color scale indicates the number of particles decoded to each time, highlighting accurate versus misassigned time points. **b,** Time error distribution for individual GEMINI particles in **Fig. 2d** plotted against their ground truths (x-axis). Each dot represents a single particle, with color indicating the magnitude of time decoding error (color bar, *right*).

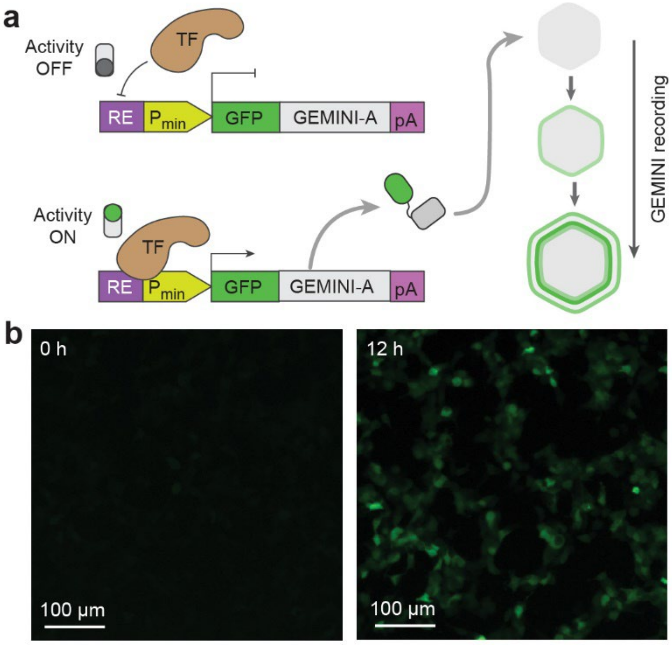


**Fig. S9| Signal transduction mechanism for the recording of NFκB signaling. a,** Schematic illustration of the signal transduction mechanism, where a promoter consists of the NFκB-response element and a minimal promoter is utilized to drive the expression of the GFP-fused A chain of GEMINI. **b,** Images showing the cytoplasmic intensity of the GFP signal at 0 h and 12 h after the induction of NFκB signaling by 10 ng $mL^{-1}$ TNFα.

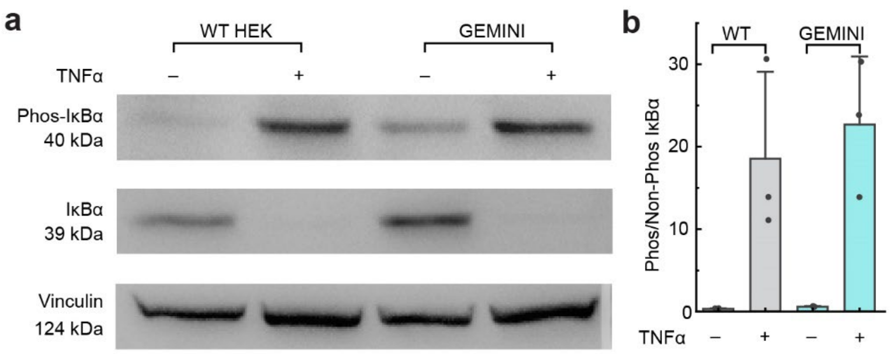


**Fig. S10 |Impact of GEMINI growth on NFκB signaling. a,** Western blots comparing the levels of phosphorylated IκBα, unphosphorylated IκBα, and the loading control (Vinculin) between WT HEK cells and the GEMINI clonal cell line, with and without incubation in 10ng $mL^{-1}$ TNFα. **b,** Statistical analysis of the ratio between phosphorylated and unphosphorylated IκBα. Similar responses to TNFα incubation were observed between WT HEK cells and the GEMINI clonal cell line. **b,** bars: mean; whiskers: standard deviation.

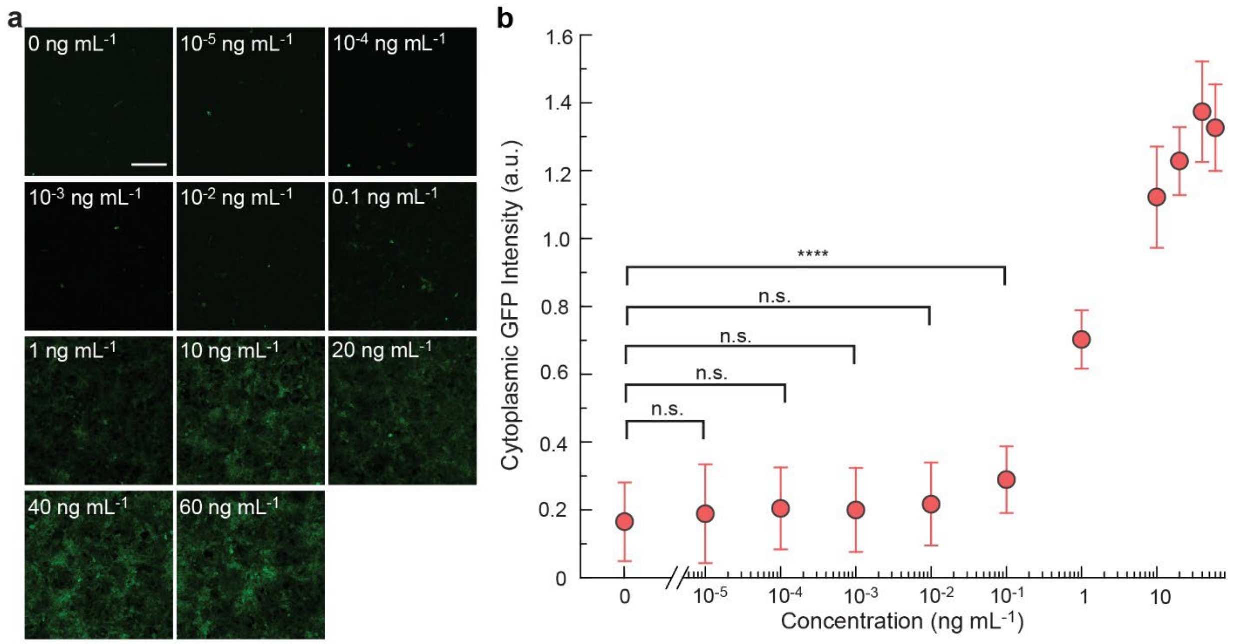


**Fig. S11| The response of a cytoplasmic NFκB reporter to various doses. a,** Images showing the cytoplasmic GFP fluorescence at various concentrations of TNF-α. Scale bar: 100 μm. **b.** Dose-response curve showing the intensity of the cytoplasmic GFP reporter at various concentrations of TNF-α. A concentration of $10^{-2}$ ng $mL^{-1}$ and lower cannot be detected by the cytoplasmic reporter. A *ca.* 700% dynamic range was obtained. **b,** circles: mean; whiskers: standard deviation.

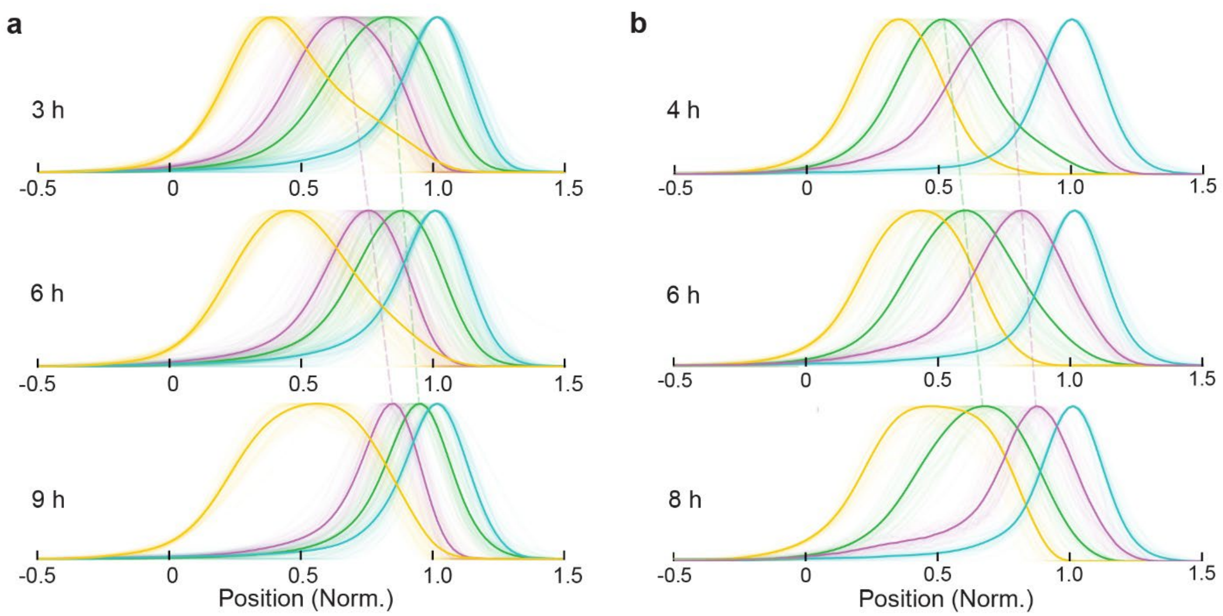


**Fig. S12| Mean fluorescence profiles in NFκB recording. a,** Mean fluorescence profiles after normalization corresponding to the recording of NFκB activation in **Fig. 3a-c**. The onsets of NFκB activation signals (green) and the accompanying timestamp (violet) shifted to the right with respect to a later activation time. b, Mean fluorescence profiles after normalization corresponding to the recording of NFκB downregulation in **Fig. 3d-f**. The peaks of NFκB signals (green) and the accompanying timestamp (violet) shifted to the right with respect to a later TNFα removal.

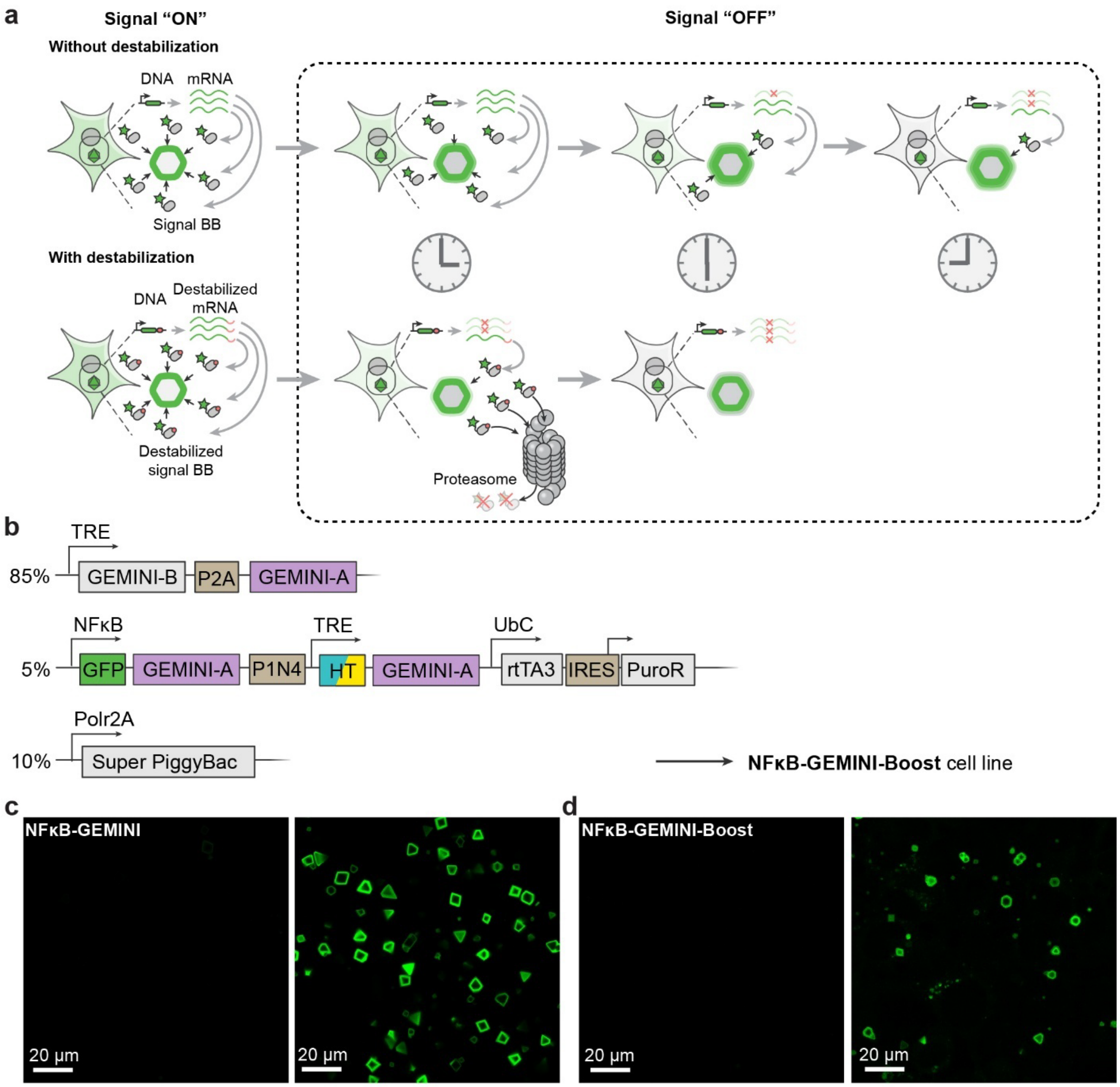


**Fig. S13| Clonal cell line for recording fast NFκB dynamics. a,** Schematic illustration of the impact of mRNA and protein stability on resolving the temporal dynamics of cellular events. Destabilization of both mRNA and protein of signal-BB accelerates their removal from the cytoplasm when the signal is turned off, resulting in a sharper decay of the “OFF” signal in GEMINI particles. **b,** DNA constructs used in the development of NFκB-GEMINI-Boost cell lines. NFκB-GEMINI-Boost contains a destabilization domain P1N4 that expedites the turnover of mRNA and protein. **c,** Images of GEMINI particles grown in the NFκB-GEMINI cells without (left) and with (right) incubation in 10 ng mL-1 TNF-α. **d,** Images of GEMINI particles grown in the NFκB-GEMINI-Boost cells without (left) and with (right) incubation in 10 ng mL-1 TNF-α. TNF-α was added 24 hours after DOX induction. Scale bars: 20 μm.

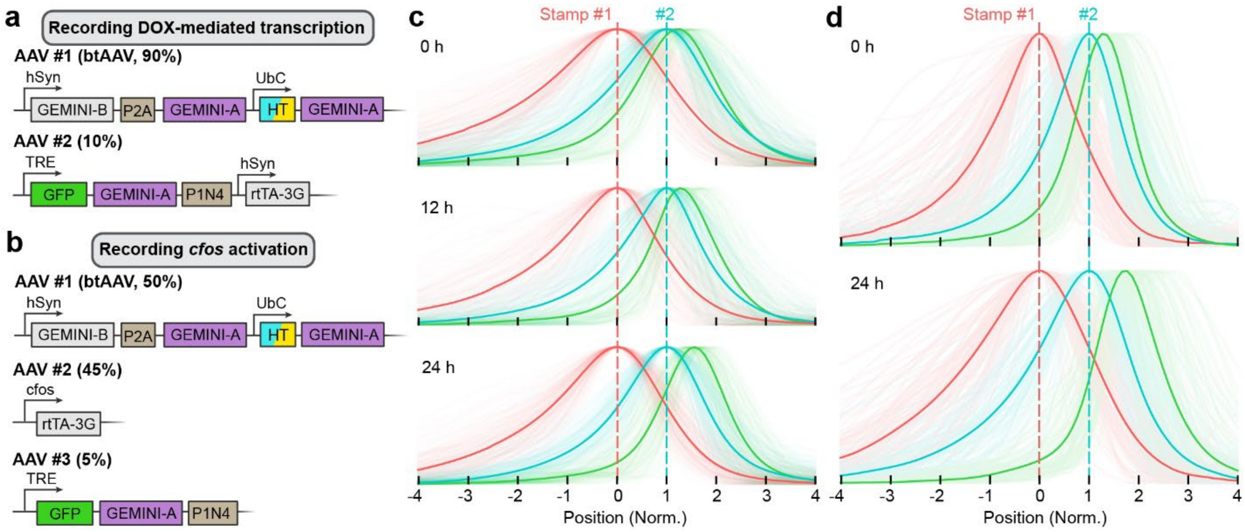


**Fig. S14| *In vivo* GEMINI recording. a,** DNA constructs and the ratio of AAV doses for *in vivo* recording of DOX-mediated transcription, corresponding to the experiments in **Fig. 5d-f**. **b,** DNA constructs and the ratio of AAV doses for *in vivo* recording of seizure-induced *cfos* activation, corresponding to the experiments in **Fig. 5g-i**. **c,** Mean fluorescence profiles of GEMINI recording of DOX-mediated transcription, with DOX administered at 0 h (top), 12 h (middle), and 24 h (bottom). **c,** Mean fluorescence profiles of GEMINI recording of activity-dependent *cfos* activation, with seizure being induced at 0 h (top) and 24 h (bottom).

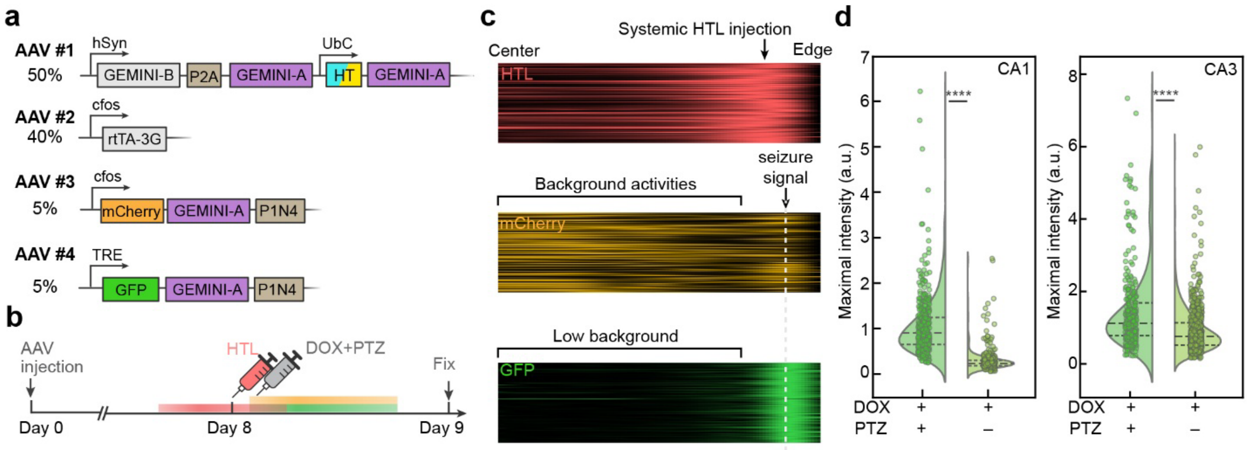


**Fig. S15| *In vivo* transduction of *cfos* activity via direct and DOX-mediated transcription. a,** Schematic of the DNA constructs used in the AAVs co-injected into the mouse hippocampus for comparing two methods of signal transduction, where signal BBs were expressed under the direct control of the cfos promoter (mCherry-GEMINI-A, AAV #3) and via the Tet-ON system (GFP-GEMINI-A, AAV #2 and #4). **b,** Experimental design showing that HTL was injected retro-orbitally on day 8 post-AAV injection. DOX and PTZ were administered intraperitoneally approximately 30 minutes later. **c,** Normalized fluorescence profiles of HTL (stamp, red), mCherry (direct *cfos* signal, yellow), and GFP (DOX-gated *cfos* signal, green) along the radial axis of individual GEMINI particles. Each row represents one particle, with the profiles normalized to the particle centers (left edge) and HTL peaks. Both transcription methods captured *cfos* activation, though background activity was more prominent without Tet-ON gating (yellow). **d,** Statistical analysis of maximal fluorescence intensity in GEMINI particles with and without PTZ administration. DOX was injected in all cases. Hippocampal regions CA1 (left) and CA3 (right) were analyzed separately. Dash lines: 25%, 50%, and 75%. ****: $p<0.0001$.

**Table S1| Plasmids cloned and/or used in this work.**

| Plaslmid ID | Description | Addgene ID |
|---|---|---|
| YY120: pPB_TRE::GEMINI(B)-p2A-GEMINI(A) | PiggyBac plasmid expressing the two GEMINI building blocks (A and B) in an equimolar ratio under the control of a doxycycline-inducible promotor. | 228881 |
| YY105: pPB_TRE::HaloTag-GEMINI(A)_UbC::rtTA3-IRES-PuroR | PiggyBac plasmid expressing the HaloTag-GEMINI(A) under the control of a doxycycline-inducible promotor, and rtTA3 and PuroR under the control of a UbC promoter. | 228882 |
| YY171: pPB_NFκB::GFP-GEMINI(A)_TRE::HaloTag-GEMINI(A)_UbC::rtTA3-IRES-PuroR | PiggyBac plasmid expressing the GFP-GEMINI(A) under the control of a synthetic NFκB-responsive promotor, and rtTA3 and PuroR under the control of a UbC promoter. | 228883 |
| JL054: pPB_NFκB::GFP-GEMINI(A)-P1N4_TRE::HaloTag-GEMINI(A)_UbC::rtTA3-IRES-PuroR | PiggyBac plasmid expressing the destabilized GFP-GEMINI(A) under the control of a synthetic NFκB-responsive promotor, and rtTA3 and PuroR under the control of a UbC promoter. | 228884 |
| YY203: pAAV_cfos::mCheery-GEMINI(A)-P1N4 | AAV plasmid expressing the destabilized mCherry-GEMINI(A) under the control of a cfos promotor. | 228885 |
| YY206: pAAV_hSyn::GEMINI(B)-p2A-GEMINI(A)_UbC::HaloTag-GEMINI(A) | AAV plasmid expressing the wo GEMINI building blocks (A and B) in an equimolar ratio under the control of a hSyn promoter, and HaloTag-GEMINI(A) under the control of a UbC promoter. | 228886 |
| pAAV-FAH-rtTA3G | AAV plasmid expressing rtTA3G under the control of a cfos promotor. | 120309 |
| YY225: pAAV_TRE::GFP-GEMINI(A)-P1N4 | AAV plasmid expressing the destabilized GFP-GEMINI(A) under the control of a doxycycline-inducible promotor. | 228888 |

| | | |
|---|---|---|
| YY229: pAAV_TRE::GFP-GEMINI(A)-P1N4_hSyn::rtTA3G | AAV plasmid expressing the destabilized GFP-GEMINI(A) under the control of a doxycycline-inducible promotor, and rtTA3G under the control of a hSyn promotor | 228889 |

**Table S2| Protein variants that assemble spontaneously *in cellulo*.**

| Variants | Amino acid sequence |
|---|---|
| Lattice #1_v2<br><br>(GEMINI) | A chain:<br>SKAKIGIVTVSDRASAGITADISGKAIILALNLYLTSEWEPIYQVIPDE<br>QKVIERTLIKMADIQDCCLIVTTGGTGPAKRDVTPEATEAVCDRMM<br>PGFGELMRAESLKEVPTAILSRQTAGLRGDSLIVNLPGDPASISDC<br>LLAVFPAIPYCIDLMEGPYLECNEAMIKPFRPKAK<br><br>B chain:<br>VRGIRGAITVNSDTPTSIIIATILLLEKMLEANGIQSYEELAAVIFTVTE<br>DLTSAFPAEAARQIGMHRVPLLSAREVPVPGSLPRVIRVLALWNTD<br>TPQDRVRHVYLSEAVRLRPDLESAQ |
| Lattice #2_v3 | A chain:<br>EEIVEKAERKLKFLLQEAEEGGKEDALEIAEKLAELAKEALRVLAEA<br>GGSPELMLRLMETAARALARIARLGGDELREEIKKIMAELVAKAISL<br>LIRMLKRSGSSYEEIAEAVAKAVAKIVEAAKESGMSEDEIAEIVARVI<br>SEVIRTLKESGSSAEVIAEIVARIVAEIVEALKRSGTSEDEIAEIVARVI<br>SEVIRTLKESGSSSILIALIVARIVAEIVEALKRSGTSEDEIAEIVARVIS<br>EVIRTLKESGSSYEIIALIVAMIVAEIVRALLRSGTSEEEIAKIVARVM<br>NEVLRTLRESGSDFEVIREILRLILAAIRAALQKGGVSEDEIMRIEIKI<br>LLMLLRLSTAELERATRSLKAITEELKKNPSEDALVEHNRAIVEHNR<br>IIVFNNILIALVLEAIVRAIK<br><br>B chain:<br>RSLREQEELAKRLMELLLKLLRLQMTGSSDEDVRRLMLRIIELVEEI<br>EELAREQK |
| Lattice #3_v1 | A chain:<br>ATMALAYVMLGLLLSLLNRLSLAAEAYKKAIELDPNDALAWLLLGSV<br>LEKLKRLDEAAEAYKKAIELKPNDASAWKELGKVLEKLGRLDEAAK<br>AYAEAIKLDPSDAEAAKELGKVLEKLGQLELAERAYQLAIELDPN<br><br>B chain:<br>KMEELFKKHKIVAVLRANSVEEAKEKALAVFRGGVHLIEITFTVPDA<br>DTVIKELSFLKEKGAIIGAGTVTSLEQCQKAVESGAEFIVSPHLDPEI<br>SKFCKINGVFYMPGVMTPTELVKAMKLGHTILKLFPGEVVGPQFV<br>KAMKGPFPNVKFVPTGGVNDQNVCEWFKAGVLAVGVGSALVKG<br>TPEQVEMLAVLFVAKIAGCTE |
| Lattice #3_v2 | A chain:<br><br>ATMALAYVMLGLLLSLLNRLSLAAEAYKKAIELDPNDALAWLLLGSV<br>LEKLKRLDEAAEAYKKAIELKPNDASAWKELGKVLEKLGRLDEAAK<br>AYAEAIKLDPSDAEAAKELGKVLEKLGQLELAERAYQLAIELDPN<br><br>B chain: |

| | |
|---|---|
| | KMEELFKKHKIVAVLRANSVEEAKEKALAVFRGGVHLIEITFTVPDA<br>DTVIKELSFLKEKGAIIGAGTVTDKRQCKKAVESGAEFIVSPHLDPE<br>ISEFCKMMGVFYMPGVMTPTELVKAMKLGHTILKLFPGEVVGPQF<br>VKAMKGPFPNVKFVPTGGVNDQNVCEWFKAGVLAVGVGSALVK<br>GTPEQVEMLAVLFVAKIAGCTE |
| Lattice #3_v3 | A chain:<br><br>ATMALAYVMLGLLLSLLNRLSLAAEAYKKAIELDPNDALAWLLLGSV<br>LEKLKRLDEAAEAYKKAIELKPNDASAWKELGKVLEKLGRLDEAAK<br>AYAEAIKLDPSDAEAAKELGKVLEKLGQLELAERAYQLAIELDPN<br><br>B chain:<br>MRGHHHHHHGSSKMEELFKKHKIVAVLRANSVEEAKEKALAVFR<br>GGVHLIEITFTVPDADTVIKELSFLKEKGAIIGAGTVTSLEQCKKAVE<br>SGAEFIVSPHLDPEISKFCKIEGVFYMPGVMTPTELVKAMKLGHTIL<br>KLFPGEVVGPQFVKAMKGPFPNVKFVPTGGVNDQNVCEWFKAG<br>VLAVGVGSALVKGTPEQVEMLAVLFVAKIAGCTE |
